# A molecular cell atlas of the human lung from single cell RNA sequencing

**DOI:** 10.1101/742320

**Authors:** Kyle J. Travaglini, Ahmad N. Nabhan, Lolita Penland, Rahul Sinha, Astrid Gillich, Rene V. Sit, Stephen Chang, Stephanie D. Conley, Yasuo Mori, Jun Seita, Gerald J. Berry, Joseph B. Shrager, Ross J. Metzger, Christin S. Kuo, Norma Neff, Irving L. Weissman, Stephen R. Quake, Mark A. Krasnow

## Abstract

Although single cell RNA sequencing studies have begun providing compendia of cell expression profiles, it has proven more difficult to systematically identify and localize all molecular cell types in individual organs to create a full molecular cell atlas. Here we describe droplet- and plate-based single cell RNA sequencing applied to ∼75,000 human lung and blood cells, combined with a multi-pronged cell annotation approach, which have allowed us to define the gene expression profiles and anatomical locations of 58 cell populations in the human lung, including 41 of 45 previously known cell types or subtypes and 14 new ones. This comprehensive molecular atlas elucidates the biochemical functions of lung cell types and the cell-selective transcription factors and optimal markers for making and monitoring them; defines the cell targets of circulating hormones and predicts local signaling interactions including sources and targets of chemokines in immune cell trafficking and expression changes on lung homing; and identifies the cell types directly affected by lung disease genes and respiratory viruses. Comparison to mouse identified 17 molecular types that appear to have been gained or lost during lung evolution and others whose expression profiles have been substantially altered, revealing extensive plasticity of cell types and cell-type-specific gene expression during organ evolution including expression switches between cell types. This atlas provides the molecular foundation for investigating how lung cell identities, functions, and interactions are achieved in development and tissue engineering and altered in disease and evolution.

## Introduction

Over the past two centuries, hundreds of human cell types have been discovered, categorized, and studied by microscopy, creating the classical atlases that provide the cellular foundation for modern medicine^1–5^. In the past several decades, cell-type-specific marker genes have been identified that supplement the histological descriptions and provide molecular definitions and functions of the cell types^6–9^. This reached its apex in the systematic profiling studies that elucidate genome-wide expression profiles of purified cell populations and, more recently, of individual cells by single cell RNA sequencing (scRNAseq)^10–14^.

Although initial scRNAseq studies focused on specific cell types, tissue compartments, and biological processes, large scale molecular cell atlases of organs and even whole organisms are now possible because of improvements in throughput and cost that allow expression profiling of thousands of individual cells, without cell purification or prior knowledge of cell identity to obtain expression profiles of at least the most abundant and easy to isolate cell types^15–27^. Perhaps the greatest current challenge is obtaining profiles of rare and fragile cell types, or even just assessing the “completeness” of a molecular cell atlas. Indeed, most scRNAseq efforts to date have left the identities and tissue locations of many transcriptionally-distinct cell populations (computationally-defined cell “clusters”) uncertain, obscuring their relationship to the classical, histologically-defined cell types and leaving open the possibility that some represent previously unrecognized cell types, subtypes or cell states. A complete molecular cell atlas could identify new cell types and new biochemical functions and interactions of known cell types, and would provide the molecular foundation for investigating how cells are specified and how they are altered in disease and evolution.

We set out to create a comprehensive molecular cell atlas of the adult human lung, both for its basic science value and potential clinical applications since pulmonary diseases and infections are among the leading causes of morbidity and mortality worldwide^28, 29^. This is a substantial technical challenge because the lung is comprised of 45 histologically-defined cell types with diverse structures and functions that vary in abundance over five orders of magnitude^30^ (Table S1). Here we describe a systematic single cell RNA sequencing approach toward capturing, annotating, and analyzing the expression profiles of all lung cell types. We do so by using fresh blood and lung tissue obtained intraoperatively, balancing the abundance across the major tissue compartments of the 75,000 cells analyzed, employing both broad cell capture (droplet-based) and deep expression (plate-based) scRNAseq profiling strategies, and using the distinguishing molecular features along with ascertained spatial information of the novel and ambiguous cell clusters to assign cellular identities. We identify 41 of 45 previously known lung cell types and subtypes including all but the exceedingly rare ones, plus 14 new ones. We show how this comprehensive lung cell atlas provides novel insights into the functions, regulation, and interactions of the known as well as the new cell types; into which cell types are targeted in disease and how each type can be specified in development and tissue engineering; and into organ evolution by revealing lung cell types that have been gained, lost, and altered from mouse to human. The atlas provides a benchmark for analysis of diseased lung tissue, and the approach can be readily applied to other organs to create a human molecular cell atlas.

## Results

### Single cell RNA sequencing of ∼75,000 cells identifies 58 human lung cell populations

To create a comprehensive atlas of the human lung, we set out to capture the full diversity of cell types in its four major tissue compartments (epithelial, endothelial, stromal, immune), and across the proximal-to-distal axis of the lung. We acquired histologically normal lung tissue samples intraoperatively from bronchi (proximal), bronchiole (medial), and alveolar (distal) regions (Fig. S1) along with samples of peripheral blood and immediately began processing (Fig. 1a). Lung samples were independently dissociated into single cell suspensions, and each lung cell suspension was then separated into epithelial (EPCAM^+^), endothelial/immune (CD31^+^/CD45^+^) and stromal (EPCAM^-^, CD31^-^/CD45^-^) populations by fluorescence-activated cell sorting (FACS) or magnetic-assisted cell sorting (MACS) (Fig. S2a). This allowed us to balance compartmental representation for sequencing, which for lung is otherwise dominated by immune and endothelial cells. Index sorting further provided surface antigen levels and other cell parameters for the FACS-sorted cells. Some of the blood cells were also index sorted by FACS to balance representation of the major immune cell lineages (Figs. 1a, S2b). Sequencing libraries were prepared from MACS-sorted lung cell populations and unsorted blood cells using droplet-based 10x Chromium (10x) protocol, or from FACS-sorted individual lung and blood cells using SmartSeq2 (SS2) protocol^31^. The higher cell throughput and lower cost of 10x enabled discovery of rare cell types, whereas SS2 gave deeper transcriptomic information that aided cell classification and detection of genes expressed at low levels such as transcription factor and receptor genes. There were also platform-specific idiosyncrasies; for example alveolar macrophages were the most abundant cell type detected in 10x samples but rare in SS2 samples, whereas neutrophils were represented only in SS2 samples. We sequenced thousands of cells from each compartment for each subject (Table S2) to directly compare as many cell types as possible without batch correction, and we did so for three subjects (two males, one female) to address individual differences (see below). High quality transcriptomes were obtained from ∼75,000 cells, 65,662 using 10x and 9,404 using SS2 (Fig. 1a).

**Figure 1.**
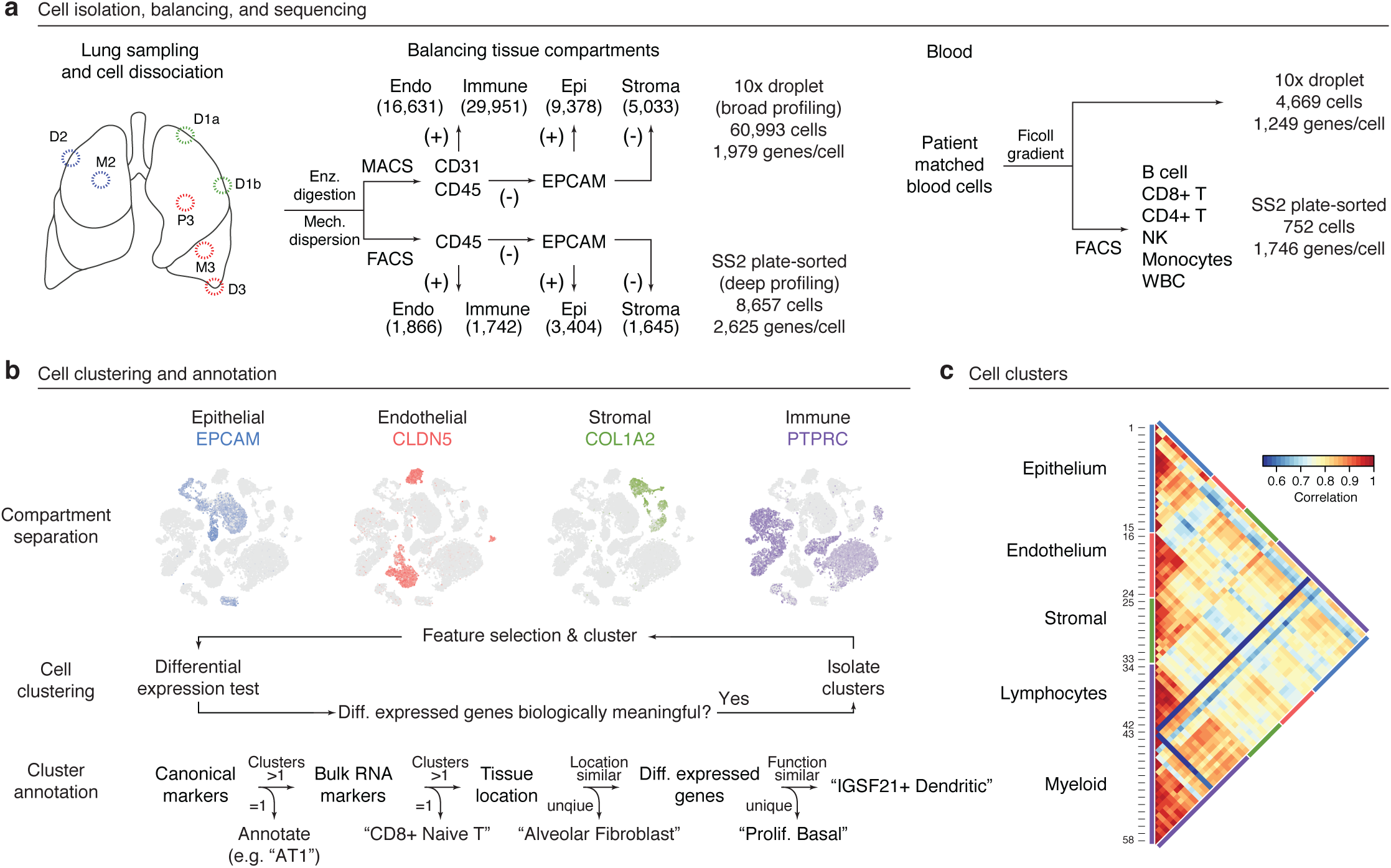
Strategy for single cell RNA sequencing and annotation of human lung and blood cells. **a**, Workflow for capture and mRNA sequencing of single cells from the healthy unaffected regions indicated (D, distal; M, medial; P, proximal lung tissue) of fresh, surgically resected lungs with focal tumors from three subjects (1, 2, 3) and their matched peripheral blood. Cell representation was balanced among the major tissue compartments (Endo, endothelial; Immune; Epi, epithelial; Stroma) by magnetic and fluorescence activated cell sorting (MACS and FACS) using antibodies for the indicated surface markers (CD31, CD45, EPCAM; +, marker-positive; -, marker-negative). Cell capture and single cell RNA sequencing (scRNAseq) was done using 10x droplet technology or SmartSeq2 (SS2) analysis of plate-sorted cells. Number of profiled cells from each compartment are shown in parentheses. For blood, immune cells were isolated on a high density Ficoll gradient, and unsorted cells captured and profiled by 10x and sorted cells (using canonical markers for the indicated immune populations) by SS2. Total cell number (all 3 subjects) and median expressed genes per cell are indicated for each method. **b,** Cell clustering and annotation pipeline. Cell expression profiles were computationally clustered by nearest-neighbor relationships and clusters were then separated into tissue compartments based on expression of compartment-specific markers (*EPCAM* (blue), *CLDN5* (red), *COL1A2* (green), and *PTPRC* (purple)), as shown for tSNE plot of lung and blood cell expression profiles obtained by 10x from Patient 3. Cells from each tissue compartment were then iteratively re-clustered until differentially-expressed genes driving clustering were no longer biologically meaningful. Cell cluster annotation was based on expression of canonical marker genes from the literature, markers found through RNA sequencing of purified cell populations (Bulk RNA markers), ascertained tissue location, and inferred molecular function from differentially-expressed genes. **c,** Heatmap of pairwise Pearson correlations of the average expression profile of each cluster in the combined 10x dataset plus SS2 analysis of neutrophils. Tissue compartment and identification number of each of the 58 clusters are indicated.

Iterative, graph-based clustering was used to identify transcriptionally distinct clusters among cells with high quality transcriptomes separately for each subject^32^. When clustering all cells from a single subject at once, we found that the first principal components defining heterogeneity represented differences in tissue compartment, but some cell types within a compartment (e.g., basal, goblet club, neuroendocrine and ionocyte) had a tendency to co-cluster. We therefore initially grouped cells by tissue compartment based on expression of canonical compartment-specific marker genes (Fig. 1b) and then separately clustered cells within each compartment for each subject. Homologous clusters between subjects were subsequently established based on cluster-specific marker genes then merged for downstream analyses; batch correction algorithms were not needed because of centralized and robotic library preparation. Our approach identified 58 transcriptionally distinct cell populations (mean 51 per subject, Fig. 1c, Table S2). A state-of-the-art scRNAseq analysis of human lung published while this paper was in preparation identified 21 (36%) of the 58 lung cell populations described here^33^.

### Transcriptomes of nearly all of the 45 previously known human lung cell types

The 58 identified cell populations included 15 epithelial (clusters 1-15), 9 endothelial (16-24), 9 stromal (25-33), and 25 immune (34-58) populations, greater than the number of classical cell types in each compartment (11 epithelial, 5 endothelial, 7 stromal, 20 immune, and 2 neuronal cell types) (Table S2). Using extant markers for the classical cell types and/or the homologous cell types in mice (Table S1) and single molecule fluorescence *in situ* hybridization, we identified cell clusters representing nearly all of the classical lung cell types in the epithelial (club, ciliated, basal, goblet, mucous, serous, ionocyte, neuroendocrine, alveolar type 1, alveolar type 2), endothelial (pulmonary artery, vein, capillary, lymphatic), and stromal (airway smooth muscle, vascular smooth muscle, fibroblast, myofibroblast, lipofibroblast, pericyte, mesothelium) compartments (Fig. 2a,b). No markers were known for bronchial vessels, but *in situ* staining for cluster-specific markers subsequently identified bronchial endothelial clusters (see below). The only cell type not captured was epithelial tuft cells, which are rare or absent in the normal lung^34^ so was not expected to be found (Table S1). However, seven of the 23 assigned classical cell types in these three compartments were surprisingly represented by more than one cell cluster (ciliated, basal, alveolar type 2, capillary, bronchial vessel, myofibroblast, and fibroblast), revealing molecular diversity beyond the established lung cell types, as detailed below.

**Figure 2.**
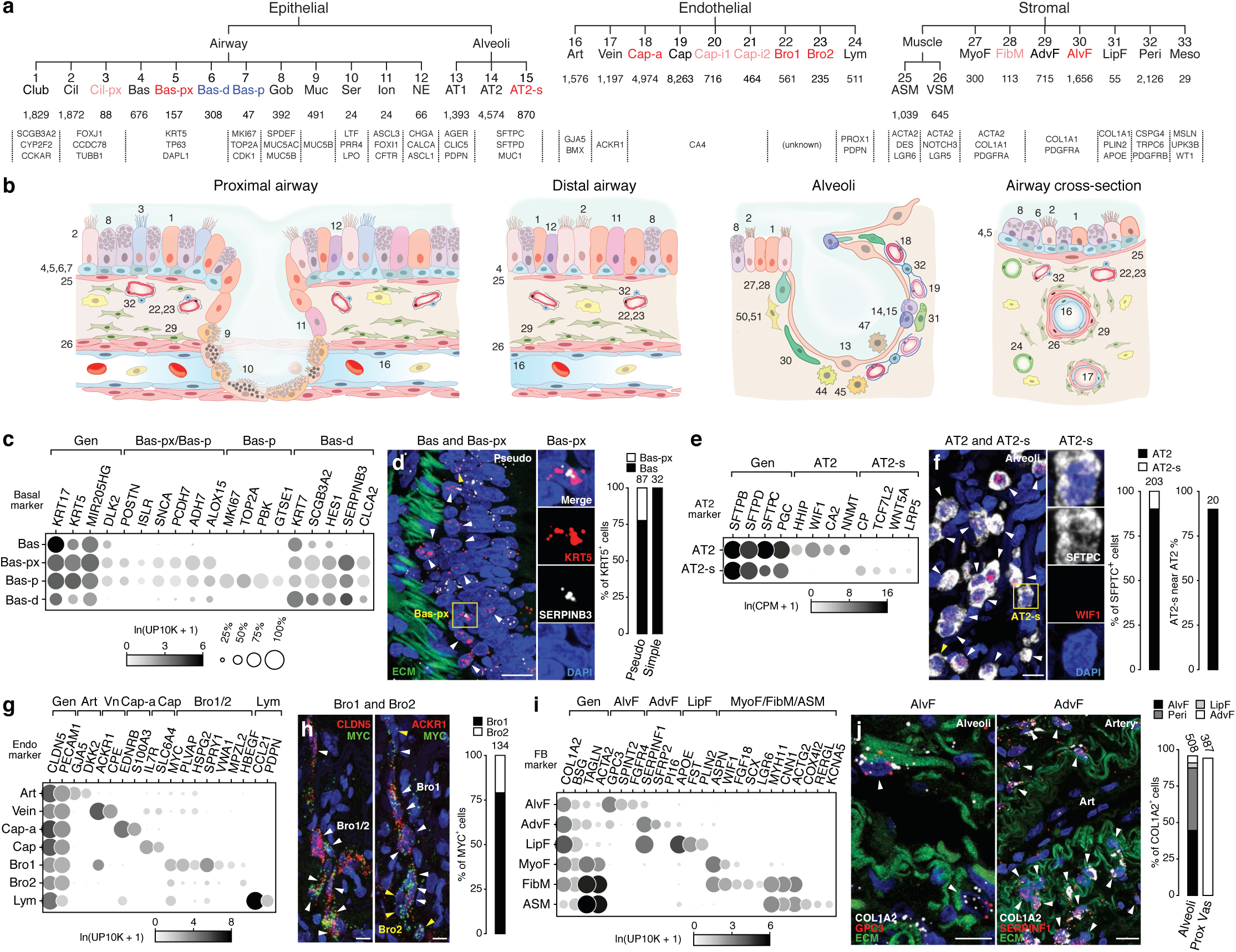
Identity and location of human lung epithelial, endothelial, and stromal cell types. **a**, Hierarchical tree showing human lung molecular cell types and their annotations in the indicated tissue compartments following iterative clustering (each level of hierarchy is an iteration) of scRNAseq profiles of cells in each compartment. Black, canonical cell types; blue, proliferative subpopulations or differentiating states; red, novel populations (light red, found only in one subject). Number of cells in each cluster and canonical marker genes are shown below. **b,** Diagrams showing localization and morphology of each lung cell type (numbering scheme as in (a) and Figure 3a). **c,** Dot plot of mean level of expression (dot intensity, gray scale) of indicated basal cell markers and percent of cells in population with detected expression (dot size). **d,** RNAscope single molecule fluorescence in situ hybridization (smFISH) and quantification for general basal marker *KRT5* (red) and Bas-px marker *SERPINB3* (white) with DAPI (nuclear) counter stain (blue) and autofluorescence (ECM, green). Scale bar, 20 µm. Note Bas-px cell (*KRT5 SERPINB3* double positive, yellow arrowhead and box) enrichment at base of pseudostratified airways. **e,** Dot plot of expression of AT2 markers. **f,** smFISH and quantification for general AT2 marker (*SFTPC*, white) and AT2-s marker *WIF1* (red puncta, arrowheads). Scale bar, 10 µm. Note AT2-s cells (*SFTPC^pos^ WIF1^neg^*, yellow arrowhead and box) intermingled with AT2 cells (*SFTPC^pos^ WIF1^pos^*, white arrowheads). **g,** Dot plot of expression of endothelial markers. **h,** smFISH for general endothelial marker *CLDN5* (red, left panel), bronchial vessel-specific markers *MYC* (green, left panels) and Bro1-specific marker *ACKR1* (red, right) on serial sections of bronchial vessel cells (arrowheads) adjacent to an airway (see Fig S4h), co-stained for DAPI (blue). Scale bar, 10 µm. Quantification shows abundance of Bro1 and Bro2 cells. **i,** Dot plot of stromal markers. **j**, smFISH and quantification for the general fibroblast marker *COL1A2* (white), alveolar fibroblast (AlvF) marker *GPC3* (red, left) and adventitial fibroblast (AdvF) marker *SERPINF1* (red, right) with DAPI counterstain (blue) and autofluorescence (ECM, green). Adventitial fibroblasts (arrowheads, k) localize around blood vessels (ECM, green). Scale bars, 10 µm. Quantification shows abundance of stromal cell types in alveolar and proximal regions, pericyte (Peri) and lipofibroblast (LipF) staining in Fig S4i-j. All stains were done on lungs from at least two human subjects distinct from those used for profiling; quantifications represent ≥10 fields of view.

Immune cells were the most heterogeneous compartment presumably because of the critical role of the lung as first responder to inhaled toxins and pathogens, and because the analyzed lung samples were not perfused so included circulating and egressed as well as lung resident immune cells. To aid assignment of identities to the immune cell clusters, we first defined the transcriptional profiles of circulating immune cells by sorting human blood cells using established surface markers followed by bulk RNA sequencing of the sorted cell populations. This defined the transcriptional profiles of 21 functionally-characterized classes of circulating immune cells (Fig. S5a, Table S3). We also obtained scRNAseq profiles of ∼5000 circulating blood cells from two of the subjects whose lung cells we analyzed. Canonical immune cell markers along with the ascertained panels of differentially-expressed genes allowed us to assign identities to 25 molecularly distinct immune cell clusters obtained in our scRNAseq analysis of the human lung samples, including all of the 20 previously known lung immune cell types except eosinophils (Fig. 3a, S5b, see below).

**Figure 3.**
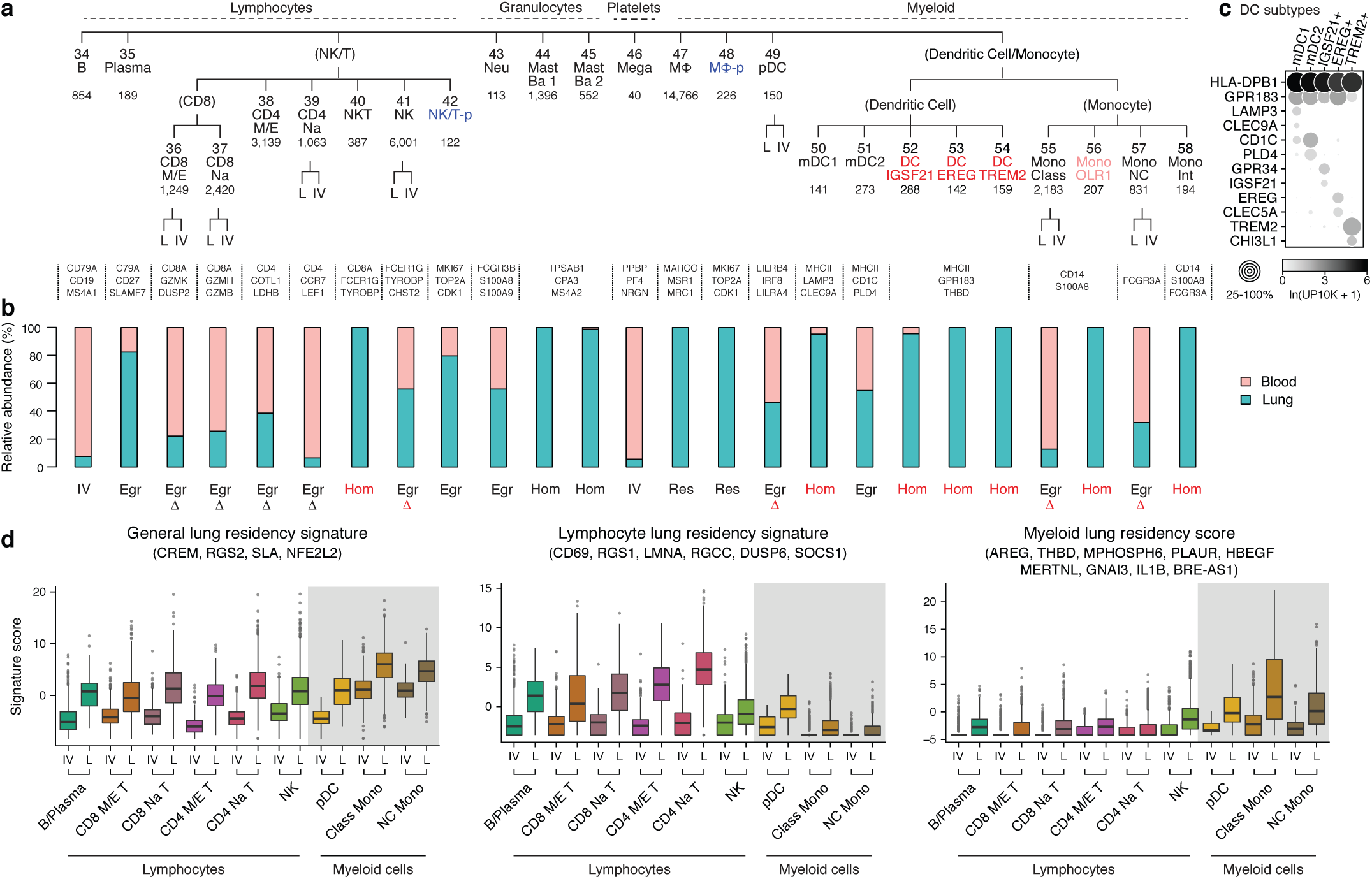
Identity and residency of human lung immune cells. **a**, Hierarchical tree showing human lung immune cell molecular types (grouped by lineage) and their annotations based on expression of marker genes in cell clusters of scRNAseq profiles as in Fig. 2a. Black, canonical cell types; blue, proliferative subpopulations; red, novel populations (light red, found only in one subject). Number of cells in each cluster (combined 10x and SS2 datasets) is indicated. Cell types showing significant expression differences based on whether cell was obtained from lung (L) vs. blood (IV, intravascular) samples are shown as additional level of hierarchy. **b,** Relative abundance of each immune cell type (shown in panel a) in lung (blue) and blood (red) samples. Cell types with >90% enrichment in lung samples are annotated “lung resident” (Res, alveolar macrophages) or “lung homing” (Hom), those with >90% enrichment in blood as “intravascular” (IV), and all others as “egressed” (Egr). (Note these assignments should be considered provisional because enrichment values can be influenced by efficiency of harvesting during cell dissociation and isolation.) Red lettering/symbols, immune cells not previously known to home to (be enriched in) lung or have differential expression (delta symbol) following egression from blood vessel into tissue. **c,** Dot plot showing expression of canonical dendritic markers (*HLA-DPB1*, *GPR183*), myeloid dendritic cell type 1 and 2 (mDC1 and mDC2) markers (*LAMP3A*, *CLEC9A*, *CD1C*, *PLD4*), and six markers for three novel dendritic populations (IGSF21+, EREG+, and TREM2+) in the dendritic immune cell clusters indicated (clusters 50-54). UP10K, UMIs per 10,000. **d,** Box-and-whisker plots of general, lymphocyte-specific, and myeloid-specific lung residency (egression) gene expression signature scores based on expression of indicated genes in 10x profiles of the indicated immune cells types isolated from the indicated sample sites (IV, from blood; L, from lung). Many previously known lymphocyte residence genes (e.g. *S1PR1*, *RUNX3*, *RBPJ, HOBIT*) were expressed at low levels and only uncovered in SS2 profiles with better transcript capture. Gray shading, myeloid immune cells.

Thus, our approach provides genome-wide expression profiles for nearly all the classical lung cell types (41 of 45, 91%), from the most abundant cell types (e.g. capillary endothelial cells, ∼23% of lung cells) down to exceedingly rare ones, with estimated abundances as low as 0.01% (e.g., neuroendocrine cells, ionocytes) (Table S1). One-quarter of the classical cell types (11 of 45, 24%) previously lacked high quality single cell expression data, either because they had never been profiled or correctly annotated (mucous, capillary, bronchial vessel, airway smooth muscle, lipofibroblast, myofibroblast) or had only been profiled after culturing (artery, pericyte, mesothelium, platelet/megakaryocyte, and intermediate monocyte) (Table S1). The only canonical cell types not captured by our approach are extremely rare and primarily found in disease settings (tuft cells) or have structures or features that require special isolation or enrichment methods (intrinsic neurons, Schwann cells, eosinophils). This comprehensive dataset of cell expression profiles suggests novel functions, signaling interactions, and contributions to disease for many canonical cell types, such as a role for lung pericytes in regulation of microvascular tone through circulating hormones and as a culprit in pulmonary hypertension (see below).

### Discovery of new molecularly-defined human lung cell types, subtypes and states

Although specific clusters were identified for nearly all canonical cell types, many cell types were surprisingly represented by more than one cluster, so the specific identities of 25 clusters remained uncertain. Most of the extra clusters were unlikely to have arisen from batch effects or subject-specific pathology because all except three clusters were found in more than one subject by scRNAseq or in situ hybridization (Table S2). This suggested that the distinct expression profiles uncovered for these cell types might represent discrete molecular states of the known cell types or previously unrecognized cell types or subtypes. To begin to distinguish these possibilities, we analyzed the differentially-expressed genes among clusters assigned to each canonical cell type to identify cluster-specific biochemical and biological functions and examined the cells’ structure and spatial distribution in the lung.

We first identified clusters representing common cell states. Three of the clusters (cluster 7, Bas-p; 42, NK/T-p; 48, MP-p) were significantly enriched in expression of cell cycle genes relative to the other cluster(s) of the same cell type, indicating that these represent the proliferative states of basal cells, NK cells, T cells, and macrophages, respectively, and these are the most proliferative cell types in the adult lung (Fig. 2c, S3a). Approximately 4% of basal cells were in the proliferative cluster (cluster 7, 47 cells), most of which were isolated from proximal lung samples (Fig. S3b); this suggests that proximal basal cells are more proliferative than distal ones, consistent with immunostains for proliferation marker KI67. (Fig. S4a). Another cluster (cluster 6, Bas-d, 308 cells), comprising 28% of all basal cells, had reduced expression of *KRT5* and increased expression of *HES1*, *KRT7*, and *SCGB3A2*, indicating that they were differentiating to other epithelial fates^35, 36^, consistent with their transitional morphology (Fig. 2c, S4b). Differentiating basal cells also derived mostly from proximal lung samples (Fig. S3b, S4b). These expression patterns and immunostaining suggest that a surprisingly^37^ large proportion of human basal cells are proliferating or differentiating, especially in the proximal airways where approximately one-third of isolated basal cells appear active.

The other basal cell clusters corresponded to quiescent basal cells and localized by smFISH and isolation site either to exclusively proximal (large, pseudostratified) airways (cluster 5, Bas-px), or to both proximal and distal (small, simple) airways (cluster 4, Bas) (Fig. 2d, S3b, S4c). Despite their similar morphology, Bas-px and Bas basal cells are distinguished by hundreds of genes, suggesting they are molecularly distinct cell types or subtypes that differ in hormone production (*ALOX15*, *ADH7*, *SNCA*) and adhesion (*POSTN*, *ISLR*, *PCDH7*), and, based on morphology and immunostaining for proliferation and differentiation markers, perhaps their ability to function as stem cells (Fig. 2c). We also found distinct clusters (clusters 2, 3) and associated molecular signatures along the proximal-distal axis for ciliated cells, indicating additional heterogeneity among epithelial cells along this axis, consistent with a recent study^33^ (Fig. S3b,c). smFISH of *DHRS9*, restricted in expression to proximal ciliated cells, localized this population to larger pseudostratified airways (Fig. S4d).

We uncovered two clusters (14, 15) of alveolar type 2 (AT2) cells (Fig. 2e), which produce surfactant that prevents alveolar collapse. These were intermingled throughout the alveolar epithelium, as shown by in situ localization of cluster-specific markers (Fig. 2f). One of the clusters (cluster 14, marked by *WIF1*, *HHIP*, *CA2*) expressed higher levels of some canonical AT2 markers (e.g., surfactant genes *SFTPA1* and *SFTPC* and transcription factor *ETV5*), and also selectively expressed inhibitors of Wnt (*WIF1*) and Hedgehog (*HHIP*) signaling as well as the cell cycle (*CDKN1A*), implying the cells are quiescent (Fig. S3d, left). The other, 10-fold less abundant cluster (cluster 15, marked by *CP* and lacking *WIF1* and *HHIP*) selectively expressed detoxification genes (*CP*, *GSTA1*, *CYP4B1*) as well as Wnt pathway genes including a ligand (*WNT5A*), co-receptor (*LRP5*), regulatory protein (*CTNNBIP1*), and canonical transcription factor (*TCF4/TCF7L2*) (Fig. S3d, right); we refer to these as AT2-signaling (AT2-s). AT2-s could be alveolar stem cells, homologous to the rare subpopulation of Wnt-active AT2 cells recently identified in mouse (AT2^stem^)^38, 39^, whereas cluster 15 could be classical AT2 cells, homologous to “bulk” AT2 cells of mouse. However, the homology and possible functional similarity between human AT2-s and mouse AT2^stem^ is provisional because although both show elevated Wnt signaling or components, the many other observed differences between human AT2-s and “bulk” AT2 are not shared by mouse AT2^stem^.

We also found unexpected molecular diversity in the endothelial compartment beyond the five canonical endothelial cell types (artery, vein, capillary, bronchial vessel, lymphatic). Two populations (clusters 22, 23) were identified as bronchial endothelial cells (Bro1, Bro2) by their localization around bronchi by in situ hybridization for the cluster-specific marker genes *MYC* and *ACKR1* as well as panendothelial marker *CLDN5* (Fig. 2g,h; S4h). This indicates that bronchial endothelial cells are molecularly distinct cell types from their counterparts in the pulmonary circulation and lymphatics, distinguished by expression of extracellular matrix (*VWA1*, *HSPG2*), fenestrated morphology^40^ (*PLVAP*) and cell cycle associated (*MYC*, *HBEGF*) genes (Fig. 2g). Four clusters of endothelial cells in the pulmonary circulation expressed capillary markers. Two of these (clusters 18, 19) are a pair of intermingled and intercommunicating but not interconverting alveolar capillary cell types (Cap-a, Cap), as shown by cluster-specific lineage-labeling and mapping in mice (Gillich et al); the other two (clusters 20, 21) are rare cell types showing mixed features of Cap-a and Cap cells that we call capillary-intermediate type 1 (Cap-i1) and 2 (Cap-i2).

We also identified new cell types or subtypes in the stroma, the least characterized lung compartment. Two cell clusters (29, 30) expressed classical fibroblast markers (*COL1A1*, *COL1A2)* (Fig. 2i) but RNA *in situ* hybridization showed one population (cluster 30, marked by *SPINT2*, *FGFR4*, *GPC3*) localized to the alveolus (“alveolar fibroblasts”) in human and mouse (Fig. 2j,left; S4k,l) and the other (cluster 29, marked by *SFRP2, PI16*, *SERPINF1*) to vascular adventitia and nearby airways (“adventitial fibroblasts”) in human and mouse (Fig. 2j,right; S4m). Both express genes involved in canonical fibroblast functions including extracellular matrix biosynthesis, cell adhesion, and angiogenesis regulators and other intercellular signals and modulators. However the specific genes for these functions often differ, for example adventitial fibroblasts expressed different Wnt pathway modulators (*SFRP2*) than those of alveolar fibroblasts (*NKD1*, *RSPO1*) (Fig. S3e). Each cluster also appears to have distinct functions: specific expression of voltage-gated sodium channel *SCN7A* and glutamate receptor *GRIA1* suggest alveolar fibroblasts are excitable cells that receive glutamatergic input (Table S4). The expression profiles also suggest novel, shared immune functions including immune cell recruitment (*IL1RL1*, *IL32*, *CXCL2*, *MHCII*) and the complement system (*C2*, *C3*, *C7*, *CFI*, *CFD*, *CFH*, *CFB*) (Table S4).

Two other stromal clusters (clusters 27 and 28) were enriched for *ACTA2*, a canonical marker of myofibroblasts (Fig. 2i), cells that help form alveoli during development and can act inappropriately later in disease. One of these (cluster 27, marked by *WIF1*, *FGF18*, *ASPN*) are classical myofibroblasts, which were isolated from distal lung samples and localized to alveolar ducts (Fig. S4o). The other population (cluster 28) showed higher expression of contractile genes (*MYH11*, *CNN1*, *TAGLN*), was preferentially isolated from proximal lung samples, and was found both intermingled with airway smooth muscle and in alveoli (Fig. S3b,S4n), so we call them “fibromyocytes.” Both populations shared expression of genes for canonical fibroblast functions (e.g. ECM genes *VCAN*, *COL12A1*, *CLU*), though the specific genes differed from both alveolar and adventitial fibroblasts, and they also both expressed a rich set of genes associated with TGF-beta signaling (*LTBP1*, *LTBP2*, *ASPN*, *DPT*, *TGFBR3*, *TGFBI*, *SCX*, *MDFI*) (Table S4).

### Profiles of lung immune cells and blood cells identify lung homing and residency signatures

To distinguish lung resident and itinerant immune cell clusters from intravascular cells circulating through the lung (Fig. 3a), we first compared the relative abundance of each immune cell population in lung samples and matched peripheral blood from the same subject (Fig. 3b). Eleven of the clusters were comprised of cells only from lung samples, with no or only rare exception (clusters 40, 44, 45, 47, 48, 50, 52, 53, 54, 56, 58). This indicated that not only are alveolar macrophages (clusters 47, 48), myeloid dendritic type 1 cells (50), and basophil/mast cells (44, 45) lung resident^41^ or greatly enriched in the lung interstitium or lumen as expected, but surprisingly so are natural killer T cells (40) and intermediate monocytes (58). We also discovered four novel lung myeloid populations (52, 53, 54, 56) that are lung resident or greatly enriched. These appear to have specialized functions and localizations: IGSF21+ dendritic cells express genes implicated in asthma (*CCL2*, *CCL13, IGSF21*) and localize proximal vessels, rare EREG+ dendritic cells are enriched in developmental signals (*EREG*, *VEGFA*, *AREG*) and also localize to proximal vessels, and TREM2+ dendritic cells express lipid handling machinery (*APOC1*, *APOE*, *CYP27A1*) and localize to both vessels and alveoli (Fig. 3c, S5d-g).

The other immune cell types were found in both the lung and blood samples. For some cell types, every cell—whether from blood or lung— clustered together. This included B cells (cluster 34), plasma cells (35), platelets/megakaryocytes (46), plasmacytoid dendritic cells (49), and neutrophils (43). For other cell types (every T cell subset, natural killer cells, myeloid dendritic type 2 cells, and both classical and nonclassical monocytes), the cells from lung formed a separate cluster from those from the blood. Some of the differentially-expressed genes may be due to technical differences (e.g., collagenase treatment of lung but not blood^42^ or circulating RNA in blood^43^), but other detected differences such as upregulation in lung cells of lymphocyte-residence gene *CD69* likely represent genes induced following egression^44^. We identified a core transcriptional signature for all human lung resident lymphocytes (e.g. *CD69*, *LMNA*, *RGCC*), which partially overlaps with a residence signature recently defined by bulk RNAseq of CD8+ T cells in the mouse spleen, gut and liver^45^ (Fig. 3d). We also discovered a residency signature for lung myeloid cells (e.g. *AREG*, *MPHOSPH6*, *HBEGF*). This signature partially overlaps with the one for lymphocytes, supporting a core residency program for immune cells as well as specific subprograms for myeloid cells and lymphocytes.

### Lung cell type markers, transcription factors, hormone targets, and signaling interactions

Our nearly complete gene expression atlas for the adult human lung has important implications for medicine. We identified the most selective marker genes for each of the previously known and newly identified cell types (Fig. 4a, Table S4). A battery of ∼200 markers can unambiguously distinguish virtually all lung cell types (Fig. 4b), so could be used with multiplex in situ hybridization methods^46–48^ to simultaneously detect in clinical specimens any alterations in their numbers, structures, and spatial relationships, ushering in a new era of diagnostic pathology. A similar compendium of cell-type-selective membrane proteins (Table S4) could be used to purify or therapeutically target specific lung cell types. We also identified the transcription factors selectively enriched in each cell type (Fig. 4e, Table S4), putative “master regulators” that can be used to create each cell type by cellular reprogramming, and whose efficacy in reprogramming can be evaluated with the markers described above. We found ∼400 such transcription factors (1 to 36 for nearly every cell type in our SS2 data), over half of which are novel (Fig. 4e, red text). This includes what may be the long-sought master regulators (such as *MYRF*) of alveolar epithelial type 1 (AT1) cells, which comprise nearly all of the lung’s gas exchange surface and are impacted in many important lung diseases, and a regulator of pericyte identity (*TBX5*). We confirmed specific expression of *MYRF* in AT1 cells and *TBX5* in pericytes by single molecule *in situ* hybridization (smFISH) (Fig. 4c,d). These ∼400 cell-type selective transcription factors are excellent candidates for creating differentiation protocols for nearly all the cell types needed to engineer a human lung.

**Figure 4.**
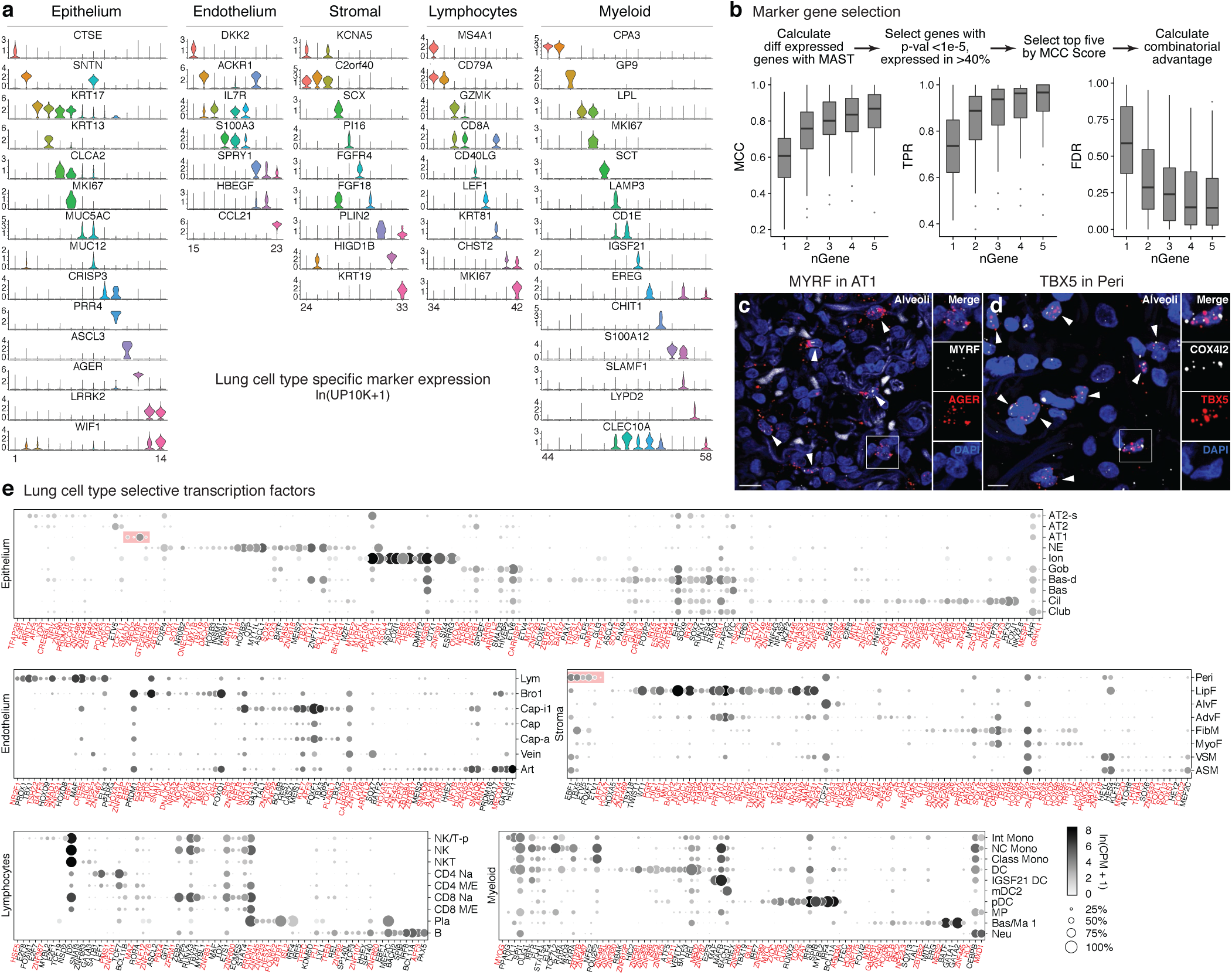
Markers and transcription factors that distinguish human lung cell types. **a,** Violin plots of expression levels (ln(UP10K + 1)) in 10x profiles of the most sensitive and specific markers (gene symbols) for each human lung cell type in its tissue compartment. **b,** Scheme for selecting the most sensitive and specific marker genes for each cell type using, Matthews Correlation Coefficient (MCC). The box-and-whisker plots below show MCCs, True Positive Rates (TPR), and False Discovery Rates (FDR) for each cell type using the indicated number (nGene) of the most sensitive and specific markers. Note all measures saturate at 2-4 genes, hence simultaneous in situ probing of a human lung for the ∼100-200 optimal markers would assign identity to nearly every cell. **c,** Alveolar region of human lung probed by smFISH for AT1 marker *AGER* and transcription factor *MYRF* mRNA. *MYRF* is selectively expressed in AT1 cells (arrowheads; 97% of *MYRF*+ cells were *AGER*+, n=250 scored cells). Scale bar, 10 µm. **d,** Alveolar region of human lung probed by smFISH for pericyte marker *COX4I2* and transcription factor *TBX5* mRNA. *TBX5* is enriched in pericytes (arrowheads, 92% of *TBX5*+ were *COX4I2*+, n=250). Scale bar, 5 µm. **e,** Dot plot of expression of enriched transcription factors in each lung cell type from SS2 profiles. Red, genes not previously associated with the cell type. Red shading, transcription factors including *MYRF* that are highly enriched in AT1 cells, and *TBX5* and others highly enriched in pericytes.

The expression atlas allowed us to map the direct cell targets in the lung of circulating hormones, based on expression patterns of their cognate receptors. Receptors for some hormones (e.g., glucocorticoid receptor *NR3C1*, insulin receptor *INSR*, IGF receptor *IGF1R*, and adiponectin receptors *ADIPOR1* and *ADIPOR2*; all related to cell growth) are broadly expressed, implying direct action of these hormones throughout the lung, with some notable exceptions such as lymphocytes (Fig. 5a). Other hormones have highly specific and unexpected targets, such as somatostatin (*SSTR1* expressed only in arteries), mineralocorticoids (*NR3C2*, goblet cells), parathyroid hormone (*PTH1R*, smooth muscle, pericytes), ghrelin (*GHSR*, neuroendocrine cells), melanocortin (ionocytes and lipofibroblasts), and gastric inhibitory peptide (*GIPR*) and oxytocin (*OXTR*) (ciliated cells). Pericytes are predicted targets of angiotensin, endothelin, parathyroid hormone, and prostacyclin, which could regulate their contractile machinery to tune alveolar perfusion (Fig. 5b) similar to vascular smooth muscle control of arterial flow. The receptor genes for half of all circulating hormones (e.g. opioids, human growth hormone, and glucagon) were not detectably expressed in any lung cell type so may not directly influence lung physiology.

**Figure 5.**
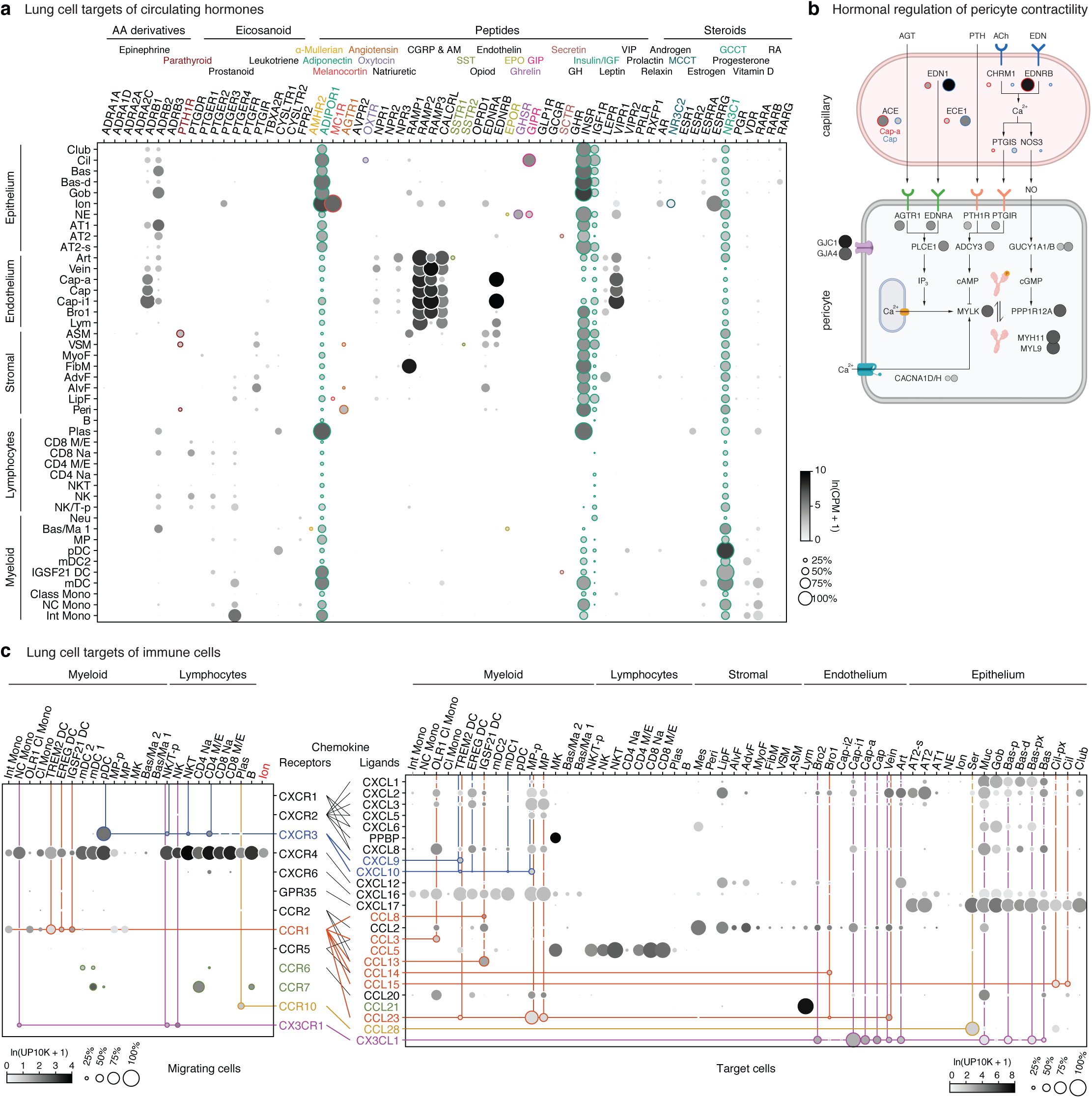
Lung cell targets of circulating hormones and immune cells. **a**, Dot plot of hormone receptor gene expression in lung cells from combined SS2 profiles. Hormones for each receptor are shown at top. Teal, broadly-expressed receptors in lung; other colors, selectively-expressed receptors (<3 lung cell types). Small colored dots near cell types show selectively targeted cell types. AA, amino acid; CGRP, Calcitonin gene-related peptide; AM, adrenomedullin; SST, somatostatin; EPO, erythropoietin; GIP, gastric inhibitory peptide; GH, growth hormone; IGF, insulin-like growth factor; MCCT, mineralocorticoid; GCCT, glucocorticoid; RA, retinoic acid. **b,** Schematic of inferred pericyte cell contractility pathway and its regulation by circulating hormones (AGT, PTH) and capillary-expressed signals (EDN, NO). Dots show expression of indicated pathway genes: values at left (outlined red) in each pair of dots in capillary diagram (top) show expression in Cap-a cells (aerocytes) and at right (outlined blue) show expression in general Cap cells. Note most signal genes are preferentially expressed in Cap relative to Cap-a cells. **c,** Dot plots of expression of chemokine receptor genes (left) and their cognate ligand genes (right) in human lung cells from combined 10x expression profiles. Only cell types and chemokines with detected expression are shown; ionocytes (red lettering) are the only non-immune cell with chemokine receptor expression. Colored lines highlight predicted target cells (chemokine-expressing cells, right) of migrating immune cell types and ionocytes (left) that express a cognate chemokine receptor gene; heavier weight lines indicate previously unknown interactions.

We next used the expression map to predict other signaling interactions among lung cells by identifying complementary expression patterns of ligands and their receptors among cells in the airways and accompanying vessels and among alveolar cells using CellPhoneDB^25^ (see Methods). This analysis predicts diverse sets of up to hundreds of interactions of each lung cell type and its neighbors, with stromal cells the leading interactors largely due to growth factor-mediated signaling interactions (Fig. S6).

Expression of chemokine receptors provides insight into immune cell homing in the lung (Fig. 5c). Our data confirmed canonical homing interactions such as expression of *CCR7* by CD4+ T cells guiding them to *CCL21*-producing lymphatic vessels (green in Fig. 5c), and provides specificity for others such as *CCR10*-mediated plasma cell homing to epithelial mucosa through *CCL28* from serous cells in submucosal glands (gold). It also predicts new interactions such as *CXCR3*-mediated homing of pDCs to *CXCL9*-expressing TREM2+ dendritic cells (blue), and *CX3CR1*-mediated homing of nonclassical monocytes to *CX3CL1*-expressing endothelial cells and airway epithelial cells (purple). The novel dendritic populations do not express *CCR6* and *CCR7*, which guide myeloid dendritic cells to *CCL21*-expressing lymphatic vessels, suggesting they do not egress via lymphatics. However, all three express *CCR1* (orange), which could mediate their attraction to veins that express ligand *CCL23*, as well as to bronchial endothelial cells (*CCL14*), ciliated cells (*CCL15*), and lymphocytes (*CCL5*). We also identified ionocytes (bold, red), the main source of *Cftr* expression in the mouse airway, as the only non-immune cell to express appreciable levels of any chemokine receptor *(CXCR4)*, whose ligand (CXCL12) is curiously expressed almost exclusively by non-immune cells.

### Expression map of human lung disease genes

We used the lung atlas to map the cellular sites of expression of the 233 extant genes that cause or contribute to human lung disease, curated from the Online Mendelian Inheritance in Man (OMIM) database^49^ (179 genes) and GWAS studies^50^ (54 genes, -log(p-value) > 20, see Methods). Disease genes that showed cell-type-specific expression within the lung are of special interest (Fig. 6a), because they can pinpoint the cell type(s) in which the disease originates, especially significant for the many lung diseases whose cellular origins are poorly understood.

**Figure 6.**
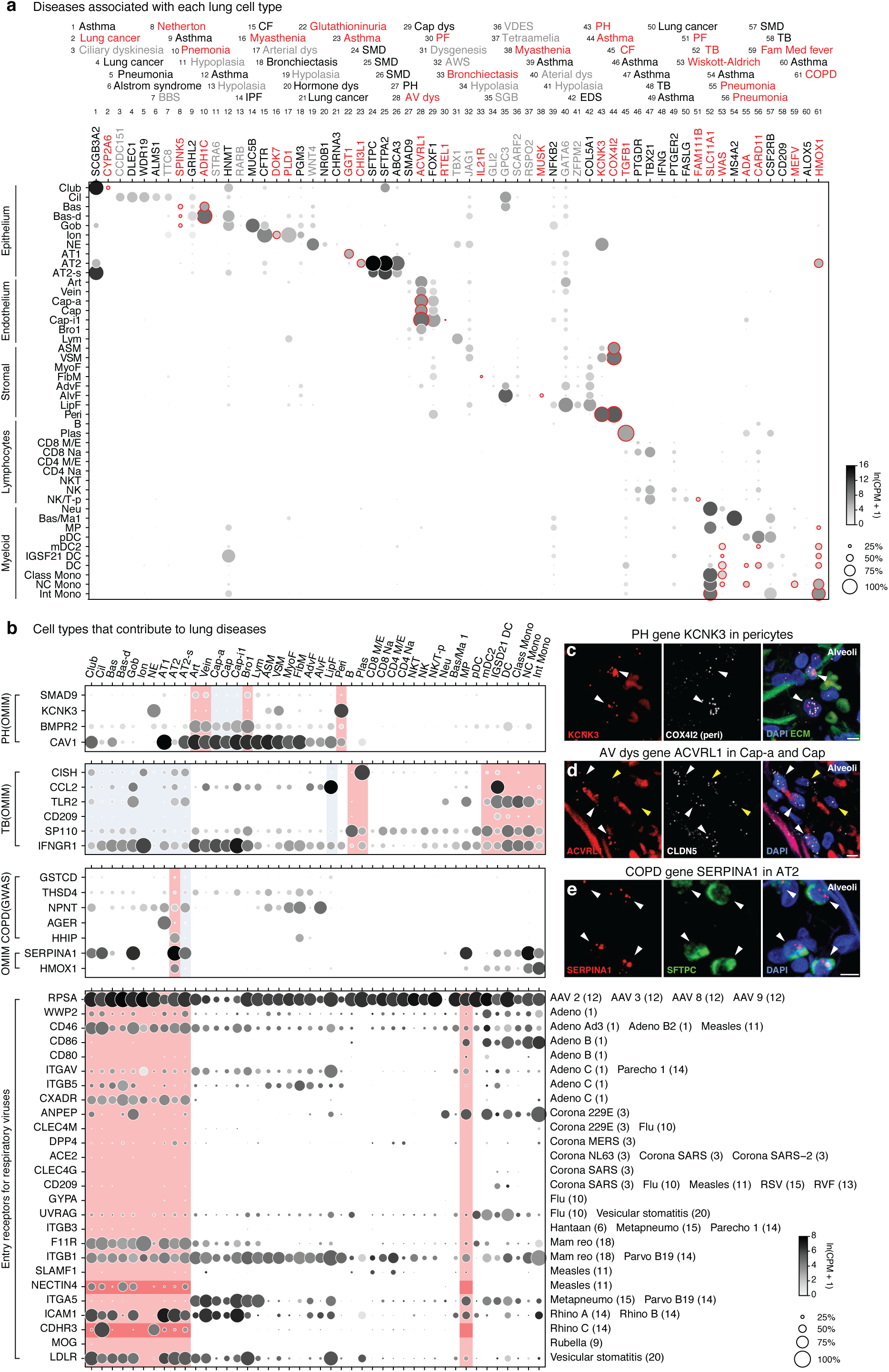
Mapping the cellular origins of human lung diseases by cell-selective expression of disease genes. **a**, Dot plots of expression of lung disease genes (numbered with the associated disease indicated above) that are enriched in specific lung cell types from combined SS2 expression profiles. Red, novel cell type association of gene/disease; gray, diseases with developmental phenotype. BBS, Bardet-Biedl syndrome; Dys, dysplasia; IPF, idiopathic pulmonary fibrosis; SMD, surfactant metabolism dysfunction; PH, pulmonary hypertension; SM, smooth muscle; SGB, Simpson-Golabi-Behmel; TB, tuberculosis; AWS, Alagille-Watson syndrome; VDES, Van den Ende-Gupta syndrome; EDS, Ehlers-Danlos syndrome; CF, Cystic fibrosis; Fam Med, Familial Mediterranean; COPD, Chronic Obstructive Pulmonary disease. **b,** Dot plots of expression of all genes implicated (OMIM, Mendelian disease gene from OMIM database; GWAS, genome-wide association at 10^-20^ significance) in the indicated disease (PH, pulmonary hypertension, first panel; tuberculosis susceptibility, second; COPD/emphysema, third). Red shading, cell type strongly implicated in disease; blue shading, weaker but still notable association. Note canonical AT2 cells (red) express every major COPD/emphysema disease gene whereas AT2-s cells (blue) express only some. Fourth panel, dot plot showing expression of genes used by viruses (listed on the right) to enter the lung. Fourth panel, dot plot showing expression of genes used by viruses indicated at right to enter lung. Red shading, cell types inhaled viruses could directly access (epithelial cells and macrophages); darker red shading shows expression values for measles receptor *NECTIN4* (measles receptor) and rhinovirus C receptor *CDHR3*. **c,** smFISH of an alveolar region of adult human lung for PH disease gene *KCNK3* (red), pericyte marker *COX4I2* (white), and DAPI counterstain (blue). Note pericyte-specific expression (arrowheads, 91% of pericytes were *KCNK3*+, n=77). Scale bar, 5 µm. **d,** smFISH of alveolar region of an adult human lung for atrioventricular (AV) dysplasia gene *ACVRL1* (red), endothelial marker *CLDN5* (white), and DAPI. Note *ACVRL1* and *CLDN5* double positive capillaries (white arrowheads, 70% of capillaries were *ACVRL1*+, n=102) with some *CLDN5* single positive capillaries (yellow arrowheads). Scale bar, 5 µm. **e,** smFISH and quantification of alveolar region of adult human lung for COPD/emphysema gene *SERPINA1* and AT2 marker *SFTPC*, and DAPI. Note AT2-specific expression (arrowheads, 93% of AT2 cells were *SERPINA1*+, n=176). Scale bar, 5 µm.

This supported the known or suspected ‘culprit’ cells for 27 genes related to 12 diseases, such as arteries in pulmonary hypertension (*SMAD9*, *BMPR2*, *CAV1*; Fig. 6b), ciliated cells in ciliary dyskinesia (*CCDC151*), and capillaries in alveolar capillary dysplasia (*FOXF1*). It also identified potential novel culprit and contributing cells for 21 genes implicated in 15 diseases, including pericytes in pulmonary hypertension (potassium channel *KCNK3*), capillaries in atrioventricular dysplasia (*ACVRL1*), and AT2 cells in COPD (*SERPINA1*, *HMOX1*, *HHIP*). We confirmed pericyte, capillary, and AT2 expression of disease genes by smFISH (Fig. 6c-e).

We also mapped lung expression of 80 genes encoding proteins implicated as cellular receptors for human viruses (Fig. S8a), including 26 used by airborne viruses that enter primarily through epithelial cells lining the respiratory tract (Fig. 6b). *NECTIN4*, a receptor for measles virus was enriched in club, ciliated, Bas-D, and goblet cells, and *CDHR3*, the receptor for common-cold Rhinovirus C was enriched in ciliated and neuroendocrine cells, implying viral tropism and pathogenesis through these bronchial cell types. By contrast, *ACE2*, used by SARS and COVID-19 coronaviruses, and *DPP4* used by MERS coronavirus, were both detected at low levels but enriched in alveolar AT2 cells (Fig S8b), consistent with clinical data indicating lung pathology and respiratory distress of these epidemic/pandemic-prone diseases initiates in the alveolar region^51^.

### Loss, gain and diversification of lung cell types and gene expression patterns in evolution

Our recent construction of a mouse lung cell atlas from young, middle-aged and elderly adult animals^19^, supplemented with additional mouse lung cells and annotated as above for human lung (see Methods), allowed us to analyze the evolutionary conservation and divergence of lung cell types at single cell resolution and across the transcriptome. Homologous cell types were identified by conserved expression of cell-type-specific markers aligned by HomologyIDs (Fig. 7a). A striking result was that mice appear to lack 17 (29%) of the 58 molecularly-defined human lung cell populations, including many of the newly identified human molecular types (proximal ciliated (3); proximal (5), differentiating (6), and proliferating (7) basal; AT2-s (15); Cap-i1 (20) and i2 (21) capillary cells; bronchial vessel 1 (22) and 2 (23); fibromyocytes (28), lipofibroblasts (31); IGSF21+ (52), EREG+ (53), TREM2+ (54) dendritic cells; and OLR1+ monocytes (56)). Some of the missing cell populations might be rare (e.g. Cap-i1 and i2 as well as Bro1 and 2), transient (e.g. Bas-d and Bas-p), unstable (e.g. LipF), or too diverged to relate transcriptionally (e.g. TREM2+ DC) in mice so may be uncovered by further scRNA sequencing and functional follow up. By contrast, only five mouse molecular types, all immune cells (Zbtb32+ B, Ly6g5b+ T, Alox5+ T, Ccr7+ DC, and interstitial macrophages), were not found in human. These results suggest that there has been substantial molecular diversification and specialization of lung cell types during human evolution (or reduction and streamlining during mouse evolution). This also has important implications for medicine because mice would not be expected to accurately model any human disease for which an implicated cell type was missing or substantially altered.

**Figure 7.**
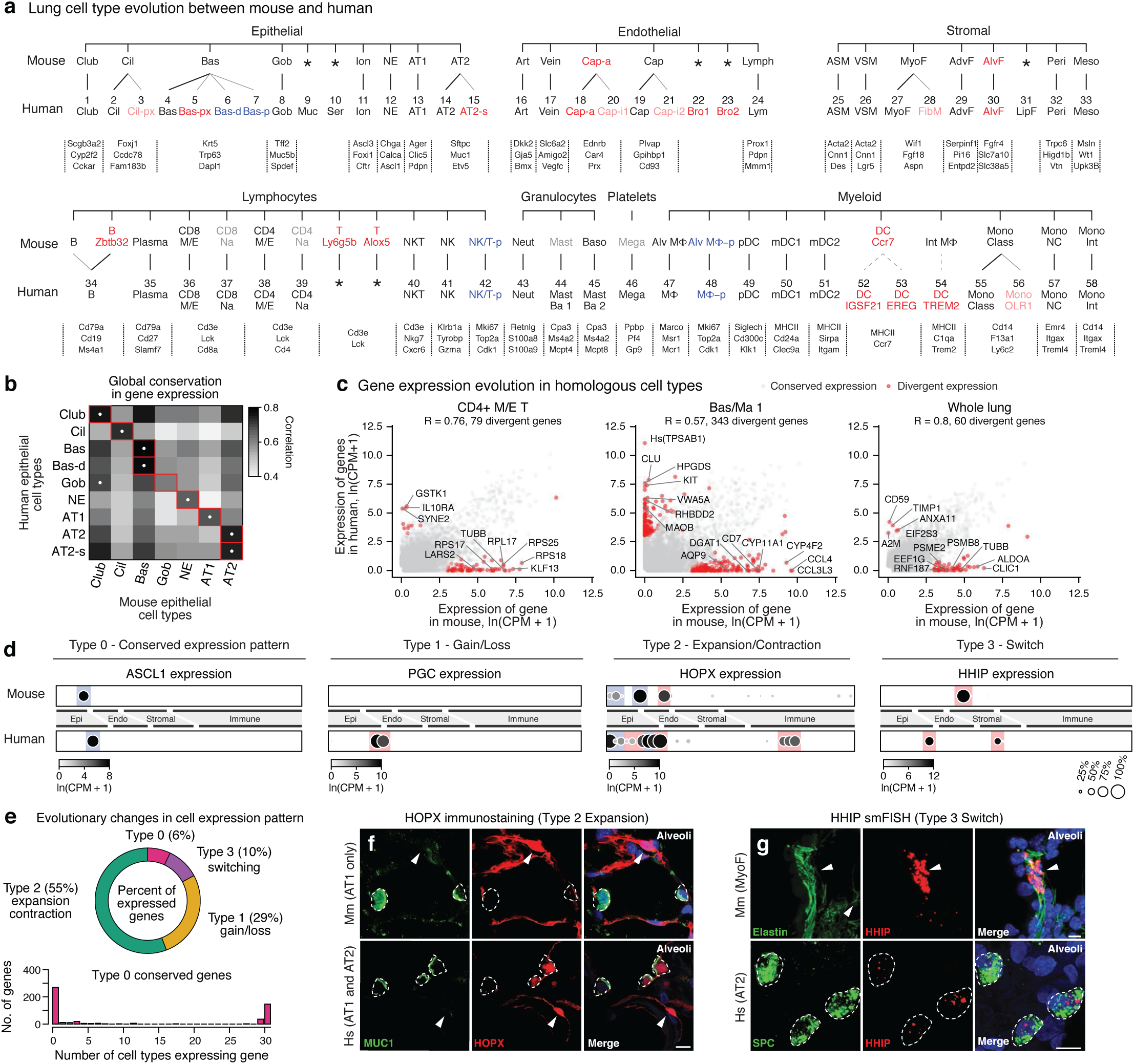
Evolutionary divergence of lung cell types and gene expression patterns. **a,** Alignment of mouse (top) lung cell types isolated, profiled by scRNAseq and clustered from each tissue compartment (see Methods) as for human (bottom, from Figs. 2a, 3a). Lines connect homologous cell types expressing classical markers shown at bottom. Thin lines indicate expansions; dashed lines, potential expansion of functionally related cell types. Note some cell types (34 total) have a unique match whereas others (8 total) have multiple matches (indicating cell type diversification during human evolution) or no identified match in mice (5 total, suggesting cell type gain in human lineage). Red text, newly identified populations (light red, identified in only 1 subject so could be subject- or disease-specific); blue, cell states more abundant in human lung; gray, extant mouse cell types not captured in our data. **b,** Heatmap showing global transcriptome correlation between indicated human and mouse epithelial cells from SS2 dataset. Red outline, matched cell types based on classical markers described in (a). White dot, human to mouse correlation. **c,** Scatter plots showing the median expression levels (dots) in the indicated cell types of each expressed human gene and mouse ortholog in the mouse and human SS2 datasets. Scale, ln(CPM+1). Note Bas/Ma 1 cells have the most differentially-expressed genes (343), and CD4^+^ M/E T cells have the least (79), but both have more than whole lung (60)—indicating genes lost in cell types are usually gained in others. Correlation scores (R values) between the average mouse and human gene expression profiles for each cell type are indicated. **d,** Dot plots of expression of homologous genes in mouse and human lung cells that exemplify the four scenarios observed for evolution of cellular expression pattern. Perfectly conserved cellular expression pattern for *ASCL1* (“Type 0” evolutionary change, NE-specific), evolutionary gain in expression in human lineage for *PGC* (acquired expression in AT2 and cells, Type 1 change), evolutionary expansion in expression from AT1, club, and ciliated cells for *HOPX* into AT2, AT2-s and other cell types indicated in human lineage (Type 2), and evolutionary switch in expression for *HHIP* from mouse myofibroblast to human AT2 cells and fibromyocytes (Type 3). Blue highlight, conserved cell expression; red, divergent cell expression. **e,** Pie chart of fraction of expressed genes in lung showing each of the four types of evolutionary changes in cellular expression patterns from mouse to human. Histogram below shows number of lung cell types that the 602 genes with perfectly conserved cellular expression patterns (Type 0) are expressed in. Note nearly all conserved genes are expressed in either a single cell type (67%) or nearly all cell types (33%). **f**, Alveolar sections from mouse (top) or human (bottom) immunostained for transcription factor HOPX (red) and AT2 marker MUC1 (green), and DAPI (blue). Note *HOPX* is expressed selectively in AT1 cells (arrowheads) in mouse but expression in humans has expanded to AT2 and AT2-s cells (dashed circles). Scale bars, 10 µm. **g**, Alveolar sections from mouse (top) and human (bottom) probed by smFISH for *Hhip and HHIP* (red) and hydrazide staining for myofibroblast marker elastin (green) in mouse and smFISH for AT2 marker *SFTPC* in human. Note evolutionary switch in *HHIP* expression from myofibroblast (mouse, arrowhead) to AT2 cells (human, dashed circles). Scale bars, 10 µm.

To characterize the conservation in the transcriptomes of homologous cell types, we compared expression levels of all active genes in each human cell type with the expression levels of the orthologous genes in the corresponding mouse cell type (Fig. S9). Most cell types showed highly conserved expression patterns as expected, with CD4+ memory/effector T cells showing the greatest conservation (correlation coefficient 0.76) and plasma cells the least (0.56). Surprisingly, one human cell type (goblet cells) showed greater correlation with another mouse cell type (club cells, 0.68) than with the homologous type (goblet cells, 0.63) (Fig. 7b), despite the conserved expression in goblet cells of canonical markers and the master transcription factor *SPDEF* (Fig. S10a). Corresponding cell types in human and mouse diverged significantly in the expression (≥20-fold change, p < 0.05) of tens to hundreds of homologous genes, with CD4+ M/E T cells showing the fewest diverged genes (79) and basophils the most (343) (Fig. 7c). The lung as a whole had fewer divergently-expressed genes (60) than any one cell type, suggesting expression lost in one cell type is usually gained in another (Fig. 7c). Diverged genes varied above age-related expression changes in mice (Fig. S11) and included canonical cell type markers, transcription factors, signaling molecules, and disease genes (see below).

Evolutionary changes in gene expression occurred in several different cellular scenarios (Table S7). Type 0 (“conserved”) genes are expressed in exactly the same cell types in mouse and human lungs (Fig. 7d, S10b). Type 1 (“expression gain/loss”) genes show simple gain (or loss) of expression in the lung between species, which could involve a single cell type (Type 1a, e.g. *PGC* expressed in human, but not mouse, AT2 cells, Fig. 7d), multiple cell types including full tissue compartments (Type 1b, e.g. *RNASE1* expressed in human, but not mouse, epithelial and endothelial cells, Fig. S10c), or the entire lung (Type 1c, *TRIM38* expressed in all human, but not mouse, lung cell types, Fig. S10c). Type 2 (“expression expansion/contraction”) changes involve gain (or loss) of expression in additional lung cell types such that expression of the gene expanded (or contracted) in the lung during evolution. For example, Hopx, the canonical AT1-selective transcription factor in mouse, is expressed in both AT1 and AT2 cells in human (Fig. 7d,f), implying the existence of other key AT1 transcription factors (such as *MYRF* (see above), which shows AT1-selective expression in both species (Fig. S9g)) and important changes in the AT2 expression program; and expansion in endothelial cell type expression of *RAMP3*, a co-receptor for vasodilators CGRP and adrenomedullin (ADM), that presumably alters the pulmonary vascular response to these hormones (Fig. S10d).

Type 3 (“expression switch”) changes are the most surprising scenario, involving a switch in gene expression from one lung cell type to another during evolution. Two medically important examples are the COPD/emphysema disease genes *SERPINA1* and *HHIP*, both of which are selectively expressed in alveolar AT2 cells in human but in alveolar stromal cell types in mice (*Hhip*, myofibroblasts, Fig. 7d,g; *Serpina1a-e* (combined), pericytes, Fig. S10e). Other components of the hedgehog pathway were mostly conserved (Fig S10f). Extreme examples of expression switching occurred during evolution of species-specific lung cell types, such as the consolidation in expression of anti-bacterial enzymes *LTF*, *LYZ* and *BPIFB1* from multiple mouse airway epithelial cells into human-specific serous cells for airway defense, and consolidation of broadly expressed lipid-handling genes (e.g. *PLIN2*, *APOE*) from mouse alveolar fibroblasts (some of which contain lipid droplets) and myofibroblasts to the human-specific lipofibroblast to support surfactant production (Fig. S10h).

Despite the general conservation of cell type gene expression patterns noted above, only ∼6% of expressed genes showed fully conserved cellular expression patterns in the lung (Type 0), and most of these were genes broadly expressed across the lung (Fig. 7e). Thus, the expression patterns of nearly all (94%) genes are labile during organ evolution, with most genes undergoing broadening (∼55%, Type 2 change) or simple gain/loss (29%, Type 1) of expression, and more rarely cell type switching (10%, Type 3) (Table S8). It will be important to unravel the genetic mechanisms underlying such widespread evolutionary changes in gene expression and the selective forces operative for the small number of genes with conserved expression, and the consequences for cellular function and mouse models of human diseases.

## Discussion

Using droplet and plate-based single cell RNA sequencing, we constructed a nearly complete molecular cell atlas of the human lung comprising 58 molecular types (Fig. 2b). This provides expression profiles for 41 of the 45 known human lung cell types, missing just the exceedingly rare intrinsic neurons and glia (estimated ∼0.001% of lung cells) and tuft cells and eosinophils that are prominent only in diseased lungs. We identified 14 novel cell populations distributed across the four tissue compartments (epithelial, endothelial, stromal, and immune) that were as distinct molecularly as the canonical cell types. These include an abundant alveolar epithelial cell type (AT2-s) intermingled with the classical cell types (AT1 and AT2), which may have stem cell function; two molecularly distinct and intermingled alveolar capillary cell types (Cap-a, Cap); new fibroblast subtypes including alveolar and adventitial fibroblasts and ones we call “fibromyocytes” that share markers with airway smooth muscle; and three new resident dendritic cell populations (DC-IGSF21+, EREG+, TREM2+). Each of the newly identified molecular cell types must be thoroughly characterized to define their structures, functions, stabilities, development, and role in disease, as we have done for the two types of alveolar capillary cells (Gillich et al.). If there are other human lung cell types, subtypes, or cell states, they must be exceedingly rare, fragile, region-(lobe) or stage-specific (younger or older donors), or so similar to the 58 molecular types described here that they are not resolved by current profiling and clustering methods.

This comprehensive molecular cell atlas has important implications for human physiology and medicine. The cellular expression patterns predict tens to hundreds of signaling interactions between each of the cell types, many of them new and unexpected. The atlas suggests where hormones act, including 12 hormones with specific cell targeting inferred from the expression pattern of their cognate receptors. It provides the relative abundance of each immune cell type isolated from the lung and circulation—and allowed us to identify lung resident and homing cell types and to infer the expression changes they undergo following egression as well as the cellular sources of the homing signals. The atlas provides expression maps of the hundreds of genes that have been implicated genetically in lung diseases and as receptors for human viruses, and uncovered dozens of genes with highly specific lung expression patterns that point to likely “culprit cells” or targets in 28 lung diseases and respiratory infections. It also identified the transcription factors selectively expressed in lung cell types that can be used to rationally design cellular reprogramming strategies for tissue engineering of an entire lung.

Our recent creation of a similar lung atlas in mouse provided an unprecedented opportunity in evolutionary biology to comprehensively compare lung cell types and their molecular functions across species^19^. This revealed that mice lack 17 of 58 (∼29%) of the molecular cell types and subtypes in the human lung, including most (∼86%, 12 of 14) of the newly discovered molecular types. This suggests a dramatic expansion of cell types in the human lineage (or streamlining in the mouse lineage), perhaps to provide new functions, durability, or regenerative capacity for our much larger (6000-fold) lungs and 30-times longer lifespan^52, 53^.

Even homologous cell types diverged in expression of tens to hundreds of genes. Indeed, we found that gene expression is remarkably labile during evolution, with only 7% of expressed genes showing conserved cellular expression patterns. Most genes (∼55%) have undergone evolutionary broadening or consolidation in their expression pattern, and some (29%) show simple gain or loss of lung expression; the rest (10%) switched cell types. These results suggest widespread gain, loss, or conversion of cell-type-specific transcriptional enhancers during mammalian evolution. Expression changes included important transcription factors, signaling molecules, and disease genes—highlighting the need for caution when using mouse models. Indeed, the atlases predict where mouse models will fail, such as for human diseases where a culprit cell is missing in mouse or does not express the homolog of the disease gene.

Our approach improves on state-of-the-art methods that recently uncovered just over a third (21) of the 58 lung molecular types described here^33^, and provides a paradigm for the human cell atlas project that can be easily adapted for other organs. Best practices that served this project and should be useful for other organ atlases include: compiling from the literature a table of known cell types, their relative abundance, and extant markers; obtaining fresh samples intraoperatively (or immediately post-mortem) from multiple positions in the organ along with matched peripheral blood; balancing tissue compartments after dissociation to ensure adequate representation of all cell types including difficult to dissociate and fragile cells; and profiling large numbers of cells (tens of thousands) from each of several subjects with both broad cell coverage (90%, e.g. microfluidic droplet sequencing) and deep gene coverage (10%, e.g., plate-based SmartSeq2). Subject and tissue compartment data should be clustered separately and iteratively until the obtained subclusters are no longer distinguished by biologically meaningful genes. Clusters should then be identified using extant marker genes, characterized by the functions of selectively-expressed genes, and histologic structures and locations of the cells in the organ confirmed by in situ hybridization, immunostaining for cluster-specific markers, or sampling location.

The tissue location of each cell cluster, including most of the newly identified molecular types, provides an initial atlas of not only cell identities and expression profiles but their environment, which enhances the value of each transcriptome by suggesting local interactions between cell types such as targets of the signals they express and sources of signals for the receptors they express. Perhaps the greatest utility of the atlas will come from comparisons to similarly rich profiles of additional lungs to parse normal variation from technical noise, from other life stages to elucidate how each cell type develops and ages, and to profiles of diseased lung tissue to define which and how cells are molecularly altered in disease^54, 55^. Indeed, this atlas should speed the molecular transition in pathology because now almost every lung cell type can be distinguished by the optimal set of sensitive and specific markers. And, with the progress in spatial transcriptomics^46–48^, it may soon be possible to obtain full expression profiles of all cells in situ, which would provide context for the individual cell variations seen in scRNAseq data and potentially reveal transient and specialized cell interactions and the dynamics of disease.

## Methods

### Human lung tissue and peripheral blood

Freshly resected lung tissue was procured intraoperatively from patients undergoing lobectomy for focal lung tumors. Normal lung tissues (∼5 cm^3^) were obtained from uninvolved regions and annotated for the specific lung lobe and location along the airway or periphery. Pathological evaluation (by G.B.) confirmed normal histology of the profiled regions, except for areas of very mild emphysema in Patient 1. Patient 1 was a 75 year-old male with a remote history of smoking, diagnosed with early stage adenocarcinoma who underwent left upper lobe (LUL) lobectomy; two blocks of normal tissue were obtained from lung periphery (“Distal 1a and 1b”). Patient 2 was a 46 year-old male, non-smoker with a right middle lobe (RML) endobronchial carcinoid, who underwent surgical resection of the right upper and middle lobes; two blocks of tissue were selected from mid-bronchial region (“Medial 2”) and periphery (“Distal 2”) of right upper lobe (RUL). Patient 3 was a 51 year-old female, non-smoker with a LLL endobronchial typical carcinoid, who underwent LLL lobectomy; three tissue blocks were resected from the bronchus (“Proximal 3”), mid-bronchial (“Medial 2”), and periphery (“Distal 3”) of the LLL. All tissues were received and immediately placed in cold phosphate buffered saline (PBS) and transported on ice directly to the research lab for single cell dissociation procedures. Peripheral blood was collected from patients 1 and 3 in EDTA tubes. For bulk RNAseq of canonical immune populations, whole blood from healthy human donors was obtained from AllCells Inc in EDTA tubes. Patient tissues were obtained under a protocol approved by Stanford University’s Human Subjects Research Compliance Office (IRB 15166) and informed consent was obtained from each patient prior to surgery.

### Mouse lung tissue

Lung tissue for Tabula Muris Senis^56^ was obtained as previously described. We obtained additional tissue from two mice expressing Cre recombinase and two expressing estrogen-inducible Cre recombinase (Cre-ERT2) for conditional cell-specific labeling *in vivo* with the gene-targeted alleles FVB-*Tbx4-LME-Cre*^57, 58^ (lung stroma) and B6.129-*Axin2-Cre-ERT2*^57^, respectively. Cre-dependent reporter alleles *Rosa26ZsGreen1*, which expresses cytosolic ZsGreen1 following Cre-mediated recombination, and *Rosa26mTmG*, which expresses membrane-targeted GFP (mGFP) following recombination and membrane-targeted tdTomato (mTomato) in all other tissues, were used to label cells expressing *Tbx4* and *Axin2*, respectively^59, 60^. Induction of the *Axin2-Cre-ERT2* allele was done by intraperitoneal injection of tamoxifen (3 mg) once a day for three days as described^38^. All mouse experiments were approved by the Institutional Animal Care and Use Committee at Stanford University (Protocol 9780).

### Isolation of lung and blood cells

Individual human lung samples were dissected, minced, and placed in digestion media (400 µg/ml Liberase DL (Sigma 5466202001) and 100 µg/ml elastase (Worthington LS006365) in RPMI (Gibco 72400120) in a gentleMACS c-tube (Miltenyi 130-096-334). Samples were partially dissociated by running ‘m_lung_01’ on a gentleMACS Dissociator (Miltenyi 130-093-235), incubated on a Nutator at 37°C for 30 minutes, and then dispersed to a single cell suspension by running ‘m_lung_02’. Processing buffer (5% fetal bovine serum in PBS) and DNAse I (100 µg/ml, Worthington LS006344) were then added and the samples rocked at 37°C for 5 minutes. Samples were then placed at 4°C for the remainder of the protocol. Cells were filtered through a 100 µm filter, pelleted (300 x g, 5 minutes, 4°C), and resuspended in ACK red blood cell lysis buffer (Gibco A1049201) for 3 minutes, after which the buffer was inactivated by adding excess processing buffer. Cells were then filtered through a 70 µm strainer (Fisherbrand 22363548), pelleted again (300 x g, 5 minutes, 4°C), and resuspended in magnetic activated cell sorting (MACS) buffer (0.5% BSA, 2 mM EDTA in PBS) with Human FcR Blocking Reagent (Miltenyi 130-059-901) to block non-specific binding of antibodies (see below).

Immune cells, including granulocytes, were isolated from peripheral blood using a high density ficoll gradient^61^. Briefly, peripheral blood was diluted 10-fold with FACS buffer (2% FBS in PBS), carefully layered on an RT Ficoll gradient (Sigma HISTOPAQUE®-1119), and centrifuged at 400 x g for 30 minutes at room temperature. The buffy coat was carefully removed, diluted 5-fold with FACS buffer, pelleted (300 x g, 5 minutes, 4°C), and incubated in ice cold FACS buffer containing DNAse I (Worthington LS006344) for 10 minutes at 4°C. Clumps were separated by gentle pipetting to create a single cell suspension.

Mouse lung samples were processed into single cell suspensions as previously described^19^. Briefly, each lung was dissected, minced, and placed in gentleMACS c-tubes (Miltenyi 130-096-334) with digestion buffer (400 µg/ml Liberase DL (Sigma 5466202001) in RPMI (Gibco 72400120)). The minced tissue was partially dissociated by running ‘m_lung_01’ on a gentleMACS Dissociator (Miltenyi 130-093-235), incubated at 37°C on a nutator for 30 minutes, completely dissociated on a gentleMACS by running ‘m_lung_02’, and kept at 4°C or on ice for the remainder of the protocol. Cells were washed with 5% FBS in PBS, centrifuged at 300 x g for 5 minutes, resuspended in 5% FBS in PBS, filtered through a 70 μm strainer (Fisherbrand 22363548), and centrifuged again and resuspended in FACS buffer (2% FBS in PBS).

### Magnetic separation of lung tissue compartments

Immune and endothelial cells were overrepresented in our previous mouse single cell suspensions. To partially deplete these populations in our human samples, we stained cells isolated from lung with MACS microbeads conjugated to CD31 and CD45 (Miltenyi 130-045-801, 130-091-935) then passed them through an LS MACS column (Miltenyi, 130-042-401) on a MidiMACS Separator magnet (Miltenyi, 130-042-302). Cells retained on the column were designated “immune and endothelial enriched.” The flow through cells were then split, with 80% immunostained for FACS (see below) and the remaining 20% stained with EPCAM microbeads (Miltenyi 130-061-101). EPCAM stained cells were passed through another LS column. Cells retained on the column were labeled “epithelial enriched”, and cells that flowed through were designated “stromal”.

### Flow cytometry and cell sorting

Lysis plates for single cell mRNA sequencing were prepared as previous described^19^. 96-well lysis plates were used for cells from the blood and mouse samples and contained 4 μL of lysis buffer instead of 0.4 μL.

Following negative selection against immune and endothelial cells by MACS, the remaining human lung cells were incubated with FcR Block (Becton Dickinson (BD) 564219) for 5 minutes and stained with directly conjugated anti-human CD45 (Biolegend 304006) and EPCAM (eBioscience 25-9326-42) antibodies on a Nutator for 30 minutes. Cells were then pelleted (300 x g, 5 minutes, 4°C), washed with FACS buffer three times, then incubated with cell viability marker Sytox blue (1:3000, ThermoFisher S34857) and loaded onto a Sony SH800S cell sorter. Living single cells (Sytox blue-negative) were sorted into lysis plates based on three gates: EPCAM^+^CD45^-^ (designated “epithelial”), EPCAM^-^CD45^+^ (designated “immune”), and EPCAM^-^ CD45^-^ (designated “endothelial or stromal”).

Immune cells from subject matched blood were incubated with FcR Block and Brilliant Violet buffer (BD 563794) for 20 minutes and then stained with directly conjugated anti-human CD3 (BD 563548), CD4 (BD 340443), CD8 (BD 340692), CD14 (BD 557831), CD19 (Biolegend 302234), CD47 (BD 563761), CD56 (BD 555516), and CD235a (BD 559944) antibodies for 30 minutes. Cells were pelleted (300 x g, 5 minutes, 4°C), washed with FACS buffer twice, and then incubated with the viability marker propidium iodide and loaded onto a BD FACSAria II cell sorter. Living (propidium iodide-negative) single, non-red blood (CD235a^-^) cells were sorted into lysis plates along with specific immune populations: B cells (CD19^+^CD3^-^), CD8+ T cells (CD8^+^), CD4^+^ T cells (CD4^+^), NK cells (CD19^-^CD3^-^CD56^+^CD14^-^), classical monocytes (CD19^-^ CD3^-^CD56^-^CD14^+^). After sorting, plates were quickly sealed, vortexed, spun down for 1 minute at 1000 x g, snap frozen on dry ice, and stored at -80 until cDNA synthesis.

Mouse cells were incubated with the viability marker DAPI and loaded onto a BD Influx cell sorter. Living (DAPI-negative) single cells were sorted into lysis plates based on presence or absence of the fluorescent lineage label (mEGFP for *Axin2-Cre-ERT2*, ZsGreen1 for *Tbx4-LME-Cre*).

Immune cells for bulk mRNA sequencing were incubated with Fc Block for 20 minutes and then stained with one of six panels of directly conjugated antibodies for 30 minutes: anti-human CD16 (BD 558122), CD123 (BD 560826), CCR3 (R&D FAB155F), ITGB7 (BD 551082), CD3 (BD 555341), CD14 (Invitrogen MHCD1406), CD19 (BD 555414), and CD56 (BD 555517) (“basophils, neutrophils and eosinophils”); anti-human CD16 (BD 558122), CD14 (BD 347497), CD4 (BD 340443), CD3 (BD 555341), CD8 (BD 555368), CD19 (BD 555414), and CD56 (BD 555517) (“classical and nonclassical monocytes”); anti-human CD16 (BD 558122), CD1c (Miltenyi Biotec 130-098-007), CD11c (BD 340544), CCR3 (R&D FAB155F), CD123 (BD 560826), HLA-DR (BD 335796), CD3 (BD 555341), CD4 (BD 555348), CD8 (BD 555368), CD14 (Invitrogen MHCD1406), CD19 (BD 555414), and CD56 (BD 555517) (“pDCs, mDCs, CD16^+^ DCs”); anti-human IgM/IgD (BD 555778), CD19 (BD 557835), CD27 (BD 558664), CD20 (BD 335794), CD3 (BD 555341), CD4 (BD 555348), CD14 (Invitrogen MHCD1406), and CD56 (BD 555517) (“B cells”); anti-human CD16 (BD 558122), CD57 (BD 347393), CD56 (BD 557747), CD3 (BD 555341), CD4 (BD 555348), CD14 (Invitrogen MHCD1406), and CD19 (BD 555414) (“NK cells”); and anti-human CD45RA (Biolegend 304118), CCR7 (R&D FAB197F), CD62L (BD 555544), CD45RO (BD Pharmingen 560608), CD4 (BD 340443), CD8 (BD 340584), CD11b (BD 555389), CD14 (Invitrogen MHCD1406), CD19 (BD 555414), CD56 (BD 555517) (“T cells”). Cells were washed with FACS buffer twice, incubated with the viability marker propidium iodide and loaded onto a BD FACSAria II cell sorter. 40,000 cells from 21 canonical immune populations (Table S6) were sorted in duplicate into Trizol LS (Invitrogen 10296010). After sorting, all plates and samples were quickly sealed, vortexed, spun down for 1 minute at 1000 x g and then snap frozen on dry ice and stored at -80 until cDNA synthesis.

### Single cell mRNA sequencing

mRNA from single cells sorted from human and mouse lungs and human blood into lysis plates was reverse transcribed to complementary DNA (cDNA) and amplified as previously described^19^. Illumina sequencing libraries for cDNA from single cells were prepared as previously described^19^. Briefly, cDNA libraries were prepared using the Nextera XT Library Sample Preparation kit (Illumina, FC-131-1096). Nextera tagmentation DNA buffer (Illumina) and Tn5 enzyme (Illumina) were added, and the sample was incubated at 55°C for 10 minutes. The reaction was neutralized by adding “Neutralize Tagment Buffer” (Illumina) and centrifuging at room temperature at 3,220 x g for 5 minutes. Mouse samples were then indexed via PCR by adding i5 indexing primer, i7 indexing primer, and Nextera NPM mix (Illumina). Human samples were similarly indexed via PCR using custom, dual-unique indexing primers (IDT)^19^.

Following library preparation, wells of each library plate were pooled using a Mosquito liquid handler (TTP Labtech), then purified twice using 0.7x AMPure beads (Fisher A63881). Library pool quality was assessed by capillary electrophoresis on a Tapestation system (Agilent) with either a high sensitivity or normal D5000 ScreenTape assay kit (Agilent) or Fragment analyzer (AATI), and library cDNA concentrations were quantified by qPCR (Kapa Biosystems KK4923) on a CFX96 Touch Real-Time PCR Detection System (Biorad). Plate pools were normalized and combined equally to make each sequencing sample pool. A PhiX control library was spiked in at 1% before sequencing. Human libraries were sequenced on a NovaSeq 6000 (Illumina) and mouse libraries on a NextSeq 500 (Illumina).

Cells isolated from each compartment (“immune and endothelial enriched”, “epithelial enriched”, “stromal”) and subject blood were captured in droplet emulsions using a Chromium Single-Cell instrument (10x Genomics) and libraries were prepared using the 10x Genomics 3’ Single Cell V2 protocol as previously described^19^. All 10x libraries were pooled and sequenced on a NovaSeq 6000 (Illumina).

### Immune cell bulk mRNA sequencing

Total RNA from bulk-sorted canonical immune populations was reverse transcribed to cDNA, amplified, and prepared as sequencing libraries as previously described^61^. Libraries were sequenced on a NextSeq 500 (Illumina).

### Immunohistochemistry

Mouse and human lungs were collected as previously described^38, 62^. After inflation, lungs were removed en bloc, fixed in 4% paraformaldehyde (PFA) overnight at 4°C with gentle rocking, then cryo-embedded in Optimal Cutting Temperature compound (OCT, Sakura) and sectioned using a cryostat (Leica) onto Superfrost Plus Microscope Slides (Fisherbrand). Immunohistochemistry was performed using primary antibodies raised against the following antigens and used at the indicated dilutions to stain slides overnight at 4°C: pro-SftpC (rabbit, Chemicon AB3786, 1:250 dilution), E-cadherin (rat, Life Technologies 131900 clone ECCD-2, 1:100), Mucin 1 (Muc1, hamster, Thermo Scientific HM1630, clone MH1, 1:250), Ki67 (rat, DAKO M7249, clone TEC-3, 1:100), Carbonic Anhydrase 2 (rabbit, Abcam EPR19839, 1:100). Primary antibodies were detected with Alexa Fluor-conjugated secondary antibodies (Jackson ImmunoResearch) unless otherwise noted, then mounted in Vectashield containing DAPI (5 ug/ml, Vector labs). Images were acquired with a laser scanning confocal fluorescence microscope (Zeiss LSM780) and processed with ImageJ and Imaris (version 9.2.0, Oxford Instruments). All immunostains were performed on at least 2 human subjects distinct from the donors used for sequencing, and quantifications were based on at least 10 fields of view in each.

### Single molecule *in situ* hybridization

Samples were fixed in either 10% neutral buffered formalin, dehydrated with ethanol and embedded in paraffin wax or fixed in 4% paraformaldehyde and embedded in OCT compound. Sections from paraffin (5 µm) and OCT (20 µm) blocks were processed using standard pre-treatment conditions for each per the RNAscope multiplex fluorescent reagent kit version 2 (Advanced Cell Diagnostics) assay protocol. TSA-plus fluorescein, Cy3 and Cy5 fluorophores were used at 1:500 dilution. Micrographs were acquired with a laser scanning confocal fluorescence microscope (Zeiss LSM780) and processed with ImageJ and Imaris (version 9.2.0, Oxford Instruments). All smFISH experiments were performed on at least 2 human subjects distinct from the donors used for sequencing, and quantifications were based on at least 10 fields of view for each. Fields of view were scored manually, calling a cell positive for each gene probed if its nucleus had >3 associated expression puncta. Proprietary (Advanced Cell Diagnostics) probes used were: KRT5 (547901-C2), SERPINB3 (828601-C3), SFTPC (452561-C2), WIF1 (429391), CLDN5 (517141-C2, 517141-C3), MYC (311761-C3), ACKR1 (525131, 525131-C2), COL1A2 (432721), GPC3 (418091-C2), SERPINF1 (564391-C3), C20rf85 (560841-C3), DHRS9 (467261), GJA5 (471431), CCL21 (474371-C2), COX4I2 (570351-C3), APOE (433091-C2), ACGT2 (828611-C2), ASPN (404481), IGSF21 (572181-C3), GPR34 (521021), EREG (313081), GPR183 (458801-C2), TREM2 (420491-C3), CHI3L1 (408121), MYRF (499261), AGER (470121-C3), TBX5 (564041), KCNK3 (536851), ACVRL1 (559221), SERPINA1 (435441), HHIP (464811), Slc7a10 (497081-C2), Fgfr4 (443511), Pi16 (451311-C2), Serpinf1 (310731), Hhip (448441-C3), Sftpc (314101-C2), Nkx2-1 (434721-C3), and Myrf (524061).

## Bioinformatic Methods

### Read alignments and quality control

Reads from single cells isolated using 10x chromium were demultiplexed and then aligned to the GRCh38.p12 human reference (from 10x Genomics) using Cell Ranger (version 2.0, 10x Genomics). Cells with fewer than 500 genes detected or 1000 unique molecular identifiers (UMIs) were excluded from further analyses.

Reads from single cells isolated by flow cytometry were demultiplexed using bcl2fastq (version 2.19.0.316, Illumina), pruned for low nucleotide quality scores and adapter sequences using skewer (version 0.2.2), and aligned to either (depending on organism) the GRCh38.p12 human reference genome with both the gencode-vH29 and NCBI-108 annotations or the GRCm38.p6 mouse reference genome with the NCBI-106 annotation (with fluorescent genes mEGFP, tdTomato, and ZsGreen1 supplemented) using STAR (version 2.6.1d) in two-pass mapping mode, in which the first pass identifies novel splice junctions and the second pass aligns reads after rebuilding the genome index with the novel junctions. The number of reads mapping to each annotated gene were calculated by STAR during the second pass alignment, and cells with fewer than 500 genes detected or 50,000 mapped reads were excluded from later analyses. Reads from mRNA sequencing of canonical immune populations were demultiplexed, aligned, and quantified using the same pipeline.

### Cell clustering, doublet calling, and annotation

Expression profiles of cells from different subjects and different capture approaches (10x and SS2) were clustered separately using the R software package Seurat (version 2.3)^63^. Briefly, counts (SS2) and UMIs (10x) were normalized across cells, scaled per million (SS2) or per 10,000 (10x), and converted to log scale using the ‘NormalizeData’ function. These values were converted to z-scores using the ‘ScaleData’ command and highly variable genes were selected with the ‘FindVariableGenes’ function with a dispersion cutoff of 0.5. Principle components were calculated for these selected genes and then projected onto all other genes with the ‘RunPCA’ and ‘ProjectPCA’ commands. Clusters of similar cells were detected using the Louvain method for community detection including only biologically meaningful principle components (see below) to construct the shared nearest neighbor map and an empirically set resolution, as implemented in the ‘FindClusters’ function.

Clusters were grouped and separated based on expression of tissue compartment markers (e.g. *EPCAM*, *CLDN5*, *COL1A2*, and *PTPRC*) using the ‘SubsetData’ command and the same procedure (from ‘ScaleData’ onwards) was applied iteratively to each tissue compartment until the markers enriched in identified clusters, identified using the ‘MAST’ statistical framework^64^ implemented in the ‘FindMarkers’ command, were no longer biologically meaningful (e.g. clusters distinguished by dissociation-induced genes^42^, ribosomal genes, mitochondrial genes, or ambient RNA released by abundant cells such as RBCs^43^). Doublets were identified by searching for cells with substantial and coherent expression profiles from two or more tissue compartments and/or cell types.

To assign clusters identities, we first compiled a list of all established lung cell types, their abundances, their classical markers, and any RNA markers (when available) (Table S1). RNA markers for canonical immune populations were obtained using the ‘MAST’ statistical framework (Table S4). Clusters were assigned a canonical identity based on enriched expression of these marker genes. There were no clusters that lacked expression of canonical marker genes. When two or more clusters were assigned the same identity, we first determined whether their tissue locations differed substantially (e.g. proximal versus distal, alveolar versus adventitial) and prepended these locations when applicable. When both clusters localized to the same tissue region (e.g. capillary endothelial cells or AT2 cells), we next compared their differentially expressed genes head-to-head to identify differences in molecular functions. These functional differences were also prepended, when applicable (e.g. Signaling AT2 versus AT2, Proliferative Basal versus Basal). If the clusters could not be resolved by location or function, we prepended a representative marker gene to their “canonical” identity (e.g. IGSF21+ Dendritic, EREG+ Dendritic, and TREM2+ Dendritic). Cells from different subjects with the same annotation were merged into a single group for all downstream analyses.

Approximately 35,000 mouse lung and blood cell expression profiles by SS2 and 10x from Tabula Muris Senis^19^ were combined with 522 cells isolated from *Axin2-Cre-ERT2 >Rosa26mTmG* (A.N.N.) and *Tbx4-LME-Cre > Rosa26ZsGreen1* (K.J.T.) mice and amplified by SS2. Cells were stratified by technology (10x versus SS2), re-clustered and re-annotated using the strategy described above for human lung cells.

### Re-annotation of existing human lung single cell RNA sequencing datasets

UMI tables were obtained from the Gene Expression Omnibus (GSE122960 for Reyfman et al, GSE130148 for Braga et al), clustered, and annotated using the strategy described above. New annotations for each cell are available on GitHub (see below).

### Identification of proliferation signature

Expression profiles from matched proliferating and quiescent cell types were compared head-to-head using the ‘MAST’ statistical framework implemented in the ‘FindMarkers’ command in Seurat. Differentially-expressed genes common in each proliferating cell type were converted to z-scores using the ‘ScaleData’ command in Seurat, and summed to create a “proliferation score” for each cell.

### Identification of immune egression signatures

Blood and tissue expression profiles for each immune cell type were compared head-to-head using the ‘MAST’ statistical framework implemented in the ‘FindMarkers’ command in Seurat. Differentially-expressed genes common in each subject were screened for dissociation artifact and contamination by red blood cells. Genes specific to tissue immune cells were binned based on their breadth of expression (lymphocyte, myeloid, or both), converted to z-scores using the ‘ScaleData’ command in Seurat, and summed to create an “egression score” for each cell.

### Identification of enriched marker genes, transcription factors, and disease genes

Differentially-expressed genes for each annotated cell type relative to the other cells within its tissue compartment were identified using the ‘FindMarkers’ command in Seurat with the ‘MAST’ statistical framework. To obtain the most sensitive and specific markers for each cell type, we ranked enriched genes, with a p-value less than 10^-5^ and a sensitivity greater than 0.4, by their Mathews Correlation Coefficients (MCCs). To measure the utility of using multiple markers in assigning cell identities, we calculated MCC scores for all possible combinations of each cell type’s top five marker genes.

Enriched genes were annotated as transcription factors or genes associated with pulmonary pathology based on lists compiled from The Animal Transcription Factor Database^65^, The Online Mendelian Inheritance in Man Catalog (OMIM)^49^, and Genome Wide Association Studies (GWAS) obtained from the EMBL-EBI Catalog^50^ (EFO IDs 0000270, 0000341, 0000464, 0000571, 0000702, 0000707, 0000708, 0000768, 0001071, 0003060, 0003106, 0004244, 0004312, 0004313, 0004314, 0004647, 0004713, 0004806, 0004829, 0005220, 0005297, 0006505, 0006953, 0007627, 0007744, 0007944, 0008431, 0009369, 0009370; GO IDs 0031427, 0097366; Orphanet IDs 586 182098; log(p-value) < -20). Viral entry genes were obtained from Gene Ontology (GO:0046718) and then curated and associated with their cognate virus(es) based on literature citations available in our GitHub repository.

### Cellular interaction and hormone target mapping

Interactions between cell types were predicted using CellPhoneDB, as previously described^25^. For our targeted analyses, we curated the chemokine receptor-ligand interaction map and list of hormone receptors from an extensive literature search (available on GitHub, see below).

### Human and mouse gene alignment, cell type correlation, and gene expression comparisons

The gene expression matrices from our human SS2 cells and the Tabula Muris Senis SS2 cells, supplemented with the 522 mouse cells from *Axin2-CreER > mTmG* and *Tbx4-Cre > ZsGreen1* described above, were collapsed to HomologyIDs obtained from the Mouse Genome Informatics database to enable direct comparison. We obtained median expression profiles for each cell type and calculated pairwise Pearson correlation coefficients using the ‘cor’ function in R. We defined species-specific gene expression as those enriched 20-fold in either direction (mouse > human or human > mouse) with a p-value less than 10^-5^ (calculated by ‘MAST’ as above).

To compare the expression pattern of each gene across species we binarized genes as expressed (1) or not expressed (0) in each cell type. A cell type “expressed” a gene if the median of that gene’s non-zero expression values across the constituent cells was greater than the median of every non-zero expression value for all other genes plus/minus two standard deviations (varied in 0.25 increments) and if the percentage of cells within the cell type with non-zero expression values was greater than the median percent of non-zero expression values for all other genes plus/minus two standard decisions (varied in 0.25 increments). These cutoffs were varied independently to ensure genes were robustly categorized. We then ordered these gene vectors to match homologous cell types between species with at least five cells and combined them to a single vector for each gene (V = (a – b) + 2ab, where a is the ordered human vector and b is the ordered mouse vector) that indicated for each cell type whether: Both mouse and human expressed the gene (2), only human (1), only mouse (-1), or neither (0). We then classified genes by the following: Conserved if any element of V equaled 2 and all other elements equaled 0, Type 2 if any element equaled 2 and any other equaled 1 or -1, not expressed if all elements equaled 0, Type 3 if elements were both positive and negative, and Type 1 if elements were either positive or negative and 0.

### Data and code availability

Counts/UMI tables, cellular metadata, Seurat objects, and scanpy objects are available on Synapse (accession syn21041850). The data can be explored in a browser using cellxgene at https://hlca.ds.czbiohub.org/. Code for demultiplexing counts/UMI tables, clustering, annotation, and other downstream analyses are available on GitHub (https://github.com/krasnowlab/HLCA). Human sequencing data will be available by data access agreement on the European Genome-phenome Archive (EGA) upon publication of this manuscript. Mouse sequencing data will be available on the National Institute of Health’s Sequence Read Archive (SRA) also upon publication of this manuscript.

## Acknowledgements

We are grateful to the tissue donors and the clinical staff at Stanford Medical Center who made tissue collection possible, especially Jalen Benson and Emily Chen. We are especially grateful to Jim Spudich who spurred this study. We also thank the Stanford Shared FACS Facility for their expertise and sorting services, especially Dr. Lisa Nichols and Meredith Weglarz; members of Chan Zuckerberg Biohub and Quake Lab who supported this work, particularly Aaron McGeever, Dr. Brian Yu, Bob Jones, and Saroja Kolluru; Dr. Maya Kumar for discussions on annotation of stromal cells; and Maria Petersen for illustrating the lung schematic (Fig. 2b) and Dr. Camilla Kao for help with figure formatting. Some computing for this project was performed on the Sherlock cluster; we thank Stanford University and the Stanford Research Computing Center for providing computational resources and support that contributed to the results. We thank Jim Spudich and members of the Krasnow lab for valuable discussions and comments on the manuscript, and Alexander Lozano for discussions on bioinformatic analyses. This work was supported by funding from the Chan Zuckerberg Biohub (S.R.Q.) and the Howard Hughes Medical Institute (M.A.K.). K.J.T was supported by a Paul and Mildred Berg Stanford Graduate Fellowship. M.A.K. is an investigator of the Howard Hughes Medical Institute.

## Author Contributions

K.J.T., A.N.N., L.P., R.S., A.G., C.S.K., R.J.M., and M.A.K. conceived the project and designed the lung and blood cell isolation strategy, J.B.S. and C.S.K. designed clinical protocols, reviewed clinical histories and coordinated patient care teams to obtain profiled tissues, G.B. provided expert clinical evaluation and micrographs of donor tissue histology, K.J.T., A.N.N., R.S., and A.G. processed tissue to single cell suspensions, K.J.T., A.N.N., L.P. A.G., R.S., S.D.C. sorted cells for SS2, A.N.N., L.P., S.C., and R.V.S. prepared sequencing libraries, and K.J.T., R.V.S. and L.P. processed and aligned sequencing data. R.S., J.S., and Y.M. performed and supervised bulk mRNA sequencing on defined immune populations. K.J.T., A.N.N., R.S. A.G., and R.J.M. provided tissue expertise and annotated cell types. K.J.T., A.N.N., and M.A.K. designed and implemented bioinformatic methods and interpreted results. K.J.T., A.N.N., and A.G. performed follow up stains. M.A.K., S.R.Q., N.F.N., I.L.W., C.S.K., and R.J.M. supervised and supported the work. K.J.T., A.N.N., and M.A.K. wrote the manuscript, and all authors reviewed and edited the manuscript.

## Extended Data Figure Legends

**Extended Data Figure S1.**
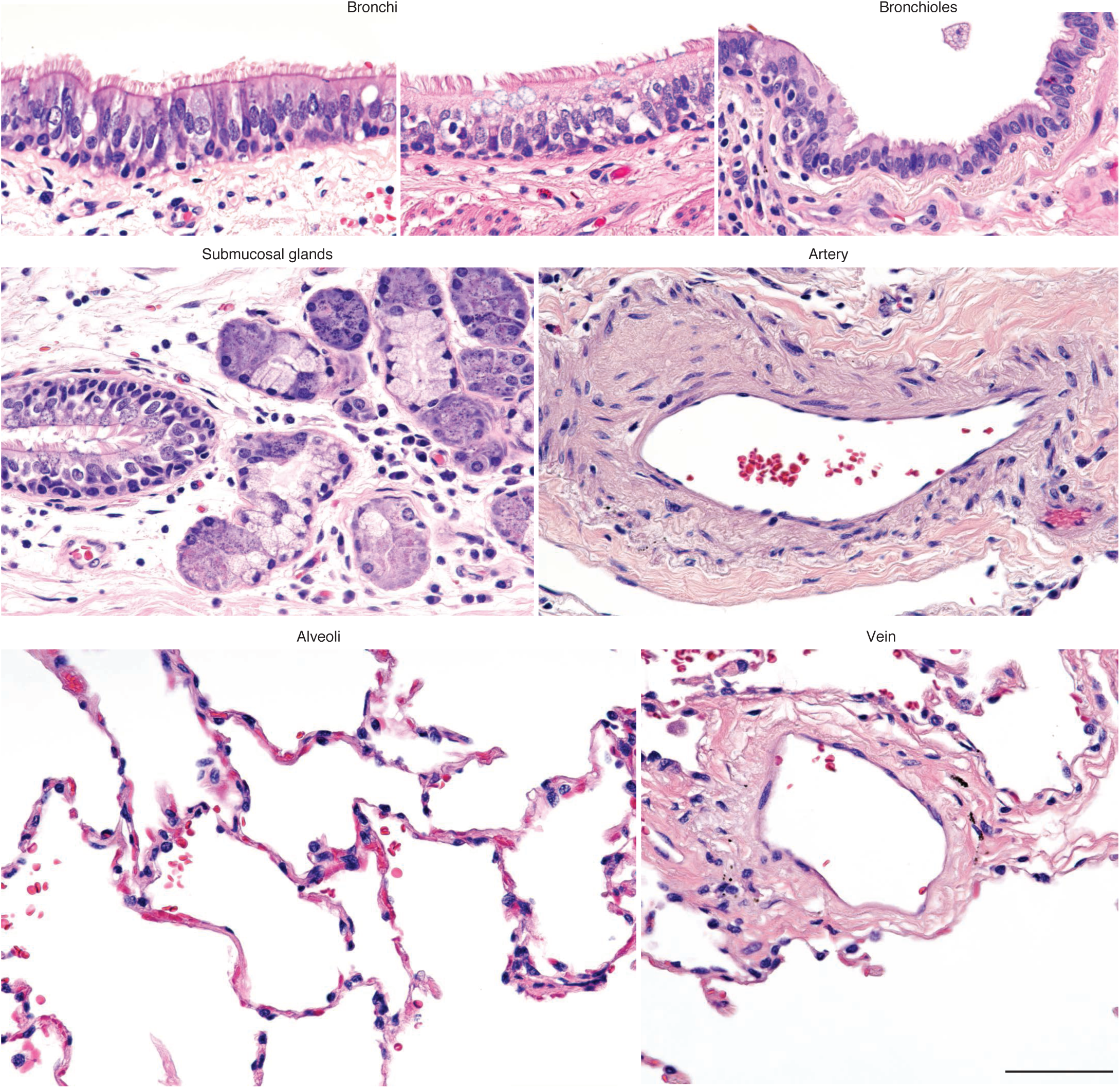
Histology of anatomical landmarks in donor lungs. Representative micrographs of donor lungs from formalin-fixed, paraffin-embedded (FFPE) sections stained with haematoxylin and eosin showing bronchi, bronchioles, submucosal glands, arteries, veins, and alveoli near regions used for single cell RNA sequencing. Scale bar, 100 µm.

**Extended Data Figure S2.**
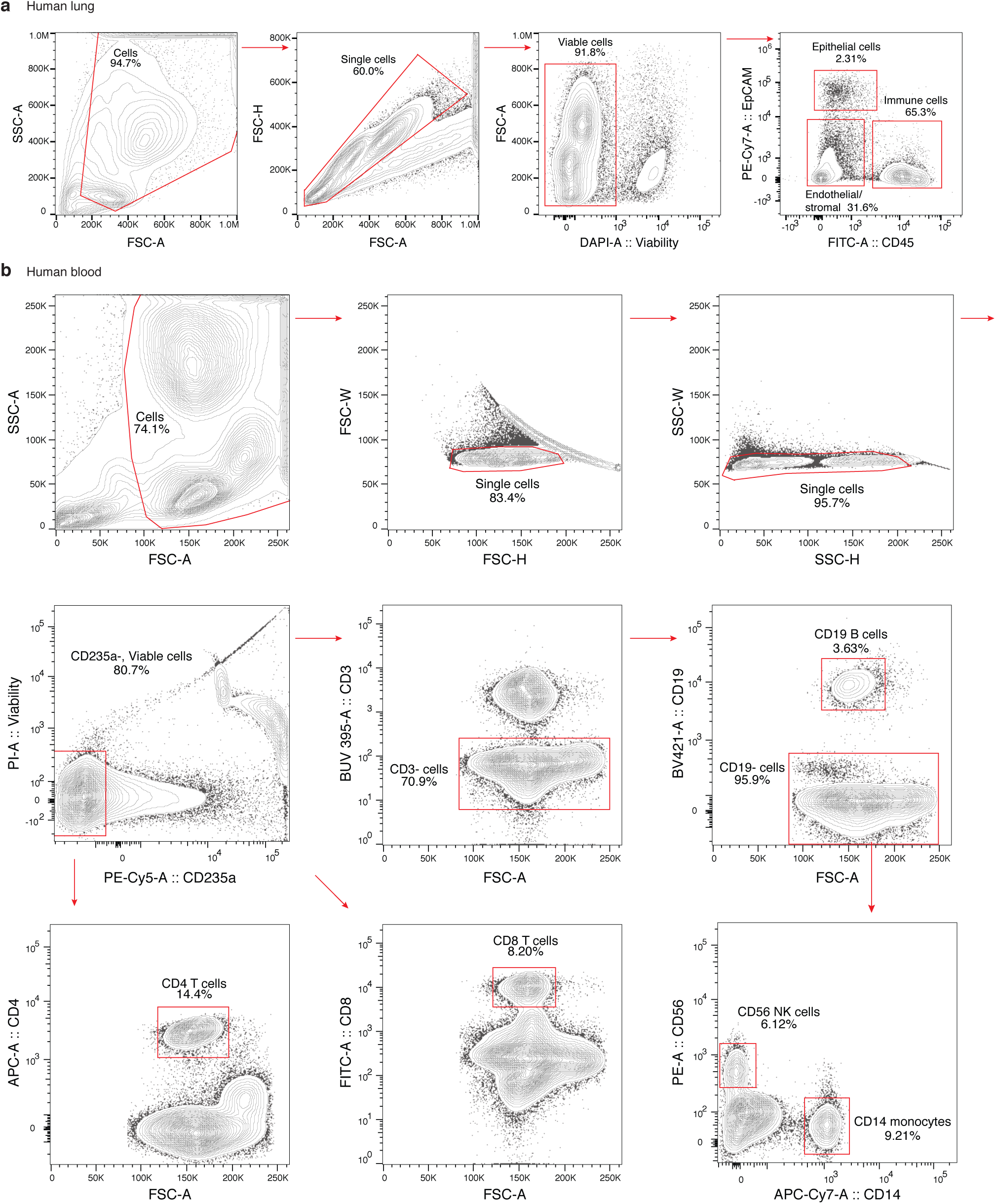
FACS gating strategies for human lung and peripheral blood cells. **a**, Sequential FACS data and sorting gates (red) for dissociated human lung cells from subject sample D1b (plate B001223) following MACS depletion of highly abundant immune (CD45^+^) and endothelial (CD31^+^) cells. The final sort (right) was of viable single cells from the lung epithelial (EPCAM^+^CD45^-^), immune (CD45^+^EPCAM^-^), and stromal/endothelial (EPCAM^-^ CD45^-^) compartments into 384-well plates for SS2 scRNAseq. **b,** Sequential FACS data and sorting gates (red) for white blood cells isolated on a Ficoll gradient of matched subject peripheral blood (subject 1, plate BP1). Viable, single CD235a^-^ (non-RBC) cells were captured without additional gating (panel 4), or further sorted as CD8 T (CD8^+^; panel 8), CD4 T (CD4^+^; panel 7), B (CD19^+^CD3^-^; panel 6), NK (CD19^-^CD3^-^CD56^+^CD14^-^; panel 9), or CD14^+^ monocytes (CD19^-^CD3^-^CD56^-^CD14^+^; panel 9) for SS2 scRNAseq. Contours, 5% increments in cell density.

**Extended Data Figure S3.**
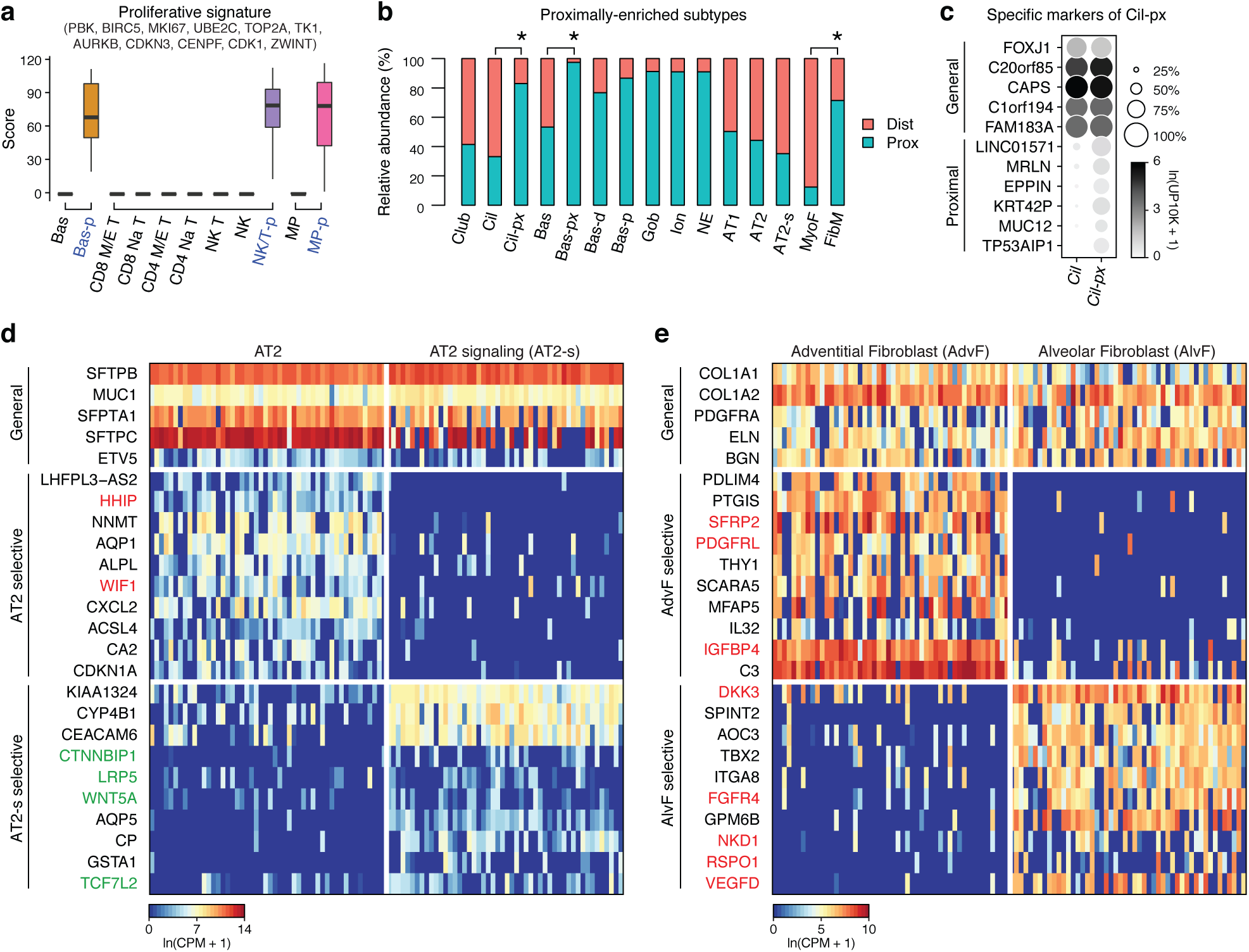
Expression and positional differences among identified subtypes of canonical lung populations. **a**, Proliferative signature (based on expression of indicated genes) of each cluster of basal cells, NK/T cells, and macrophages. Note three clusters with a high signature: basal-proliferative (Bas-p), NK/T-proliferative (NK/T-p), and macrophage-proliferative (MP-p) **b,** Relative abundance of epithelial and stromal cell types in scRNAseq analysis of human lung samples obtained from proximal (blue; samples P3) and distal (red; samples D1a, D1b, D2, D3) sites in the lung. In addition to the expected proximal enrichment of some airway cell types (goblet, ionocytes, neuroendocrine cells) and distal enrichment of alveolar cell types (AT1, AT2, AT2-s, myofibroblasts), note the three cell types (ciliated, cil; basal, bas; myofibroblasts, MyoF) with transcriptionally-distinct subsets that differ in their proximal-distal enrichment: ciliated (cil) vs. ciliated-proximal (cil-px), basal (bas) vs. basal-proximal (bas-px), myofibroblasts (MyoF) vs. fibromyocyte (FibM). (Note relative enrichment values should be considered provisional because enrichment values can be influenced by efficiency of harvesting during cell dissociation and isolation.) **c,** Dot plot of expression in ciliated cells (Cil) and proximal ciliated cells (Cil-px) of canonical (General) ciliated cell markers and Cil-px-specific (Proximal) markers. **d,** Heatmap of expression of representative general AT2 marker genes, AT2-selective and AT2-s selective marker genes indicated in AT2 (left panel) and (right panel) human lung cell clusters from the SS2 expression profiles. Note AT2 selective marker genes include negative regulators of the Hedgehog and Wnt signaling pathways (e.g., *HHIP* and *WIF1*, highlighted in red) and AT2-s selective markers include Wnt ligands, receptors, and transcription factors (e.g., *WNT5A*, *LRP5*, and *TFC7L2* highlighted in green). Values shown are ln(CPM+1) for 50 randomly-selected cells in each cluster from the SS2 analysis. **e,** Heatmap of expression of representative general, adventitial-selective, and alveolar-selective fibroblast markers in 50 randomly-selected cells in the adventitial fibroblast (left) and alveolar fibroblast (right) clusters from SS2 expression profiles. Note specialization in growth factors (AdvF: *PDGFRL*, *IGFBP4*, AlvF: *FGFR4*, *VEGFD*) and morphogen (AdvF: *SFRP2*, AlvF: *NKD1*, *DKK3*) signaling/regulation in red.

**Extended Data Figure S4.**
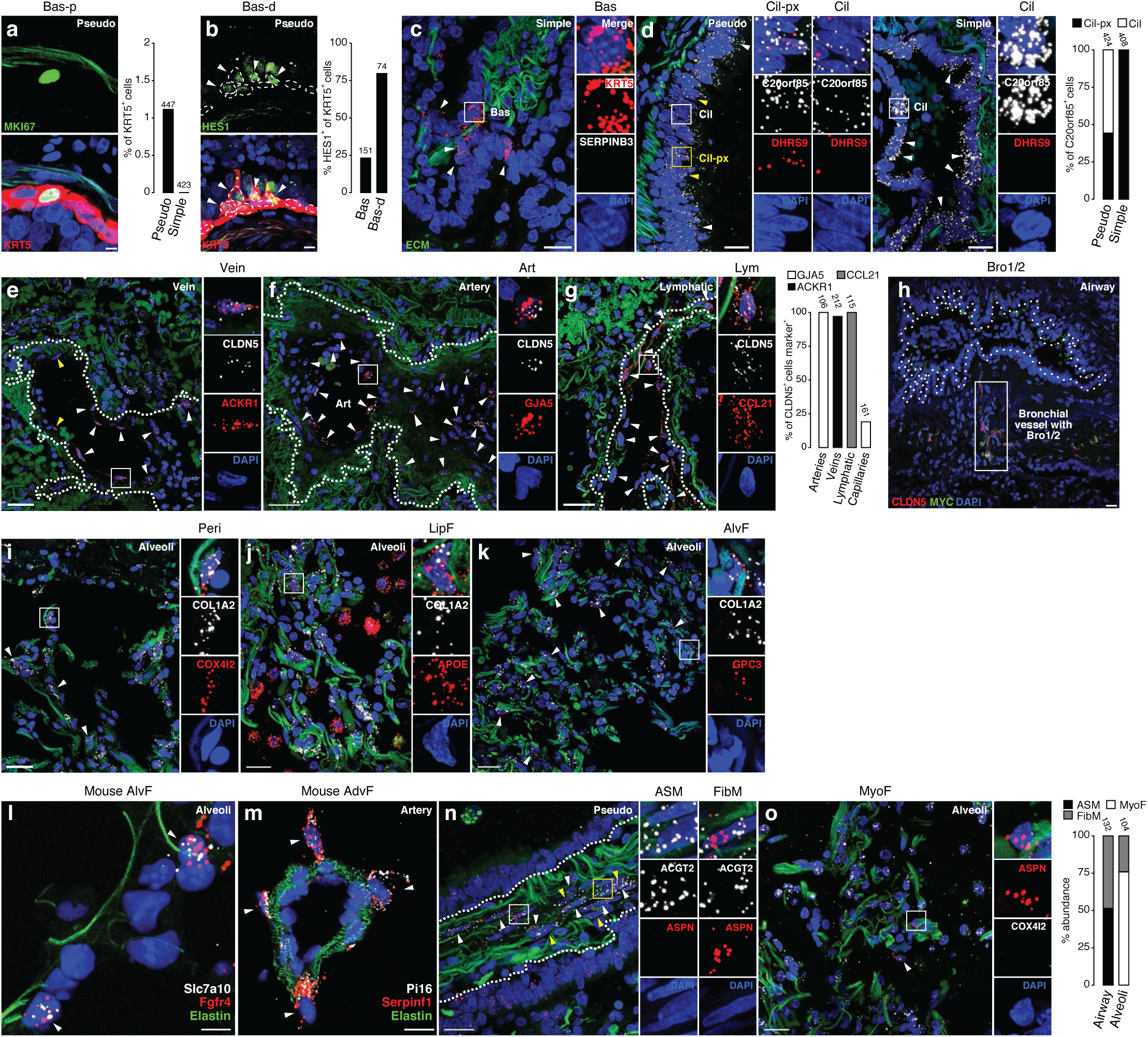
smFISH and immunostaining of cell markers to localize epithelial, endothelial, stromal clusters in the lung. **a,** Immunostaining of adult human pseudostratified airway for proliferation marker MKI67 (green) in basal cells (marked by KRT5, red) with DAPI (nuclear) counter stain (blue). Scale bars, 5 µm. Quantification shows abundance of proliferating (MKI67-expresing) basal cells (Bas-p) in pseudostratified and simple epithelial airways; n, KRT5+ cells scored in sections of two human lungs. **b,** Immunostaining of adult human pseudostratified airway for differentiation marker HES1 (green) in basal cells (marked by KRT5, red) with DAPI (nuclear) counter stain (blue). Scale bars, 10 µm. Note apical processes extending from HES1+ basal cells (arrowheads) indicating migration away from basal lamina as they differentiate. Other HES1^+^ cells have turned off basal marker KRT5. Dashed outlines, basal cell nuclei. Quantification shows fraction of basal cells (Bas, cuboidal KRT5+ cells on basement membrane) and Bas-d cells (KRT5+ cells with apical processes) that were HES1+. **c,** smFISH of adult human simple epithelial airway for general basal cell marker *KRT5* (red) and proximal basal cell marker *SERPINB3* with DAPI counter stain (blue) and ECM autofluorescence (green). *SERPINB3* is not detected, indicating only Bas cells but not Bas-px cells are present in simple airways (compare with proximal, pseudostratified airway in Fig. 2d). Scale bar, 10 µm. **d,** smFISH and quantification of human pseudostratified epithelial (left panel) and simple epithelial (right panel) airways for general ciliated marker *C20orf85* (white) and proximal ciliated marker *DHRS9* (red) with DAPI counterstain (blue) and ECM autofluorescence (green). Note restriction of Cil-px ciliated cells (expressing proximal marker) to pseudostratified airways. Scale bars, 10 µm. **e-g,** smFISH and quantification of vessel types indicated (dotted outlines) showing expression of vein marker *ACKR1* (red, panel e), artery marker *GJA5* (red, panel f), lymphatic marker *CCL21* (red, panel g), and general endothelial marker *CLDN5* with DAPI counter stain (blue) and ECM autofluorescence (green). Scale bars, 50 µm, 30 µm, and 40 µm. **h,** Micrograph (low magnification) of bronchial vessel (boxed region) in Fig. 2h showing vessel location near airway (dotted outline). **i-k,** smFISH and quantification of human alveolar region showing expression of pericyte marker *COX4I2* (red, panel i), lipofibroblast marker *APOE* (red, panel j), alveolar fibroblast marker *GPC3* (red, panel k), and general stromal marker *COL1A2* with DAPI counter stain (blue) and ECM autofluorescence (green). Note (panel j) the boxed *COL1A2* and *APOE* double-positive lipofibroblast (LipF) among single-positive *COL1A2* cells (other stromal cells) and *APOE* cells (macrophages); for quantification, see Fig. 2j. Scale bars, 20 µm. **l,** smFISH of alveolar region of adult mouse lung for alveolar fibroblast markers *Slc7a10* (white) and *Frfr4* (red) co-stained for alveolar entrance ring marker Elastin (green) with DAPI counterstain (blue). Scale bar, 5 µm. **m**, smFISH of mouse pulmonary artery for adventitial fibroblast markers *Pi16* (white) and *Serpinf1* (red) and co-stained for arterial marker Elastin (green) and counterstained with DAPI (blue). Note localization of the double-positive cells (adventitial fibroblasts, arrowheads) around artery. Scale bar, 10 µm. **n, o,** smFISH and quantification of human pseudostratified epithelial airway (n) and alveolar region (o) for airway smooth muscle (ASM) and fibrobmyocyte (FibM) markers *ACTG2* (white, n) and COX4I2 (white, o), fibromyocyte and myofibroblast (MyoF) marker *ASPN* (red), with DAPI counter stain (blue) and ECM autofluorescence (green). Note intermingled airway smooth muscle and fibromyocytes near a pseudostratified airway (dotted outline) in n, and *ASPN*^+^ *COX4I2*^-^ myofibroblasts (boxed and arrowhead) enriched in alveoli in o.

**Extended Data Figure S5.**
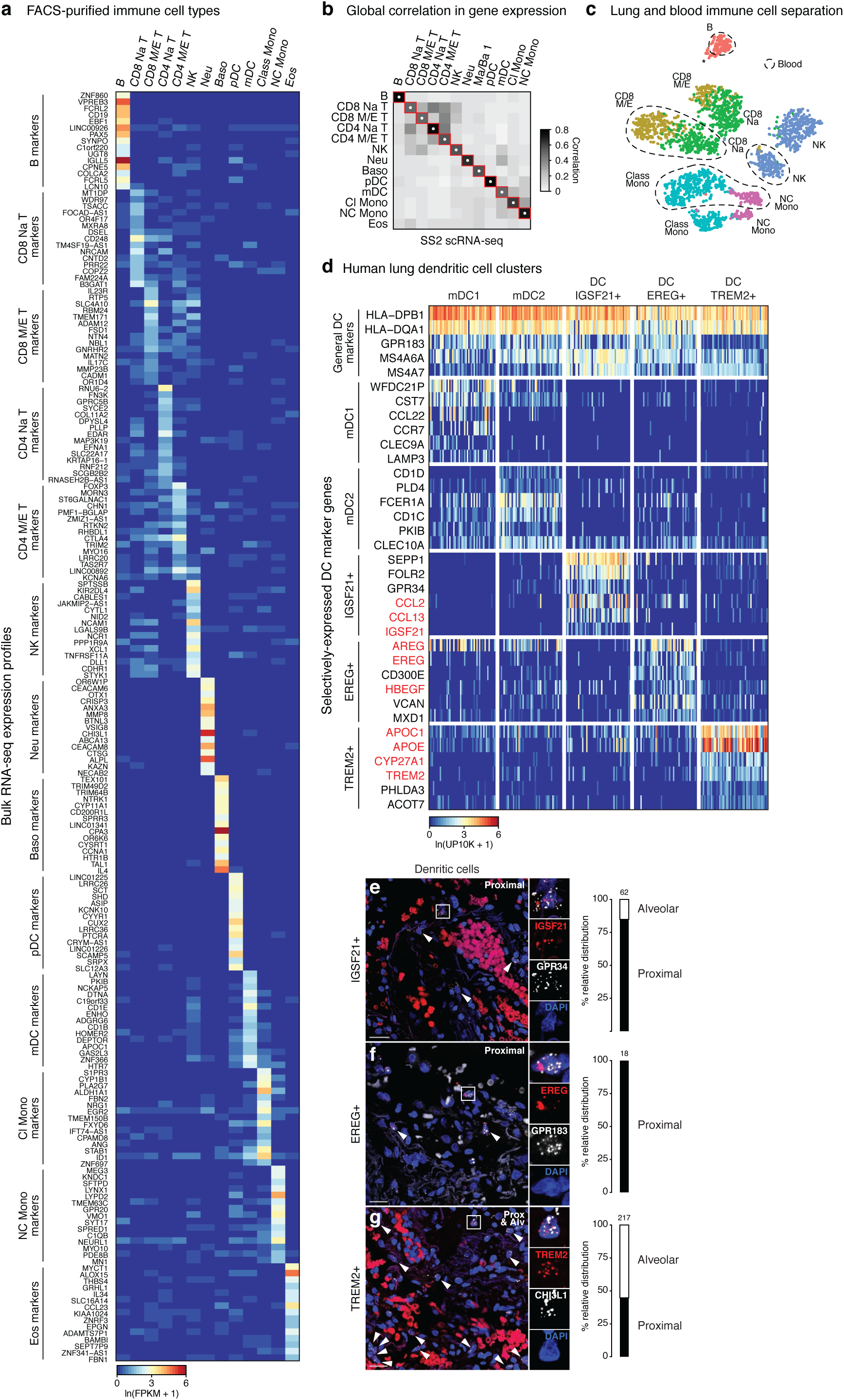
Selectively-expressed RNA markers of human immune cell types from bulk mRNA sequencing of FACS-purified immune cells. **a,** Heatmap of RNA expression of the most selectively-expressed genes from bulk mRNA sequencing of the indicated FACS-sorted immune populations (see Table S3). This dataset provided RNA markers for human immune cell populations that have been classically defined by their cell surface markers. **b,** Heatmap of pairwise correlation scores between the average expression profiles of the immune cell types indicated that were obtained from bulk mRNA sequencing (BulkSeq, panel a) to the average scRNAseq profiles of human blood immune cells in the SS2 dataset annotated by canonical markers and enriched RNA markers from the bulk RNA-seq analysis. The highest correlation in overall gene expression (white dot) of each annotated immune cell cluster in the SS2 dataset (columns) was to the bulk RNA-seq of the same FACS-purified immune population (rows), supporting the scRNAseq immune cluster annotations (red squares). **c,** tSNE of monocytes and B, T, and NK cells from patient 1 10x dataset. Note separate clusters of each immune cell type isolated from lung (no outline) and blood (dashed outline). Asterisk, small number of B cells isolated from the lung that cluster next to blood B cells. **d,** Heatmap of expression of dendritic cell marker genes in the scRNAseq profiles of the indicated dendritic cell clusters from human blood and lung 10x datasets. Note all clusters express general dendritic markers including antigen presentation machinery, but each has its own set of selectively-expressed markers. The red-highlighted markers that distinguish the three novel dendritic cell clusters we identified (IGSF21+, EREG+, TREM2+) suggest differential roles in asthma (IGSF21+), growth factor regulation (EREG+), and lipid handling (TREM2+). **e-g,** smFISH of adult human lung proximal and alveolar regions as indicated showing expression of IGSF21+ DC markers *GPR34* (white) and *IGSF21* (red) in e, EREG+ DC marker *EREG* (red) and general DC marker *GPR183* (white) in f, and TREM2+ DC markers *CHI3L1* (white) and *TREM2* (red) in g, all with DAPI (nuclear) counter stain (blue). (The non-punctate signal in red channel in panels e and g is erythrocyte autofluorescence.) Scale bars, 20 µm. Arrowheads, double positive cells. Quantification shows distribution of each dendritic type; note strong proximal enrichment of IGSF21+ and EREG+ dendritic cells.

**Extended Data Figure S6.**
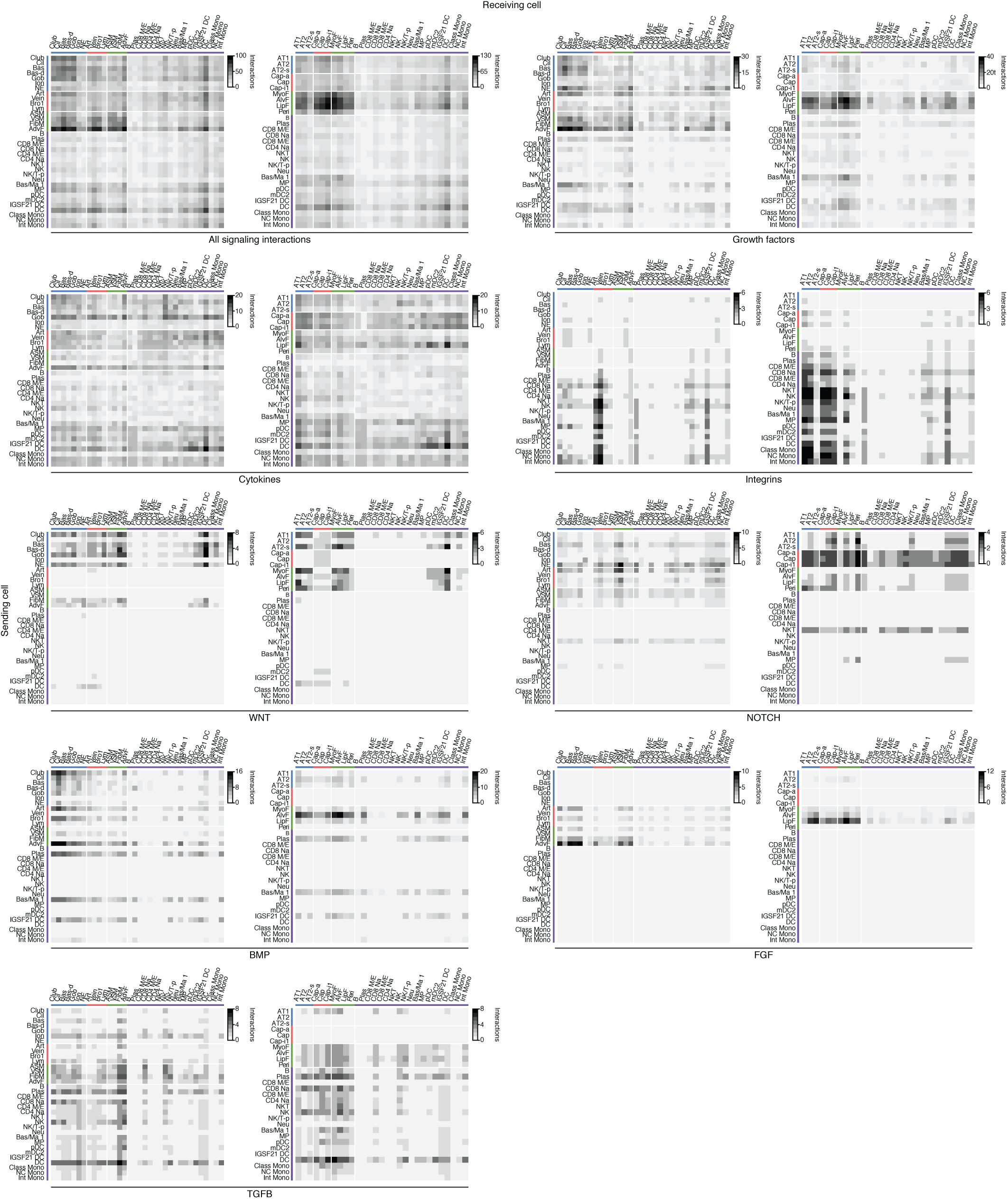
Global signaling interactions among human lung cell types inferred from expression patterns of ligands and cognate receptor genes. Heatmaps showing number of interactions predicted by CellPhoneDB software between human lung cell types located in proximal lung regions (left panel in each pair) and distal regions (right panel) based on expression patterns of ligand genes (“Sending cell”) and their cognate receptor genes (“Receiving cell”) in the SS2 dataset. The pair of heatmaps at upper left show values for all predicted signaling interactions (“All interactions”), and other pairs show values for the indicated types of signals (growth factors, cytokines, integrins, WNT, Notch, Bmp, FGF, and TFGB). Predicted interactions between cell types range from 0 (lymphocyte signaling to neutrophils) to 136 (AdvF signaling to Cap-i1). Note expected relationships such as immune cells expressing integrins to interact with endothelial cells and having higher levels of cytokine signaling relative to their global signaling, and unexpected relationships such as fibroblasts expressing majority of growth factors and lack of Notch signaling originating from immune cells.

**Extended Data Figure S7.**
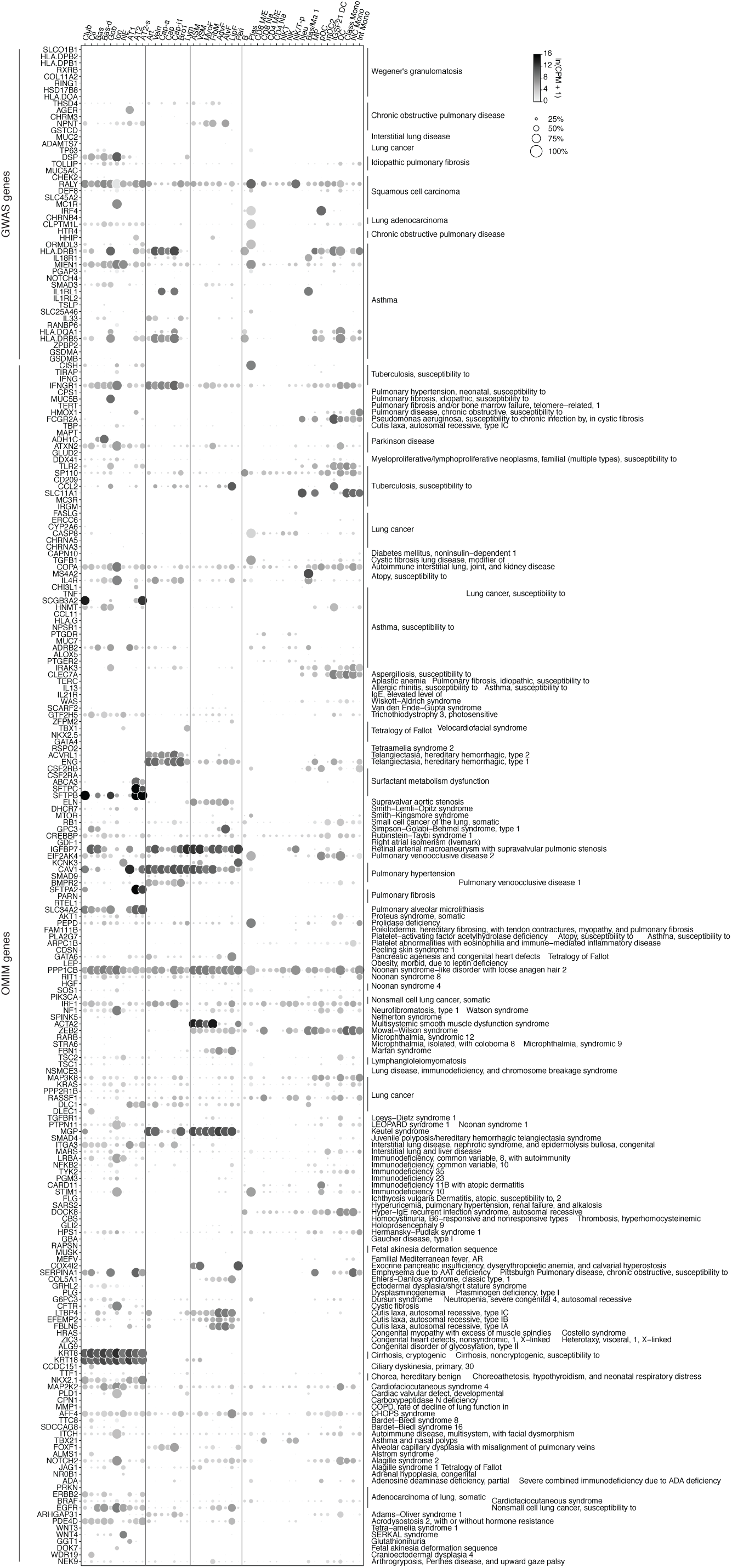
Lung cell expression patterns of genes implicated in lung disease. Dot plots of expression of 233 lung disease genes curated from Genomewide Association Studies (GWAS) and Online Mendelian Inheritance in Man (OMIM).

**Extended Data Figure S8.**
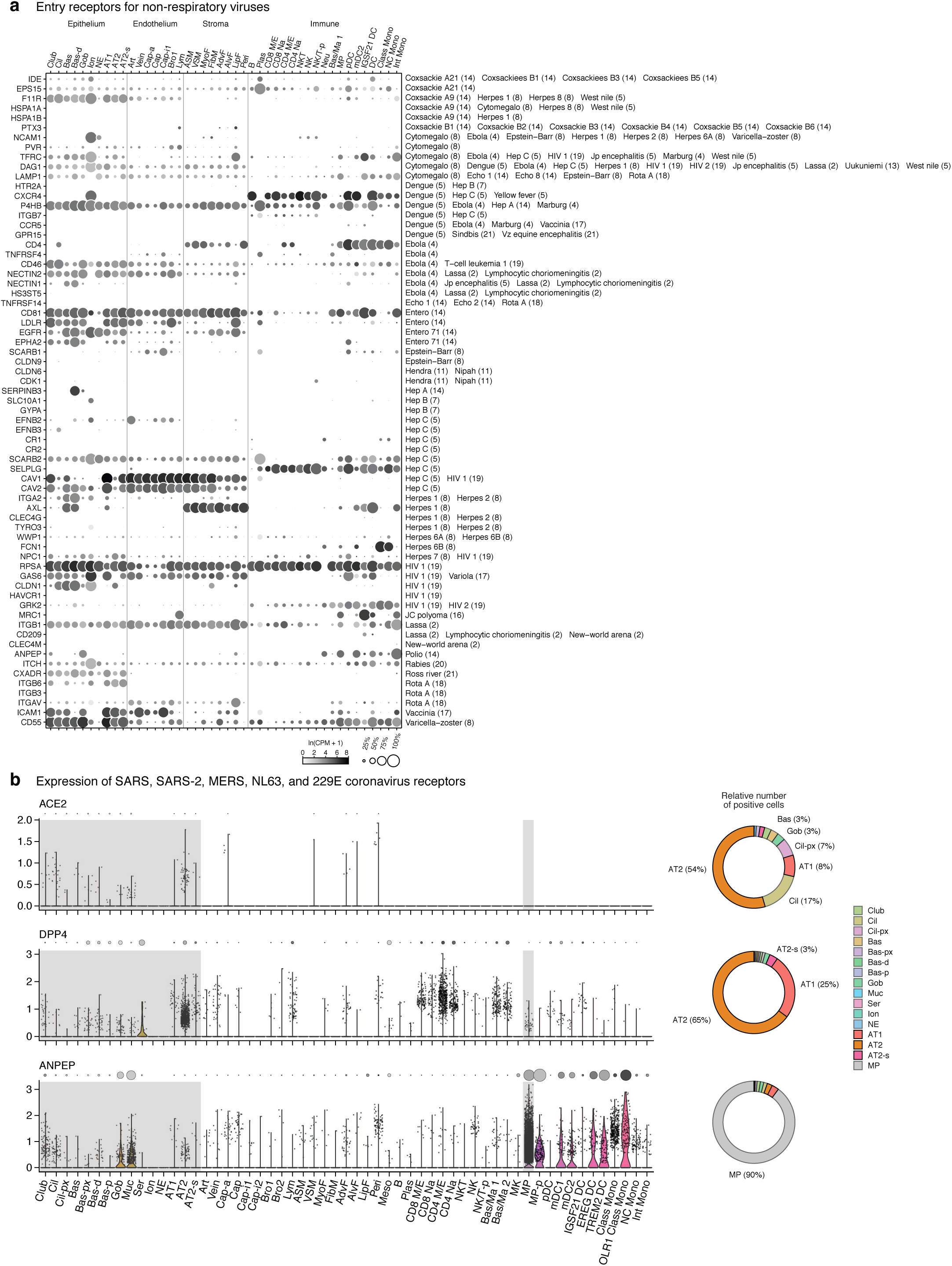
Lung cell expression patterns of viral entry receptors. **a**, Dot plot showing gene expression of entry receptors for non-respiratory viruses (compare with Fig. 6b showing expression values of receptors for respiratory viruses). **b,** Violin plots showing low but detectable gene expression of coronavirus receptors *ACE2*, *DPP4*, and *ANPEP* (left). Grey shading, cell types inhaled viruses can directly access. Right, Donut plots showing the relative number of expressing cells of cell types viruses can directly access (shaded grey in panel a), normalized by their abundance values from Table S1 (and refined by the relative abundance values in Figures 2 and S4). Note prevalence of AT2 and AT1 alveolar cells for *ACE2*, the receptor for SARS-CoV and SARS-CoV-2, and for *DPP4,* the receptor for MERS-CoV, in contrast to prevalence of macrophages for *ANPEP*, the receptor for the common cold causing coronavirus 229E.

**Extended Data Figure S9.**
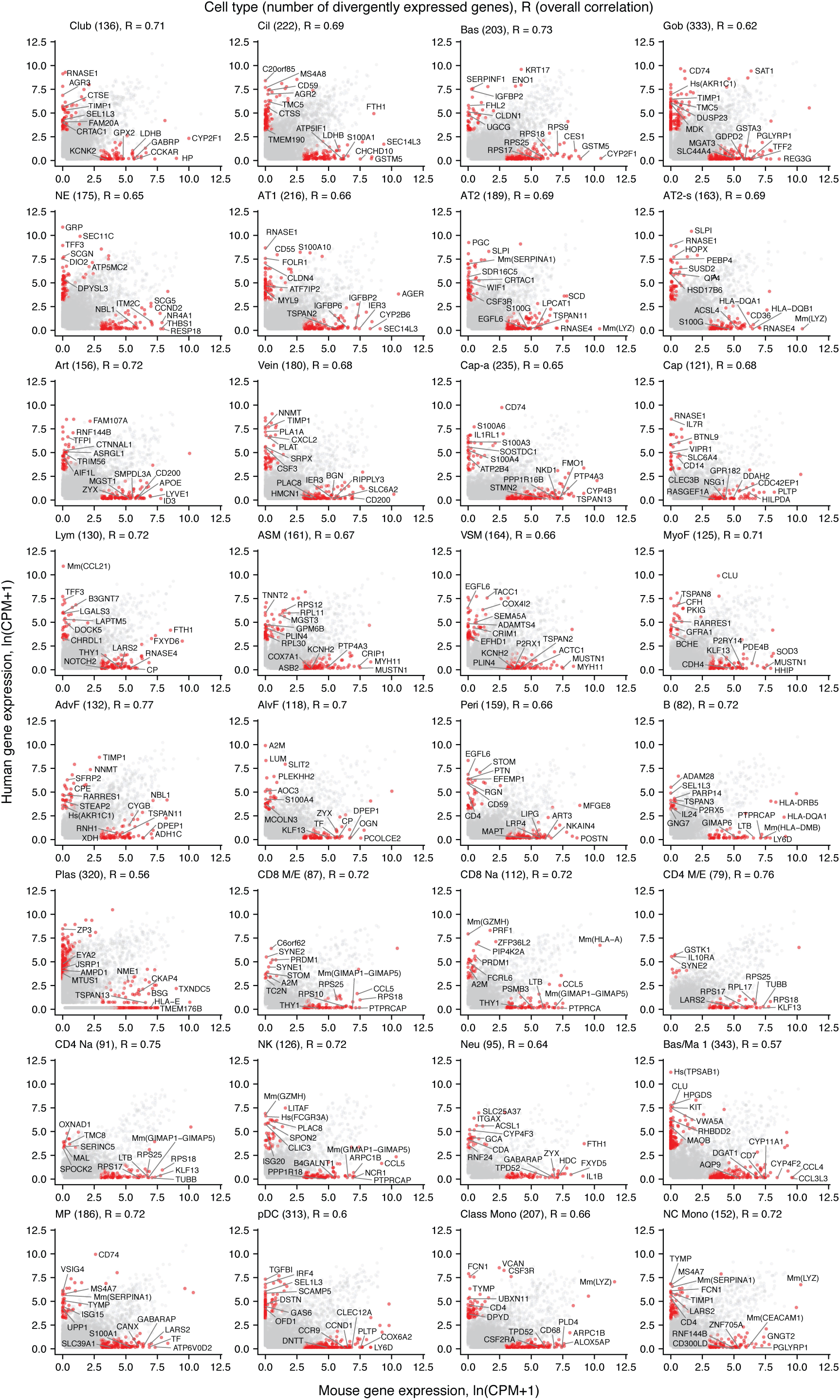
Comparison of mouse and human gene expression profiles in the homologous lung cell types. Scatter plots showing the median expression levels (ln(CPM+1)) in the indicated cell types of each expressed human gene and mouse ortholog in the mouse and human SS2 datasets. Note there are tens to hundreds of genes that show a 20-fold or greater expression difference (and p-value < 0.05) between species (red dots, with gene names indicated for some and total number given above). Bas/Ma 1 cells have the most differentially-expressed genes (343), and CD4^+^ M/E T cells have the least (79). Correlation scores (R values) between the average mouse and human gene expression profiles for each cell type are indicated. “Mm()” and “Hs()”, genes where duplications between mouse and human were collapsed to HomologyID.

**Extended Data Figure S10.**
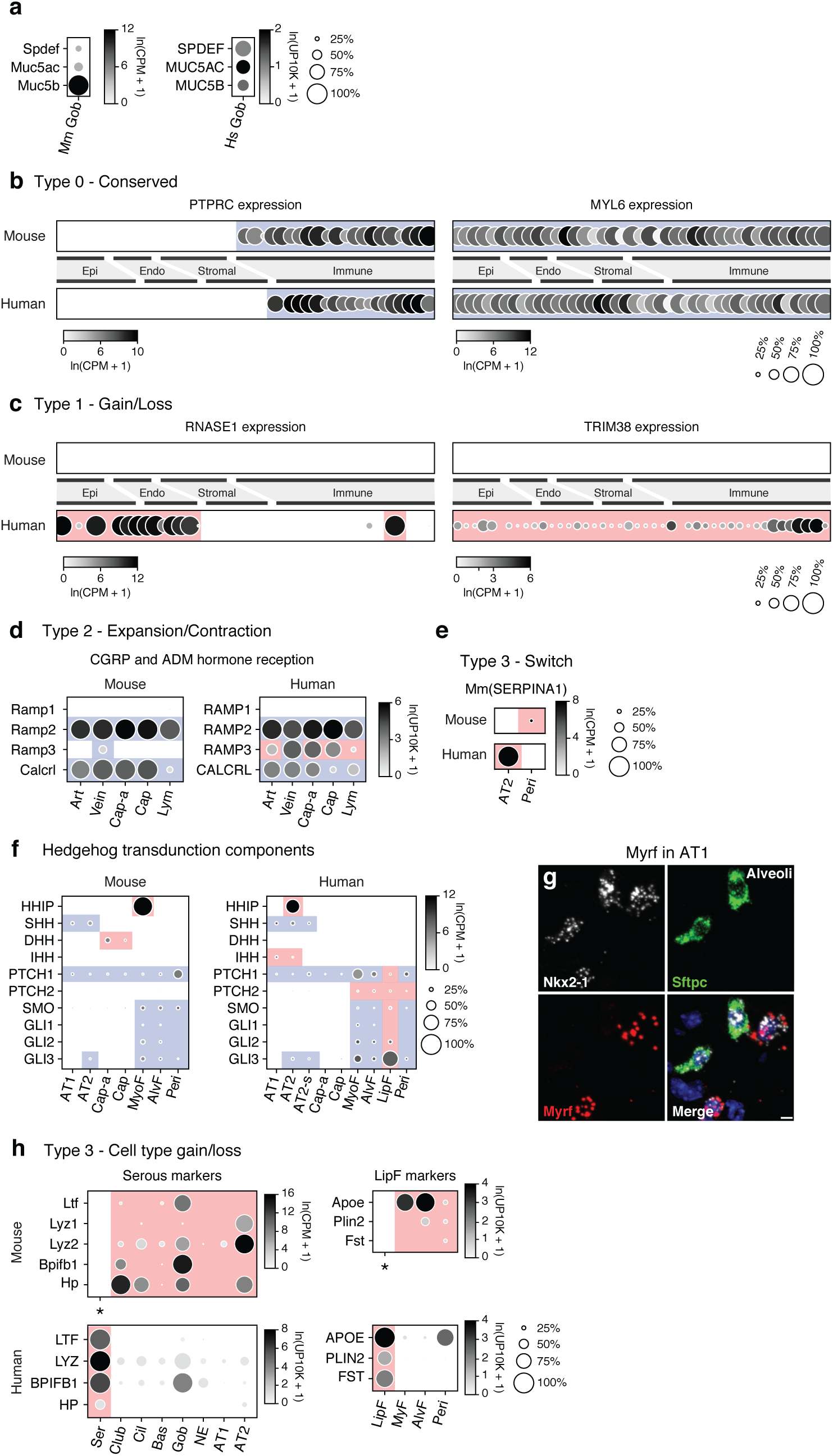
Comparison of gene expression in homologous cell types in human and mouse by scRNAseq. **a**, Dot plot of expression of canonical goblet cell markers *MUC5B* and *MUC5AC* and transcription factor *SPDEF* in mouse and human goblet cells. **b-f**, Additional examples of the four scenarios (Type 0, 1, 2, 3) for evolution of cellular expression pattern as in Fig. 7d. **b,** Dot plots of expression of the indicated genes in mouse and human lung cell types showing conserved (Type 0) expression pattern **c,** Dot plots showing gain of expression (Type 1 change) in multiple human cell types of *RNASE1* (left panel) and all human cell types of *TRIM38* (right panel). **d,** Dot plots of **e**xpression of CGRP and ADM hormone receptor genes showing expansion of expression (Type 2 change) in human endothelial cells. **e,** Dot plots of expression of emphysema-associated gene *SERPINA1* showing switched expression (Type 3 change) from mouse pericytes (top) to human AT2 cells (bottom). **f,** Dot plots comparing expression and conservation of HHIP with those of other Hedgehog pathway genes including ligands (SHH, DHH, IHH), receptors (SMO, PTCH1, PTCH2), and transducers (GLI1, GLI2, GLI3). **g**, Alveolar region of adult mouse lung probed by smFISH for general alveolar epithelial marker *Nkx2-1*, AT2 marker *Sftpc*, and transcription factor *Myrf*. Note *Myrf* is selectively expressed in mouse AT1 cells (*Nkx2-1*+ *Sftpc*-cells), as it is in humans (Fig. 5c). Scale bar, 5 µm. **h,** Dot plots of expression of the serous cell markers *LTF*, *LYZ*, *BPIFBP1*, and *HP* showing switched expression (Type 3 change) from mouse airway epithelial cells to human serous cells, which mice lack. Dot plots of expression of lipid handling genes *APOE*, *PLIN2*, and *FST* show switched expression (Type 3 change) from mouse alveolar stromal cells to human lipofibroblasts, which mice lack. “Mm()” or “Hs()”, genes where duplications between mouse and human were collapsed to HomologyIDs. Red shading, diverged expression; blue shading, conserved expression.

**Extended Data Figure S11.**
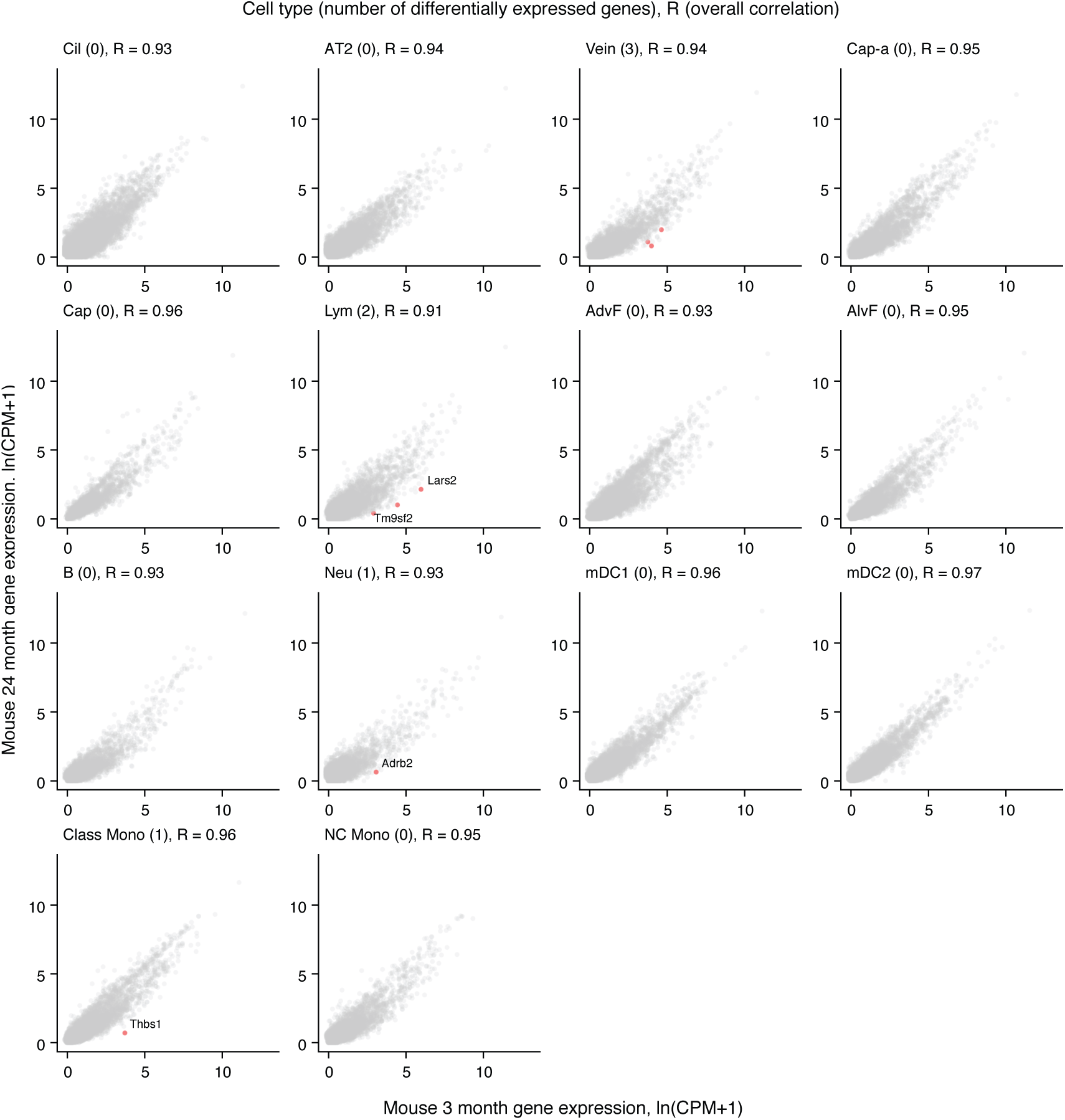
Few genes show >20-fold changes in expression in mouse lung cell types during aging. Scatter plots comparing median expression levels (ln(CPM+1)) in indicated lung cell types of each expressed gene in 3 month (x-axis) and 24 month (y-axis) mouse SS2 datasets from Tabula Muris Senis^56^. Correlation scores between average gene expression profile for each cell type at each age are indicated (R values), along with number of genes (red dots) showing 20-fold or greater expression difference (and p-value < 0.05) between ages. Names of some genes are given next to the corresponding red dot.

## Supplemental Tables

**Table S1.** Canonical cell types (45) in the human lung and their abundances, markers, and available expression data.

| Cell type | Relative abundance (%) | Number (millions)<sup>a</sup> | Canonical markers<sup>b</sup> | Extant expression profiles<br>Single cell | Primary<sup>c</sup> | Expression accession codes | Abundance reference (method)<sup>d</sup> |
| --- | --- | --- | --- | --- | --- | --- | --- |
| **Epithelium** |  |  |  |  |  |  |  |
| Club Cell | 0.5 | 1,500 | CYP2F2, SCGB3A2, CCKAR | Yes | Yes | MTAB-6149, E-MTAB-6653 | Boers et al. 1999 (e) |
| Ciliated Cell | 2 | 6,000 | FOXJ1, TUBB1, TP73, CCDC78 | Yes | Yes | GSE122960 | Raman et al. 2009 (e) |
| Basal Cell | 0.5 | 1,500 | KRT5, KRT14, TP63, DAPL1 | Yes | Yes | MTAB-6149, E-MTAB-6653 | Boers et al. 1998 (e) |
| Goblet Cell | 0.2 | 500 | MUC5B, MUC5AC, SPDEF | Yes | Yes | EGAS00001001755 | Boers et al. 1999 (e) |
| Mucous Cell | 0.03 | 80 | MUC5B |  |  |  | Widdicombe and Wine 2015 (e) |
| Serous Cell | 0.03 | 80 | PRR4, LPO, LTF | Yes | Yes | EGAS00001001755 | Basbaum et al. 1990 (e) |
| Ionocyte | 0.03 | 100 | CFTR, FOXI1, ASCL3 | Yes | Yes | EGAS00001001755 | Montoro et al. 2018 (e) |
| Neuroendocrine Cell | 0.01 | 40 | CALCA, CHGA, ASCL1 | Yes | Yes | EGAS00001001755 | Boers et al. 1996 (e) |
| Tuft Cell | 0.1 | 200 | DCLK1, ASCL2 | Yes |  | GSE102580 | Chang et al. 1986; Montoro et al. 2018 (e) |
| Alveolar Epithelial Type 1 Cell | 13 | 40,000 | AGER, PDPN, CLIC5 | Yes | Yes | MTAB-6149, E-MTAB-6653 | Crapo et al. 1982 (f) |
| Alveolar Epithelial Type 2 Cell | 7 | 20,000 | SFTPB, SFTPC, SFTPD, MUC1, ETV5 | Yes | Yes | GSE122960 | Crapo et al. 1982; Fehrenbach et al. 1994 (f) |
| Total | 23 | 70,000 |  |  |  |  |  |
| **Endothelium** |  |  |  |  |  |  |  |
| Artery Cell | 1 | 3,000 | GJA5, BMX | (bulk) | (cultured) | phs000998.v1.p1 | Townsend et al. 2012; The Lung, Chapter 74 (g) |
| Vein Cell | 1 | 3,000 | ACKR1 |  |  |  | Townsend et al. 2012; The Lung, Chapter 74 (g) |
| Capillary Cell | 23 | 70,000 | CA4 |  |  |  | Crapo et al. 1982 (f) |
| Bronchial Vessel | 0.7 | 2,000 |  |  |  |  | Deffebach et al. 1987 (g) |
| Lymphatic Cell | 0.7 | 2,000 | PROX1, PDPN | Yes | Yes | MTAB-6149, E-MTAB-6653 | Kambouchner et al. 2009; Sozio et al. 2012 (g) |
| Total | 27 | 80,000 |  | (unannotated) |  | MTAB-6149, E-MTAB-6653, EGAS00001001755 |  |
| **Stroma** |  |  |  |  |  |  |  |
| Vascular Smooth Muscle | 2 | 5,000 | CNN1, ACTA2, TAGLN, RGS5 | Yes | Yes | GSE75990 | Townsend et al. 2012; The Lung, Chapter 74 (h) |
| Airway Smooth Muscle | 1 | 4,000 | CNN1, ACTA2, TAGLN, DES, LGR6 |  |  | GSE75990 | Elliot et al. 1999; The Lung, Chapter 74 (h) |
| Fibroblast | 7 | 20,000 | COL1A1, PDGFRA | Yes | Yes | EGAS00001001755 | Crapo et al. 1982 (f,i) |
| Myofibroblast | 7 | 20,000 | COL1A1, PDGFRA, ELN, ACTA2 | Yes | Yes | EGAS00001001755 | Crapo et al. 1982 (f,i) |
| Lipofibroblast | 7 | 20,000 | COL1A1, PDGFRA, PLIN2, APOE |  |  |  | Crapo et al. 1982 (f,i) |
| Pericyte | 7 | 20,000 | CSPG4, TRPC6, PDGFRB | (bulk) | (cultured) | GSE75990 | Crapo et al. 1982 (f,i) |
| Mesothelial Cell | 0.3 | 1,000 | MSLN, UPK3B, WT1 | (bulk) | (cultured) | GSE63966 | Michailova et al. 1997 (j) |
| Total | 30 | 90,000 |  | (unannotated) |  | MTAB-6149, E-MTAB-6653, EGAS00001001755 |  |
| **PNS** |  |  |  |  |  |  |  |
| Intrinsic Neuron | 0.0003 | 1 | SNAP25 |  |  |  | Fox et al. 1980; Sparrow et al. 1999 (j) |
| Glial Cell | 0.0002 | 0.5 |  |  |  |  | Sparrow et al. 1999 (j) |
| Total | 0.0005 | 1.5 |  |  |  |  |  |
| **Immune** |  |  |  |  |  |  |  |
| B Cell | 0.5 | 1,500 | CD79A, CD24, MS4A1, CD19 | Yes | Yes | E-MTAB-6701, E-MTAB-6678 | Finkelstein et al. 1995; Banat et al. 2015 (k) |
| Plasma Cell | 0.7 | 2,000 | CD79A, CD27, SLAMF7 | Yes | Yes | E-MTAB-6701, E-MTAB-6678 | Banat et al. 2015 (k) |
| CD8+ Mem/Eff T Cell | 1 | 3,000 | CD3E, CD8A, GZMK, DUSP2 | Yes | Yes | E-MTAB-6701, E-MTAB-6678 | Finkelstein et al. 1995; Banat et al. 2015 (k,i) |
| CD8+ Naive T Cell | 1 | 3,000 | CD3E, CD8, GZMH, GZMB | Yes | Yes | MTAB-6149, E-MTAB-6653 | Finkelstein et al. 1995; Banat et al. 2015 (k,i) |
| CD4+ Mem/Eff Cell | 0.7 | 2,000 | CD3E, CD8, COTL1, LDHB | Yes | Yes | E-MTAB-6701, E-MTAB-6678 | Finkelstein et al. 1995; Banat et al. 2015 (k,i) |
| CD4+ Naive T Cell | 0.7 | 2,000 | CD3E, CD4, CCR7, LEF1 | Yes | Yes | MTAB-6149, E-MTAB-6653 | Finkelstein et al. 1995; Banat et al. 2015 (k,i) |
| Natural Killer Cell | 1 | 3,000 | KLRD1, NKG7, TYROBP | Yes | Yes | E-MTAB-6701, E-MTAB-6678 | Marquardt et al. 2017 (l) |
| Natural Killer T Cell | 0.7 | 2,000 | CD3E, CD8A, FCER1G, TYROBP | Yes | Yes | MTAB-6149, E-MTAB-6653 | Marquardt et al. 2017 (k,i) |
| Neutrophil | 0.8 | 2,500 | S100A8, S100A9, IFITM2, FCGR3B | Yes | Yes | EGAS00001001755 | Finkelstein et al. 1995; Banat et al. 2015 (k) |
| Basophil | 0.3 | 1,000 | MS4A2, CPA3, TPSAB1 | Yes | Yes | MTAB-6149, E-MTAB-6653 | Finkelstein et al. 1995 (k,i) |
| Mast Cell | 1 | 3,000 | MS4A2, CPA3, TPSAB1 | Yes | Yes | MTAB-6149, E-MTAB-6653 | Finkelstein et al. 1995; Banat et al. 2015 (k) |
| Eosinophil | 0.3 | 1,000 | SIGLEC8 | (bulk) | (cultured) |  | Finkelstein et al. 1995 (k,i) |
| Megakaryocyte | 0.3<sup>i</sup> | 1,000 | NRGN, PPBP, PF4, OST4 | (bulk) | Yes |  | Dejima et al. 2018; Skoczynski et al. 2019 (m) |
| Macrophage | 7 | 20,000 | MARCO, MSR1, MRC1 | Yes | Yes | MTAB-6149, E-MTAB-6653 | Crapo et al. 1982; Fehrenbach et al. 1994 (f) |
| Plasmacytoid Dendritic Cell | 0.3 | 800 | LILRB4, IRF8, LILRA4 | Yes | Yes | GSE94820 | Banat et al. 2015 (k,i) |
| Myeloid Dendritic Cell 1 | 0.3 | 1,000 | MHCII, CLEC9A, LAMP3 | Yes | Yes | GSE94820 | Banat et al. 2015 (k) |
| Myeloid Dendritic Cell 2 | 0.1 | 200 | MHCII, CD1C, PLD4 | Yes | Yes | GSE94820 | Banat et al. 2015 (k) |
| Classical Monocyte | 2 | 4,000 | CD14, S100A8 | Yes | Yes | E-MTAB-6701, E-MTAB-6678 | Hance et al. 1985; Hoogsteden et al. 1989 (k,i) |
| Intermediate Monocyte | 2 | 4,000 | CD14, S100A8, CD16 | (bulk) | Yes | GSE80095 | Hance et al. 1985; Hoogsteden et al. 1989 (k,i) |
| Nonclassical Monocyte | 1 | 3,000 | CD16 | Yes | Yes | GSE94820 | Hance et al. 1985; Hoogsteden et al. 1989 (k) |
| Total | 20 | 60,000 |  |  |  |  |  |
| Total (all compartments) | 100 | 300,000 |  |  |  |  |  |

**a**, numbers of each type were calculated with their abundances and the total number of lung cells (estimated by comparing volume of lungs to the whole body). **b**, Canonical markers were obtained from referenced expression data or commonly used markers in the literature. **c**, Expression profiles captured immediately following tissue dissociation are considered primary. **d**, Alveoli were assumed to occupy ~90% of the total lung volume for all estimations. **e**, Inferred from mean relative abundance in proximal, medial and distal airway epithelium. **f**, Calculated by stereology **g**, Resin casts showed similar surface area of arteries and veins. **h**, Vascular smooth muscle is estimated to be slightly more abundant than airway smooth muscle. **i**, abundance of a more general cell type was split evenly. **j**, inferred from impression of light or electron microscopy. **k**, inferred from histological abundance in non-perfused healthy tissue. **l**, inferred from abundance among immune cells with FACS. **m**, Calculated using microfluidic capture

**Table S2.**
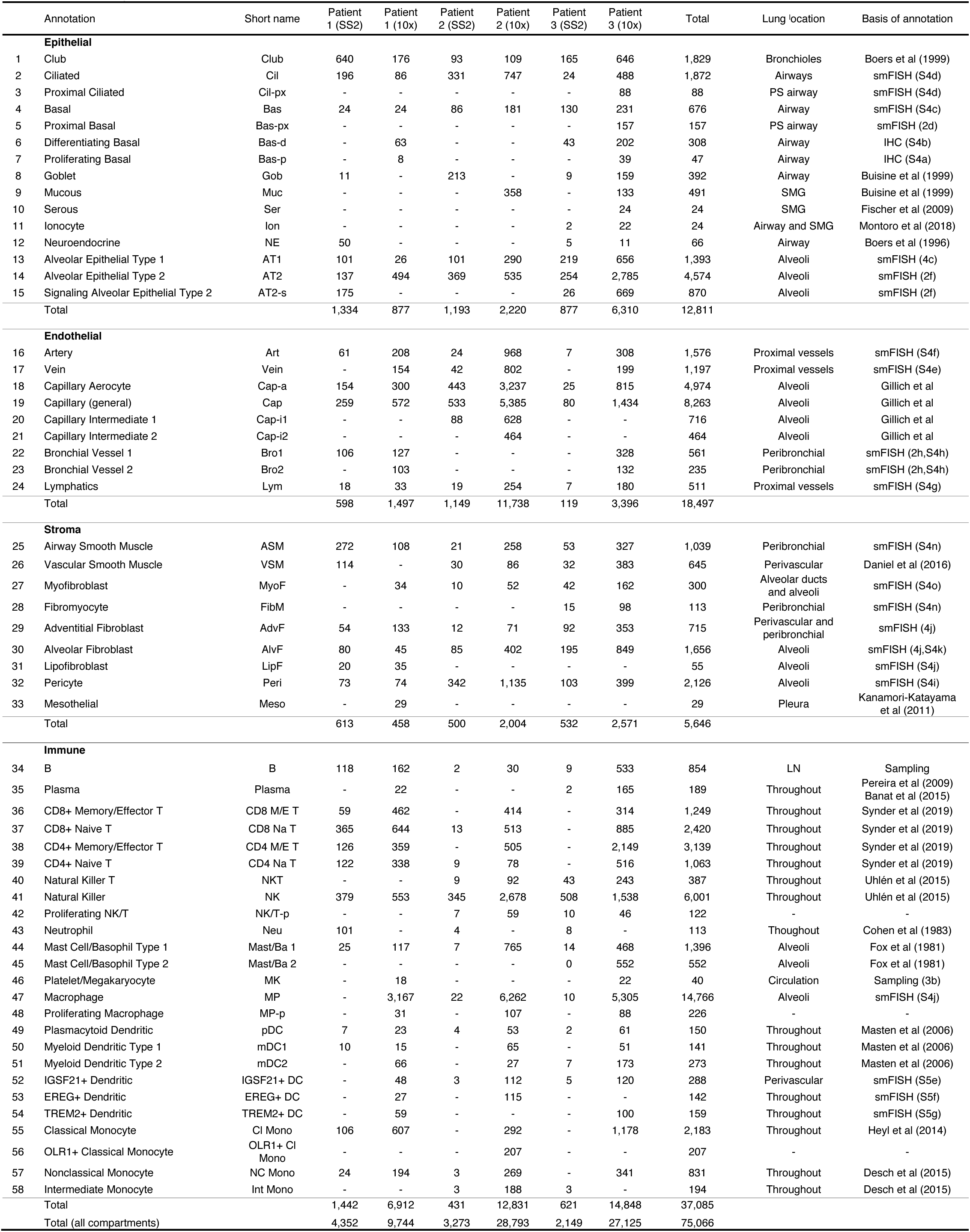
Human lung cell cluster identities and their abundances in each dataset.

|  | Annotation | Short name | Patient 1 (SS2) | Patient 1 (10x) | Patient 2 (SS2) | Patient 2 (10x) | Patient 3 (SS2) | Patient 3 (10x) | Total | Lung location | Basis of annotation |
| --- | --- | --- | --- | --- | --- | --- | --- | --- | --- | --- | --- |
| Epithelial |  |  |  |  |  |  |  |  |  |  |  |
| 1 | Club | Club | 640 | 176 | 93 | 109 | 165 | 646 | 1,829 | Bronchioles | Boers et al (1999) |
| 2 | Ciliated | Cil | 196 | 86 | 331 | 747 | 24 | 488 | 1,872 | Airways | smFISH (S4d) |
| 3 | Proximal Ciliated | Cil-px | - | - | - | - | - | 88 | 88 | PS airway | smFISH (S4d) |
| 4 | Basal | Bas | 24 | 24 | 86 | 181 | 130 | 231 | 676 | Airway | smFISH (S4c) |
| 5 | Proximal Basal | Bas-px | - | - | - | - | - | 157 | 157 | PS airway | smFISH (2d) |
| 6 | Differentiating Basal | Bas-d | - | 63 | - | - | 43 | 202 | 308 | Airway | IHC (S4b) |
| 7 | Proliferating Basal | Bas-p | - | 8 | - | - | - | 39 | 47 | Airway | IHC (S4a) |
| 8 | Goblet | Gob | 11 | - | 213 | - | 9 | 159 | 392 | Airway | Buisine et al (1999) |
| 9 | Mucous | Muc | - | - | - | 358 | - | 133 | 491 | SMG | Buisine et al (1999) |
| 10 | Serous | Ser | - | - | - | - | - | 24 | 24 | SMG | Fischer et al (2009) |
| 11 | Ionocyte | Ion | - | - | - | - | 2 | 22 | 24 | Airway and SMG | Montoro et al (2018) |
| 12 | Neuroendocrine | NE | 50 | - | - | - | 5 | 11 | 66 | Airway | Boers et al (1996) |
| 13 | Alveolar Epithelial Type 1 | AT1 | 101 | 26 | 101 | 290 | 219 | 656 | 1,393 | Alveoli | smFISH (4c) |
| 14 | Alveolar Epithelial Type 2 | AT2 | 137 | 494 | 369 | 535 | 254 | 2,785 | 4,574 | Alveoli | smFISH (2f) |
| 15 | Signaling Alveolar Epithelial Type 2 | AT2-s | 175 | - | - | - | 26 | 669 | 870 | Alveoli | smFISH (2f) |
|  | Total |  | 1,334 | 877 | 1,193 | 2,220 | 877 | 6,310 | 12,811 |  |  |
| Endothelial |  |  |  |  |  |  |  |  |  |  |  |
| 16 | Artery | Art | 61 | 208 | 24 | 968 | 7 | 308 | 1,576 | Proximal vessels | smFISH (S4f) |
| 17 | Vein | Vein | - | 154 | 42 | 802 | - | 199 | 1,197 | Proximal vessels | smFISH (S4e) |
| 18 | Capillary Aerocyte | Cap-a | 154 | 300 | 443 | 3,237 | 25 | 815 | 4,974 | Alveoli | Gillich et al |
| 19 | Capillary (general) | Cap | 259 | 572 | 533 | 5,385 | 80 | 1,434 | 8,263 | Alveoli | Gillich et al |
| 20 | Capillary Intermediate 1 | Cap-i1 | - | - | 88 | 628 | - | - | 716 | Alveoli | Gillich et al |
| 21 | Capillary Intermediate 2 | Cap-i2 | - | - | - | 464 | - | - | 464 | Alveoli | Gillich et al |
| 22 | Bronchial Vessel 1 | Bro1 | 106 | 127 | - | - | - | 328 | 561 | Peribronchial | smFISH (2h,S4h) |
| 23 | Bronchial Vessel 2 | Bro2 | - | 103 | - | - | - | 132 | 235 | Peribronchial | smFISH (2h,S4h) |
| 24 | Lymphatics | Lym | 18 | 33 | 19 | 254 | 7 | 180 | 511 | Proximal vessels | smFISH (S4g) |
|  | Total |  | 598 | 1,497 | 1,149 | 11,738 | 119 | 3,396 | 18,497 |  |  |
| Stroma |  |  |  |  |  |  |  |  |  |  |  |
| 25 | Airway Smooth Muscle | ASM | 272 | 108 | 21 | 258 | 53 | 327 | 1,039 | Peribronchial | smFISH (S4n) |
| 26 | Vascular Smooth Muscle | VSM | 114 | - | 30 | 86 | 32 | 383 | 645 | Perivascular | Daniel et al (2016) |
| 27 | Myofibroblast | MyoF | - | 34 | 10 | 52 | 42 | 162 | 300 | Alveolar ducts and alveoli | smFISH (S4o) |
| 28 | Fibromyocyte | FibM | - | - | - | - | 15 | 98 | 113 | Peribronchial | smFISH (S4n) |
| 29 | Adventitial Fibroblast | AdvF | 54 | 133 | 12 | 71 | 92 | 353 | 715 | Perivascular and peribronchial | smFISH (4j) |
| 30 | Alveolar Fibroblast | AlvF | 80 | 45 | 85 | 402 | 195 | 849 | 1,656 | Alveoli | smFISH (4j,S4k) |
| 31 | Lipofibroblast | LipF | 20 | 35 | - | - | - | - | 55 | Alveoli | smFISH (S4j) |
| 32 | Pericyte | Peri | 73 | 74 | 342 | 1,135 | 103 | 399 | 2,126 | Alveoli | smFISH (S4i) |
| 33 | Mesothelial | Meso | - | 29 | - | - | - | - | 29 | Pleura | Kanamori-Katayama et al (2011) |
|  | Total |  | 613 | 458 | 500 | 2,004 | 532 | 2,571 | 5,646 |  |  |
| Immune |  |  |  |  |  |  |  |  |  |  |  |
| 34 | B | B | 118 | 162 | 2 | 30 | 9 | 533 | 854 | LN | Sampling |
| 35 | Plasma | Plasma | - | 22 | - | - | 2 | 165 | 189 | Throughout | Pereira et al (2009)<br>Banat et al (2015) |
| 36 | CD8+ Memory/Effector T | CD8 M/E T | 59 | 462 | - | 414 | - | 314 | 1,249 | Throughout | Synder et al (2019) |
| 37 | CD8+ Naive T | CD8 Na T | 365 | 644 | 13 | 513 | - | 885 | 2,420 | Throughout | Synder et al (2019) |
| 38 | CD4+ Memory/Effector T | CD4 M/E T | 126 | 359 | - | 505 | - | 2,149 | 3,139 | Throughout | Synder et al (2019) |
| 39 | CD4+ Naive T | CD4 Na T | 122 | 338 | 9 | 78 | - | 516 | 1,063 | Throughout | Synder et al (2019) |
| 40 | Natural Killer T | NKT | - | - | 9 | 92 | 43 | 243 | 387 | Throughout | Uhlén et al (2015) |
| 41 | Natural Killer | NK | 379 | 553 | 345 | 2,678 | 508 | 1,538 | 6,001 | Throughout | Uhlén et al (2015) |
| 42 | Proliferating NK/T | NK/T-p | - | - | 7 | 59 | 10 | 46 | 122 | - | - |
| 43 | Neutrophil | Neu | 101 | - | 4 | - | 8 | - | 113 | Thougthout | Cohen et al (1983) |
| 44 | Mast Cell/Basophil Type 1 | Mast/Ba 1 | 25 | 117 | 7 | 765 | 14 | 468 | 1,396 | Alveoli | Fox et al (1981) |
| 45 | Mast Cell/Basophil Type 2 | Mast/Ba 2 | - | - | - | - | 0 | 552 | 552 | Alveoli | Fox et al (1981) |
| 46 | Platelet/Megakaryocyte | MK | - | 18 | - | - | - | 22 | 40 | Circulation | Sampling (3b) |
| 47 | Macrophage | MP | - | 3,167 | 22 | 6,262 | 10 | 5,305 | 14,766 | Alveoli | smFISH (S4j) |
| 48 | Proliferating Macrophage | MP-p | - | 31 | - | 107 | - | 88 | 226 | - | - |
| 49 | Plasmacytoid Dendritic | pDC | 7 | 23 | 4 | 53 | 2 | 61 | 150 | Throughout | Masten et al (2006) |
| 50 | Myeloid Dendritic Type 1 | mDC1 | 10 | 15 | - | 65 | - | 51 | 141 | Throughout | Masten et al (2006) |
| 51 | Myeloid Dendritic Type 2 | mDC2 | - | 66 | - | 27 | 7 | 173 | 273 | Throughout | Masten et al (2006) |
| 52 | IGSF21+ Dendritic | IGSF21+ DC | - | 48 | 3 | 112 | 5 | 120 | 288 | Perivascular | smFISH (S5e) |
| 53 | EREG+ Dendritic | EREG+ DC | - | 27 | - | 115 | - | - | 142 | Throughout | smFISH (S5f) |
| 54 | TREM2+ Dendritic | TREM2+ DC | - | 59 | - | - | - | 100 | 159 | Throughout | smFISH (S5g) |
| 55 | Classical Monocyte | Cl Mono | 106 | 607 | - | 292 | - | 1,178 | 2,183 | Throughout | Heyl et al (2014) |
| 56 | OLR1+ Classical Monocyte | OLR1+ Cl Mono | - | - | - | 207 | - | - | 207 | - | - |
| 57 | Nonclassical Monocyte | NC Mono | 24 | 194 | 3 | 269 | - | 341 | 831 | Throughout | Desch et al (2015) |
| 58 | Intermediate Monocyte | Int Mono | - | - | 3 | 188 | 3 | - | 194 | Throughout | Desch et al (2015) |
|  | Total |  | 1,442 | 6,912 | 431 | 12,831 | 621 | 14,848 | 37,085 |  |  |
|  | Total (all compartments) |  | 4,352 | 9,744 | 3,273 | 28,793 | 2,149 | 27,125 | 75,066 |  |  |

**Table S3.** Surface markers used for canonical immune cell types in bulk mRNA sequencing.

| Immune cell types and populations | Surface markers used for sorting |
| --- | --- |
| Neutrophil | SSC<sup>hi</sup>CD16+CD123-CCR3-b7 integrin- |
| Eosinophil | SSC<sup>hi</sup>CD16-CD123-CCR3+b7 integrin+ |
| Basophil | SSC<sup>lo</sup>CD16-CD123<sup>hi</sup>CCR3+b7 integrin+ |
| Classical monocyte | FSC<sup>hi</sup>CD4<sup>lo</sup>CD14<sup>hi</sup>CD16<sup>lo</sup> |
| Non-classical monocyte | FSC<sup>hi</sup>CD4<sup>lo</sup>CD14<sup>-/lo</sup>CD16<sup>hi</sup> |
| Myeloid dendritic cell | CD11c<sup>hi</sup>CD123<sup>lo</sup>CD1c+HLA-DR<sup>hi</sup>CD16- |
| CD16+ dendritic cell | CD11c<sup>hi</sup>CD123<sup>lo</sup>CD1c-HLA-DR+CD16+ |
| Plasmacytoid dendritic cell | CD11c-CD123<sup>hi</sup>CCR3-HLA-DR+b7 integrin- |
| Naïve B cell | CD19+CD20+CD27-IgM/D+ |
| Unswitched memory B cell | CD19+CD20+CD27+IgM/D+ |
| Switched memory B cell | CD19+CD20+CD27+IgM/D- |
| Immature NK cell | CD16-CD56<sup>hi</sup>CD57- |
| Mature NK cell | CD16+CD56+CD57- |
| More mature NK cell | CD16+CD56+CD57+ |
| Naïve CD4-T cell | CD4+CD45RA+CD45RO-CCR7+CD62L+ |
| Central memory CD4-T cell | CD4+CD45RA-CD45RO+CCR7+CD62L+ |
| Effector memory CD4-T cell | CD4+CD45RA-CD45RO+CCR7-CD62L- |
| Naïve CD8-T cell | CD8+CD45RA+CD45RO-CCR7+CD62L+ |
| Central memory CD8-T cell | CD8+CD45RA-CD45RO+CCR7+CD62L+ |
| Effector memory CD8-T cell | CD8+CD45RA-CD45RO+CCR7-CD62L- |
| Effector memory CD45RA+ CD8-T cell | CD8+CD45RA+CD45RO-CCR7-CD62L- |

**Table S4.** Enriched markers found in each cluster, with transcription factors, receptors/ligands, and disease associated genes annotated.

| Gene | avg_logFC | pct_in_cluster | pct_out_cluster | p_val | p_val_adj | TF | Receptor | Ligand | Enzymes | OMIM | GWAS |
| --- | --- | --- | --- | --- | --- | --- | --- | --- | --- | --- | --- |
| SCGB3A2 | 2.63 | 0.98 | 0.47 | 4.38E-51 | 2.67E-46 |  |  | TRUE |  | TRUE |  |
| MGP | 3.87 | 0.68 | 0.15 | 2.71E-41 | 1.65E-36 |  |  |  |  | TRUE |  |
| PIGR | 1.66 | 0.95 | 0.58 | 4.85E-34 | 2.96E-29 |  |  |  |  |  |  |
| VIM | 1.8 | 0.92 | 0.42 | 5.5E-33 | 3.35E-28 |  |  | TRUE |  |  |  |
| SCGB3A1 | 2.91 | 0.91 | 0.38 | 9.5E-33 | 5.79E-28 |  |  | TRUE |  |  |  |
| TIMP1 | 1.57 | 0.92 | 0.37 | 7.32E-32 | 4.46E-27 |  |  | TRUE |  |  |  |
| KLK11 | 1.41 | 0.87 | 0.28 | 1.75E-30 | 1.07E-25 |  |  |  |  |  |  |
| FAM129A | 1.45 | 0.76 | 0.17 | 7.28E-30 | 4.44E-25 |  |  |  |  |  |  |
| FMO2 | 1.57 | 0.84 | 0.25 | 4.89E-29 | 2.98E-24 |  |  |  | TRUE |  |  |
| SCGB1A1 | 3.23 | 0.5 | 0.08 | 3.27E-28 | 1.99E-23 |  |  | TRUE |  |  |  |
| KLK10 | 1.94 | 0.75 | 0.19 | 4.29E-28 | 2.62E-23 |  |  |  |  |  |  |
| SCNN1B | 1.41 | 0.9 | 0.38 | 2.32E-27 | 1.42E-22 |  | TRUE |  |  |  |  |
| CTSE | 1.51 | 0.87 | 0.36 | 3.87E-27 | 2.36E-22 |  |  |  |  |  |  |
| CXCL17 | 1.05 | 0.98 | 0.55 | 5.09E-26 | 3.1E-21 |  |  |  |  |  |  |
| CX3CL1 | 2.61 | 0.49 | 0.05 | 2.19E-25 | 1.33E-20 |  |  | TRUE |  |  |  |
| STEAP4 | 1.44 | 0.89 | 0.45 | 2.07E-24 | 1.26E-19 |  |  |  |  |  |  |
| CRACR2B | 1.47 | 0.8 | 0.29 | 5.04E-24 | 3.07E-19 |  |  |  |  |  |  |
| CP | 1.46 | 0.8 | 0.27 | 5.86E-24 | 3.57E-19 |  |  | TRUE |  |  |  |
| KLK13 | 2.84 | 0.37 | 0.02 | 2.53E-23 | 1.55E-18 |  |  |  |  |  |  |
| C16orf89 | 0.82 | 0.98 | 0.53 | 7.54E-23 | 4.6E-18 |  |  |  |  |  |  |
| CLU | 1.21 | 0.97 | 0.67 | 1.41E-22 | 8.62E-18 |  |  |  |  |  | TRUE |
| CYP2F1 | 3.35 | 0.38 | 0.02 | 1.43E-22 | 8.73E-18 |  |  |  |  |  |  |
| ERP27 | 1.41 | 0.76 | 0.23 | 2.01E-22 | 1.23E-17 |  |  |  |  |  |  |
| CD82 | 1.63 | 0.76 | 0.26 | 3.59E-22 | 2.19E-17 |  | TRUE |  |  |  |  |
| KDR | 2.36 | 0.36 | 0.02 | 8.78E-22 | 5.35E-17 |  | TRUE |  |  |  |  |
| SFTPB | 0.86 | 1 | 0.68 | 1.46E-21 | 8.91E-17 |  |  |  |  | TRUE |  |
| SOX4 | 1.73 | 0.88 | 0.63 | 3.58E-21 | 2.18E-16 | TRUE |  |  |  |  |  |
| CYP2B7P | 0.85 | 0.98 | 0.7 | 4.01E-21 | 2.44E-16 |  |  |  |  |  |  |
| MET | 1.27 | 0.88 | 0.47 | 9.63E-21 | 5.87E-16 |  | TRUE |  |  |  |  |
| CFH | 1.86 | 0.58 | 0.16 | 2.67E-19 | 1.63E-14 |  |  | TRUE |  |  | TRUE |
| ANPEP | 1.93 | 0.45 | 0.07 | 8.76E-19 | 5.34E-14 |  |  |  |  |  |  |
| MFSD4A | 1.38 | 0.54 | 0.11 | 1.04E-18 | 6.36E-14 |  |  |  |  |  |  |
| CYB5A | 0.93 | 0.97 | 0.9 | 1.38E-18 | 8.44E-14 |  |  |  |  |  |  |
| WFDC2 | 1.02 | 0.97 | 0.87 | 2.42E-18 | 1.48E-13 |  |  |  |  |  |  |
| CYP2A6 | 3.28 | 0.36 | 0.04 | 2.86E-18 | 1.74E-13 |  |  |  |  | TRUE | TRUE |
| NOTCH3 | 1.73 | 0.53 | 0.12 | 4.29E-18 | 2.61E-13 |  | TRUE |  |  |  |  |
| RNASE1 | 0.85 | 0.99 | 0.68 | 1.23E-17 | 7.52E-13 |  |  |  |  |  |  |
| CTSC | 1.49 | 0.77 | 0.39 | 3.04E-17 | 1.85E-12 |  |  |  |  |  |  |
| SLPI | 0.39 | 0.98 | 0.69 | 3.41E-17 | 2.08E-12 |  |  | TRUE |  |  |  |
| TMEM45A | 3.48 | 0.28 | 0.03 | 7.6E-17 | 4.64E-12 |  |  |  |  |  |  |
| TSPAN8 | 2.33 | 0.38 | 0.06 | 2.41E-16 | 1.47E-11 |  |  |  |  |  | TRUE |
| NT5E | 2.13 | 0.44 | 0.08 | 2.7E-16 | 1.65E-11 |  | TRUE |  |  |  |  |
| HNMT | 1.43 | 0.73 | 0.29 | 5.02E-16 | 3.06E-11 |  |  |  | TRUE | TRUE |  |
| PAG1 | 1.4 | 0.45 | 0.08 | 6.54E-16 | 3.99E-11 |  |  |  |  |  |  |
| AARD | 2.41 | 0.51 | 0.15 | 7.64E-16 | 4.66E-11 |  |  |  |  |  |  |
| HSD17B13 | 2.33 | 0.44 | 0.09 | 8.27E-16 | 5.04E-11 |  |  |  |  |  |  |
| CEACAM6 | 0.66 | 0.91 | 0.53 | 1.03E-15 | 6.27E-11 |  |  |  |  |  |  |
| MDK | 1.11 | 0.75 | 0.34 | 1.52E-15 | 9.25E-11 |  |  | TRUE |  |  |  |
| SEL1L3 | 1.17 | 0.83 | 0.42 | 2.57E-15 | 1.56E-10 |  |  |  |  |  |  |
| ASS1 | 1.28 | 0.71 | 0.28 | 3.11E-15 | 1.9E-10 |  |  |  |  |  |  |
| RHOV | 2.02 | 0.27 | 0.02 | 3.17E-15 | 1.94E-10 |  |  |  |  |  |  |
| GGT5 | 2.29 | 0.48 | 0.14 | 5.04E-15 | 3.07E-10 |  |  |  | TRUE |  |  |
| S100A13 | 0.87 | 0.85 | 0.59 | 1.47E-14 | 8.99E-10 |  |  |  |  |  |  |
| RPS6 | 0.69 | 0.99 | 0.95 | 1.92E-14 | 1.17E-09 |  |  |  |  |  |  |
| B3GNT3 | 2.8 | 0.21 | 0.01 | 4.39E-14 | 2.68E-09 |  |  |  |  |  |  |
| ARHGDIB | 1.55 | 0.72 | 0.37 | 5.1E-14 | 3.11E-09 |  |  |  |  |  |  |
| DHRS3 | 0.93 | 0.95 | 0.74 | 9.2E-14 | 5.61E-09 |  |  |  | TRUE |  |  |
| MMP7 | 2.05 | 0.43 | 0.11 | 9.28E-14 | 5.66E-09 |  |  | TRUE |  |  |  |
| ST6GALNAC | 1.2 | 0.63 | 0.23 | 9.75E-14 | 5.95E-09 |  |  |  |  |  |  |
| MRPS25 | 1.05 | 0.9 | 0.56 | 1.21E-13 | 7.36E-09 |  |  |  |  |  |  |
| SLC39A6 | 1.26 | 0.72 | 0.33 | 1.37E-13 | 8.37E-09 |  | TRUE |  |  |  | TRUE |
| PAEP | 2.79 | 0.2 | 0.01 | 2.88E-13 | 1.76E-08 |  |  |  |  |  |  |
| TCIM | 0.85 | 0.74 | 0.33 | 3.02E-13 | 1.84E-08 |  |  |  |  |  |  |
| RPL13AP5 | 0.63 | 0.97 | 0.91 | 5.64E-13 | 3.44E-08 |  |  |  |  |  |  |
| RPL13P12 | 0.65 | 0.95 | 0.81 | 7.35E-13 | 4.48E-08 |  |  |  |  |  |  |
| TENT5C | 1.33 | 0.72 | 0.36 | 8.5E-13 | 5.18E-08 |  |  |  |  |  |  |
| RPLP0 | 0.71 | 0.99 | 0.91 | 8.91E-13 | 5.43E-08 |  |  |  |  |  |  |
| KIAA1324 | 0.73 | 0.85 | 0.53 | 1.01E-12 | 6.13E-08 |  |  |  |  |  | TRUE |
| SCUBE2 | 1.64 | 0.44 | 0.11 | 1.19E-12 | 7.23E-08 |  |  |  |  |  |  |
| RPL12 | 0.7 | 0.99 | 0.92 | 1.38E-12 | 8.44E-08 |  |  |  |  |  |  |
| CLIC6 | 0.83 | 0.79 | 0.4 | 1.66E-12 | 1.01E-07 |  |  |  |  |  |  |
| INPP4B | 2.14 | 0.25 | 0.02 | 1.95E-12 | 1.19E-07 |  |  |  |  |  |  |
| TSPAN11 | 1.83 | 0.4 | 0.09 | 2.29E-12 | 1.4E-07 |  |  |  |  |  | TRUE |

75 enriched genes are shown for first cluster (Club), table continues to include remaining enriched genes (p-val > 0.05) with remaining cell types in separate sheets. Abbreviations: avg\_logFC, the natural log of the average fold change between the cell type and other cell types in its tissue compartment; pct\_in\_cluster, percentage of cells within the cluster that express the gene; pct\_out\_cluster, percentage of cells outside cluster that express the gene; p\_val\_adj, p-value with Bonferroni correction applied; TF, transcription factor; OMIM, Online Mendelian Inheritance in Man; GWAS, genome wide association study.

**Table S5.**
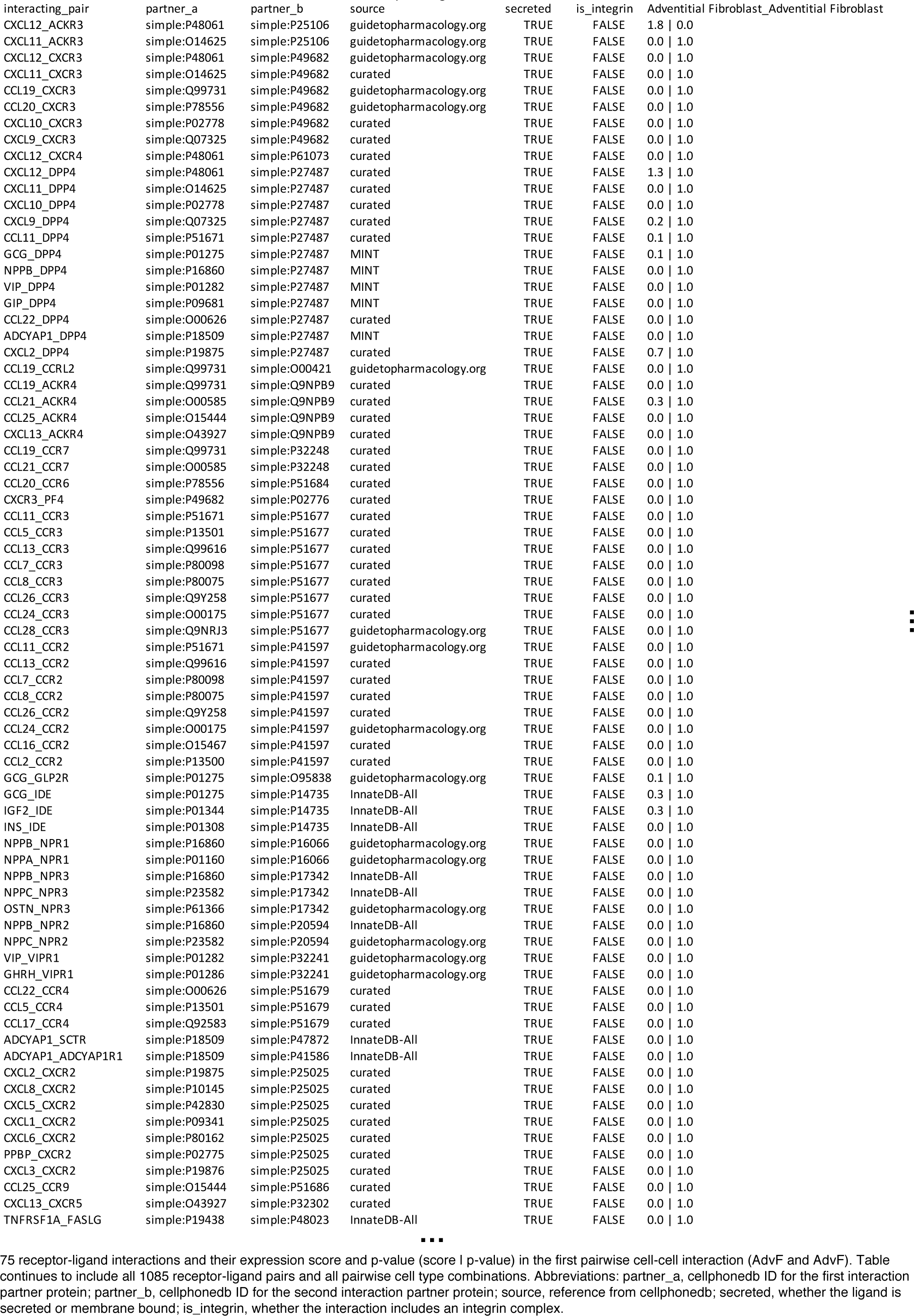
P-value and scores of each CellPhoneDB Receptor-Ligand interaction from each cluster.

**Table S6.**
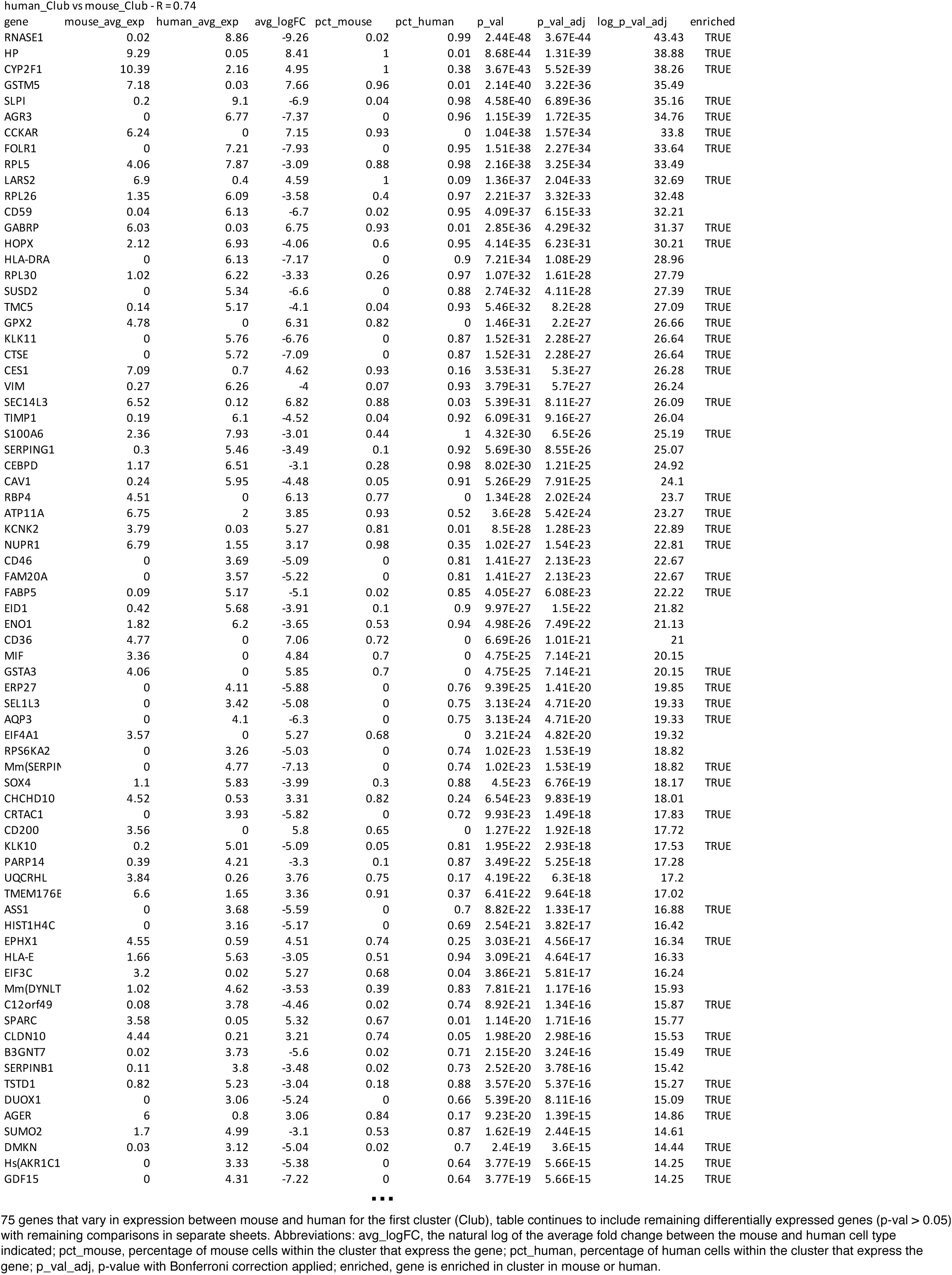
Genes specific to mouse and human in each cluster and lung wide.

**Table S7.**
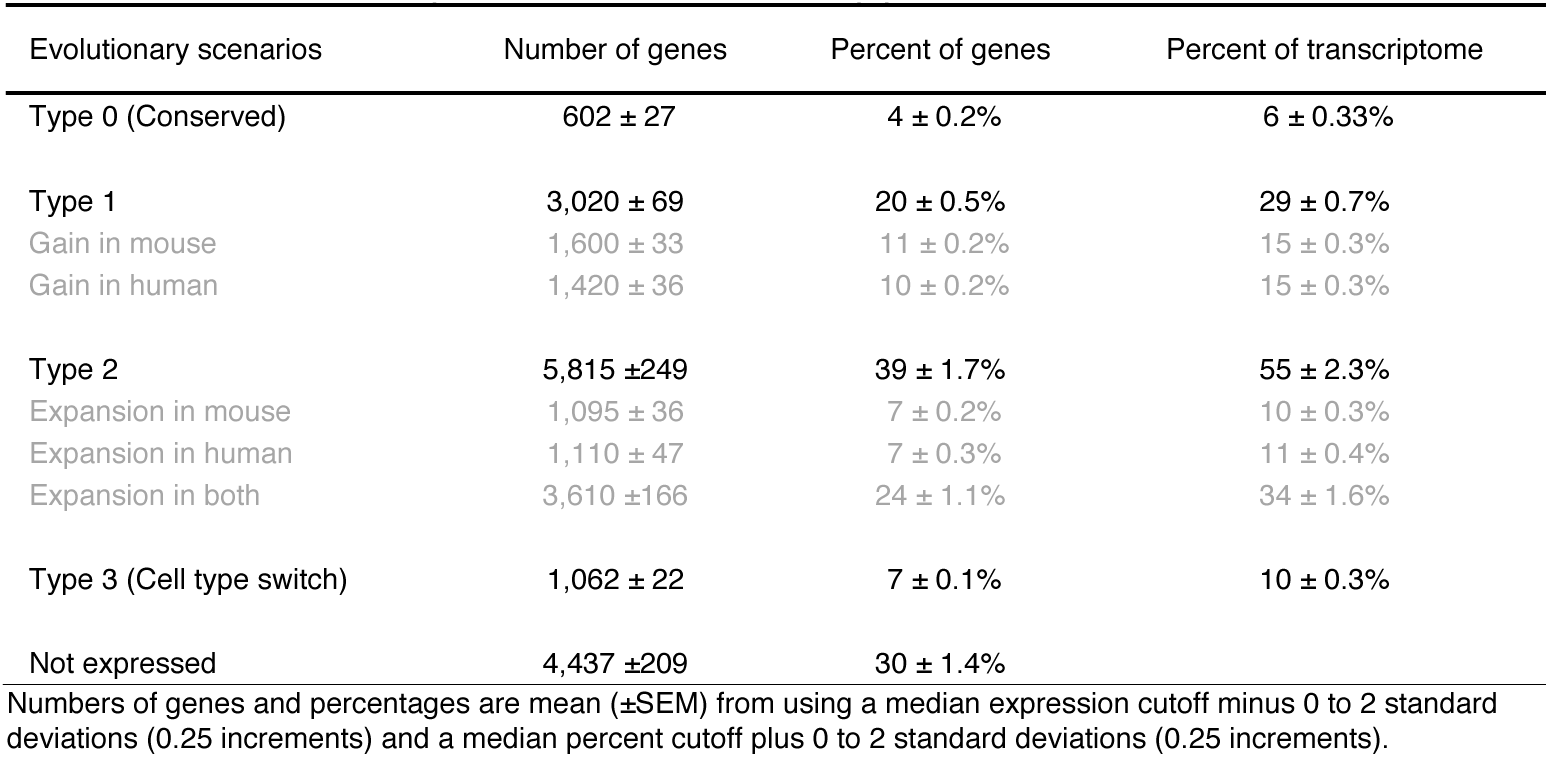
Evolutionary changes in cellular patterns of lung gene expression between mouse and human.

| Evolutionary scenarios | Number of genes | Percent of genes | Percent of transcriptome |
| --- | --- | --- | --- |
| Type 0 (Conserved) | 602 ± 27 | 4 ± 0.2% | 6 ± 0.33% |
| Type 1 | 3,020 ± 69 | 20 ± 0.5% | 29 ± 0.7% |
| Gain in mouse | 1,600 ± 33 | 11 ± 0.2% | 15 ± 0.3% |
| Gain in human | 1,420 ± 36 | 10 ± 0.2% | 15 ± 0.3% |
| Type 2 | 5,815 ± 249 | 39 ± 1.7% | 55 ± 2.3% |
| Expansion in mouse | 1,095 ± 36 | 7 ± 0.2% | 10 ± 0.3% |
| Expansion in human | 1,110 ± 47 | 7 ± 0.3% | 11 ± 0.4% |
| Expansion in both | 3,610 ± 166 | 24 ± 1.1% | 34 ± 1.6% |
| Type 3 (Cell type switch) | 1,062 ± 22 | 7 ± 0.1% | 10 ± 0.3% |
| Not expressed | 4,437 ± 209 | 30 ± 1.4% |  |

Numbers of genes and percentages are mean (±SEM) from using a median expression cutoff minus 0 to 2 standard deviations (0.25 increments) and a median percent cutoff plus 0 to 2 standard deviations (0.25 increments).

**Table S8.**
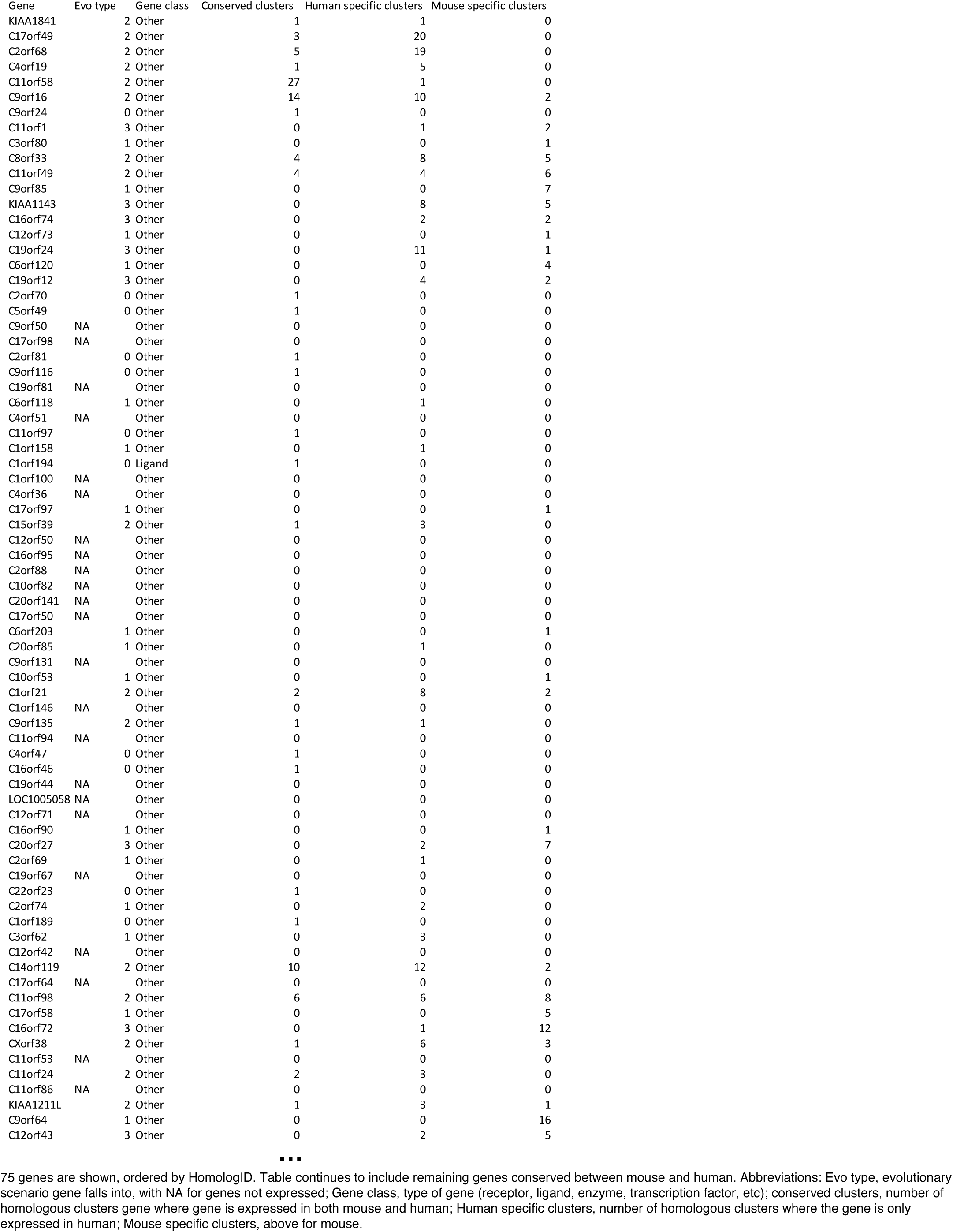
Evolutionary and functional classes of each gene.

## Notes

#### Summary of Updates

The revised manuscript includes 16 more quantified single molecule fluorescence in situ hybridization (smFISH) experiments of human lungs (Figs. 2, 4c, 6c-e, 7f,g, S4, S5), beyond the 10 such experiments in the original. This confirmed and localized all of the new human lung molecular cell types we discovered in single cell RNA sequencing (the only new molecular type not examined is the OLR1+ classical monocyte, presumably a new circulating cell type); the added experiments also localized and quantified many of the known human lung cell types, subtypes and states. We paired the smFISH experiments with textbook-quality micrographs documenting the normal classical histology of the human lung tissue in the subjects we profiled (Fig. S1). Table S2 summarizes all 58 human lung molecular types and their locations, which are diagrammed in Fig. 2b. The revision also includes 5,400 additional human lung cells profiled by SmartSeq2 (SS2), more than doubling our SS2 dataset to over 9,400 cells and our total number of profiled cells to over 75,000. Our SS2 dataset now includes 13 additional cell types we previously captured only with 10x Chromium, all of which are concordant with the previous 10x datasets while providing deeper expression coverage. We have publicly release all of our computer code and data (gene count/UMI tables, scanpy objects, and Seurat objects), and we are preparing for release of the FASTQ sequence files. We have also developed and opened an online portal of our atlas (https://hlca.ds.czbiohub.org/). Finally, and perhaps most significantly during the coronavirus pandemic, we curated from the literature a list of all human genes known to encode a receptor for a human virus including receptors for Covid-19, SARS (Severe Acute Respiratory Syndrome), and MERS (Middle East Respiratory Syndrome) coronaviruses and other respiratory viruses. We provide the lung cell expression profiles for all of these receptors (Figs. 6b, S8a), predicting lung cell tropisms of the viruses and where they initiate infection (p. 17). Our analysis predicts that Covid-19 (as well as SARS and MERS) infects alveolar cells, most prominently alveolar AT2 cells (Fig. S8b). Indeed, this data along with other emerging data and patient clinical data suggests that virus-mediated AT2 cell dysfunction or destruction is the key pathogenic event in Covid-19.

https://github.com/krasnowlab/HLCA

https://www.synapse.org/#!Synapse:syn21041850/wiki/600865

## References

1. Schwann, T. Mikroskopische Untersuchungen über die Uebereinstimmung in der Struktur und dem Wachsthum der Thiere und Pflanzen. (1839).

2. Weigert, C. Ueber die pathologischen Gerinnungsvorgänge. Archiv f. pathol. Anat. 79, 87–123 (1880).

3. Porter, K. R. et al. A study of tissue culture cells by electron microscopy: methods and preliminary observations. Journal of Experimental Medicine 81, 233–246 (1945).

4. Gehr, P. et al. The normal human lung: ultrastructure and morphometric estimation of diffusion capacity. Respiration Physiology 32, 121–140 (1978).

5. Ross, M. H. & Pawlina, W. Histology. (Lippincott Williams & Wilkins, 2006).

6. Bogoch, S. et al. Separation of cerebroproteins of human brain. Nature 204, 73–75 (1964).

7. King, R. J. et al. Isolation of apoproteins from canine surface active material. Am. J. Physiol. 224, 788–795 (1973).

8. Balis, J. U. et al. Distribution and subcellular localization of surfactant-associated glycoproteins in human lung. Lab. Invest. 52, 657–669 (1985).

9. Hermans, C. & Bernard, A. Lung epithelium-specific proteins: characteristics and potential applications as markers. Am. J. Respir. Crit. Care Med. 159, 646–678 (1999).

10. Schena, M. et al. Quantitative monitoring of gene expression patterns with a complementary DNA microarray. Science 270, 467–470 (1995).

11. Wang, Z. et al. RNA-Seq: a revolutionary tool for transcriptomics. Nat. Rev. Genet. 10, 57–63 (2009).

12. Tang, F. et al. mRNA-Seq whole-transcriptome analysis of a single cell. Nat. Methods 6, 377–382 (2009).

13. Wu, A. R. et al. Quantitative assessment of single-cell RNA-sequencing methods. Nat. Methods 11, 41–46 (2014).

14. Gawad, C. et al. Single-cell genome sequencing: current state of the science. Nat. Rev. Genet. 17, 175–188 (2016).

15. Treutlein, B. et al. Reconstructing lineage hierarchies of the distal lung epithelium using single-cell RNA-seq. Nature 509, 371–375 (2014).

16. Montoro, D. T. et al. A revised airway epithelial hierarchy includes CFTR-expressing ionocytes. Nature 560, 319–324 (2018).

17. Plasschaert, L. W. et al. A single-cell atlas of the airway epithelium reveals the CFTR-rich pulmonary ionocyte. Nature 560, 377–381 (2018).

18. Han, X. et al. Mapping the Mouse Cell Atlas by Microwell-Seq. Cell 173, 1307 (2018).

19. Tabula Muris Consortium et al. Single-cell transcriptomics of 20 mouse organs creates a Tabula Muris. Nature 562, 367–372 (2018).

20. Macosko, E. Z. et al. Highly Parallel Genome-wide Expression Profiling of Individual Cells Using Nanoliter Droplets. Cell 161, 1202–1214 (2015).

21. Cao, J. et al. The single-cell transcriptional landscape of mammalian organogenesis. Nature 566, 496–502 (2019).

22. Pijuan-Sala, B. et al. A single-cell molecular map of mouse gastrulation and early organogenesis. Nature 566, 490–495 (2019).

23. Zeisel, A. et al. Molecular Architecture of the Mouse Nervous System. Cell 174, 999– 1014.e22 (2018).

24. Saunders, A. et al. Molecular Diversity and Specializations among the Cells of the Adult Mouse Brain. Cell 174, 1015–1030.e16 (2018).

25. Vento-Tormo, R. et al. Single-cell reconstruction of the early maternal-fetal interface in humans. Nature 563, 347–353 (2018).

26. Enge, M. et al. Single-Cell Analysis of Human Pancreas Reveals Transcriptional Signatures of Aging and Somatic Mutation Patterns. Cell 171, 321–330.e14 (2017).

27. Angelidis, I. et al. An atlas of the aging lung mapped by single cell transcriptomics and deep tissue proteomics. Nature Communications 2017 8 10, 1–17 (2019).

28. Xu, J. et al. Deaths: final data for 2016. (2018).

29. World Health Organization. World health statistics 2019: monitoring health for the SDGs, sustainable development goals. (2019).

30. Franks, T. J. et al. Resident Cellular Components of the Human Lung. Proceedings of the American Thoracic Society 5, 763–766 (2012).

31. Picelli, S. et al. Full-length RNA-seq from single cells using Smart-seq2. Nature Protocols 2014 9:19, 171–181 (2014).

32. Blondel, V. D. et al. Fast unfolding of communities in large networks. J. Stat. Mech. 2008, P10008 (2008).

33. Braga, F. A. V. et al. A cellular census of human lungs identifies novel cell states in health and in asthma. Nature Medicine 2018 24:825, 1153–1163 (2019).

34. Howitt, M. R. et al. Tuft cells, taste-chemosensory cells, orchestrate parasite type 2 immunity in the gut. Science 351, 1329–1333 (2016).

35. Rock, J. R. et al. Notch-dependent differentiation of adult airway basal stem cells. Cell Stem Cell 8, 639–648 (2011).

36. Garcia, S. R. et al. Single-cell RNA sequencing reveals novel cell differentiation dynamics during human airway epithelium regeneration. bioRxiv 140, 451807 (2018).

37. Watson, J. K. et al. Clonal Dynamics Reveal Two Distinct Populations of Basal Cells in Slow-Turnover Airway Epithelium. Cell Reports 12, 90–101 (2015).

38. Nabhan, A. N. et al. Single-cell Wnt signaling niches maintain stemness of alveolar type 2 cells. Science 359, 1118–1123 (2018).

39. Zacharias, W. J. et al. Regeneration of the lung alveolus by an evolutionarily conserved epithelial progenitor. Nature 555, 251–255 (2018).

40. Stan, R. V. et al. The diaphragms of fenestrated endothelia: gatekeepers of vascular permeability and blood composition. Developmental Cell 23, 1203–1218 (2012).

41. Tan, S. Y. S. & Krasnow, M. A. Developmental origin of lung macrophage diversity. Development 143, 1318–1327 (2016).

42. van den Brink, S. C. et al. Single-cell sequencing reveals dissociation-induced gene expression in tissue subpopulations. Nat. Methods 14, 935–936 (2017).

43. Zheng, G. X. Y. et al. Massively parallel digital transcriptional profiling of single cells. Nature Communications 2017 8 8, 14049 (2017).

44. Shiow, L. R. et al. CD69 acts downstream of interferon-α/β to inhibit S1P 1 and lymphocyte egress from lymphoid organs. Nature 440, 540–544 (2006).

45. Mackay, L. K. et al. Hobit and Blimp1 instruct a universal transcriptional program of tissue residency in lymphocytes. Science 352, 459–463 (2016).

46. Moffitt, J. R. & Zhuang, X. RNA Imaging with Multiplexed Error-Robust Fluorescence In Situ Hybridization (MERFISH). Methods in Enzymology 572, 1–49 (2016).

47. Wang, X. et al. Three-dimensional intact-tissue sequencing of single-cell transcriptional states. Science 361, eaat5691 (2018).

48. Eng, C.-H. L. et al. Transcriptome-scale super-resolved imaging in tissues by RNA seqFISH. Nature 568, 235–239 (2019).

49. Online Mendelian Inheritance in Man, OMIM®. McKusick-Nathans Institute of Genetic Medicine, Johns Hopkins University (Baltimore, MD), (2018). https://omim.org/

50. Buniello, A. et al. The NHGRI-EBI GWAS Catalog of published genome-wide association studies, targeted arrays and summary statistics 2019. Nucleic Acids Res 47, D1005–D1012 (2019).

51. Huang, C. et al. Clinical features of patients infected with 2019 novel coronavirus in Wuhan, China. The Lancet 395, 497–506 (2020).

52. Limjunyawong, N. et al. Measurement of the pressure-volume curve in mouse lungs. JoVE (Journal of Visualized Experiments) e52376 (2015).

53. Seeley, R. R. et al. Essentials of anatomy and physiology. (2005).

54. Miller, A. J. et al. Basal stem cell fate specification is mediated by SMAD signaling in the developing human lung. bioRxiv 461103 (2018).

55. Lambrechts, D. et al. Phenotype molding of stromal cells in the lung tumor microenvironment. Nature Medicine 2018 24:824, 1277–1289 (2018).

56. Tabula Muris Consortium et al. A Single Cell Transcriptomic Atlas Characterizes Aging Tissues in the Mouse. bioRxiv 661728 (2019).

57. van Amerongen, R. et al. Developmental stage and time dictate the fate of wnt/β-catenin-responsive stem cells in the mammary gland. Cell Stem Cell 11, 387–400 (2012).

58. Greif, D. M. et al. Radial Construction of an Arterial Wall. Developmental Cell 23, 482– 493 (2012).

59. Muzumdar, M. D. et al. A global double-fluorescent Cre reporter mouse. Genesis 45, 593–605 (2007).

60. Madisen, L. et al. A robust and high-throughput Cre reporting and characterization system for the whole mouse brain. Nature Neuroscience 2009 13:113, 133–140 (2010).

61. Moraga, I. et al. Tuning Cytokine Receptor Signaling by Re-orienting Dimer Geometry with Surrogate Ligands. Cell 160, 1196–1208 (2015).

62. Desai, T. J. et al. Alveolar progenitor and stem cells in lung development, renewal and cancer. Nature 507, 190–194 (2014).

63. Butler, A. et al. Integrating single-cell transcriptomic data across different conditions, technologies, and species. Nature Biotechnology 2018 36:536, 411–420 (2018).

64. Finak, G. et al. MAST: a flexible statistical framework for assessing transcriptional changes and characterizing heterogeneity in single-cell RNA sequencing data. 16, 278 (2015).

65. Hu, H., et al. AnimalTFDB 3.0: a comprehensive resource for annotation and prediction of animal transcription factors. http://bioinfo.life.hust.edu.cn/AnimalTFDB/

